# Analysis of genetic dominance in the UK Biobank

**DOI:** 10.1101/2021.08.15.456387

**Authors:** Duncan S. Palmer, Wei Zhou, Liam Abbott, Nikolas Baya, Claire Churchhouse, Cotton Seed, Tim Poterba, Daniel King, Masahiro Kanai, Alex Bloemendal, Benjamin M. Neale

## Abstract

Classical statistical genetic theory defines dominance as a deviation from a purely additive effect. Dominance is well documented in model organisms and plant/animal breeding; outside of rare monogenic traits, however, evidence in humans is limited. We evaluated dominance effects in >1,000 phenotypes in the UK Biobank through GWAS, identifying 175 genome-wide significant loci (P < 4.7 × 10^−11^). Power to detect non-additive loci is low: we estimate a 20-30 fold increase in sample size is required to detect dominance loci to significance levels observed at additive loci. By deriving a new dominance form of LD-score regression, we found no evidence of a dominance contribution to phenotypic variance tagged by common variation genome-wide (median fraction 5.73 × 10^−4^). We introduce dominance fine-mapping to explore whether the more rapid decay of dominance linkage disequilibrium can be leveraged to find causal variants. These results provide the most comprehensive assessment of dominance trait variation in humans to date.

## Main text

There is a wealth of evidence suggesting that non-additive effects provide a non-negligible contribution to phenotypic variance in model organisms and animal breeding studies (*1–8*). However, these effects are not typically included in the genetic analysis of complex traits in humans. Even the simplest nonlinear contribution to effect on a given trait, Fishers dominance deviation (*9*), is generally omitted from genome-wide association studies (GWAS), on the grounds that the additive model captures most of the contribution of the locus even in the presence of a dominant architecture in all but the most extreme cases (*10*). Following Fisher 1918, we parametrise the dominance contribution at a locus orthogonally to the additive one, so that a non-zero contribution indicates any deviation from a purely additive or dosage pattern.

Empirically, outside of rare variants reported to cause recessive Mendelian disease phenotypes (*11,12*), evidence of non-additive variance explained across single nucleotide polymorphisms (SNPs) is weak. Extensions to the GREML model (*13*) to allow for estimation of dominance variance place the average contribution to broad sense heritability at around one fifth of that explained by the additive component, and the authors suggest a limited contribution to phenotypic variance. However, Chen *et al*. (*14*) found increased sample sizes in older twin pairs resulted in an increased contribution of dominance variation in twin-based estimation: an average of 0.25 in the 18 traits they analysed. A huge study drawing from twin studies across the last 60 years estimated that of the ~ 18,000 traits analysed, 69% were consistent with a simple additive genetic model (*15*).

Population biobanks of hundreds of thousands of individuals, including the UK Biobank offer the opportunity to explore questions of genetic architecture. In this work, we sought to investigate the role of dominance effects on phenotype. We examined non-additive variation in human disease using 71 truncated ICD-10 codes and 81 disease endpoints curated by FinnGen, as well as 267 quantitative traits, including measures for 31 biomarkers in 361,194 participants. As our approach requires a re-coding of genotypes which is a function of minor allele frequency (MAF), we restrict analysis to individuals in the UK Biobank with British or Irish ancestry (*16,17*). We estimated the contribution of dominance to phenotypes both at single loci through a series of 1,060 ‘dominance scans’, and tagged by common variation genome wide through an extension to LD-score regression. Our new method: dominance LD-score regression (d-ldsc) is scalable to thousands of phenotypes and can readily incorporate the genetic correlation and partitioned heritability extensions of LD-score regression (*18–20*).

### Testing for dominance effects at each variant

In this work, we wish to determine whether there is evidence for genetic effects beyond additivity at a locus contributing to variation in a collection of phenotypes. With this goal in mind, the sensible approach is to regress out any additive effect site by site, and test for evidence of any residual contribution to trait variation. Doing so amounts to testing for evidence of a dominance deviation (*9*) from additivity. This is not the same as re-coding genotypes according to the canonical biological dominance encoding [0,1,1], as illustrated in Fig. 1B. Notice that even with a MAF of 0.5, the additive model captures the majority of the variance explained by the locus (88.8%) when the true underlying architecture is [0,1,1]. Furthermore, if a dominance deviation exists, then the contribution to variance explained by the dominance deviation depends on the MAF. This can be seen by contrasting the first and second rows of Fig. 1. We use the terms non-additivity and dominance interchangeably to refer to the non-additive contribution to variance at a site which is uncorrelated to the additive model, as it encodes the dominance deviation.

**Figure 1:**
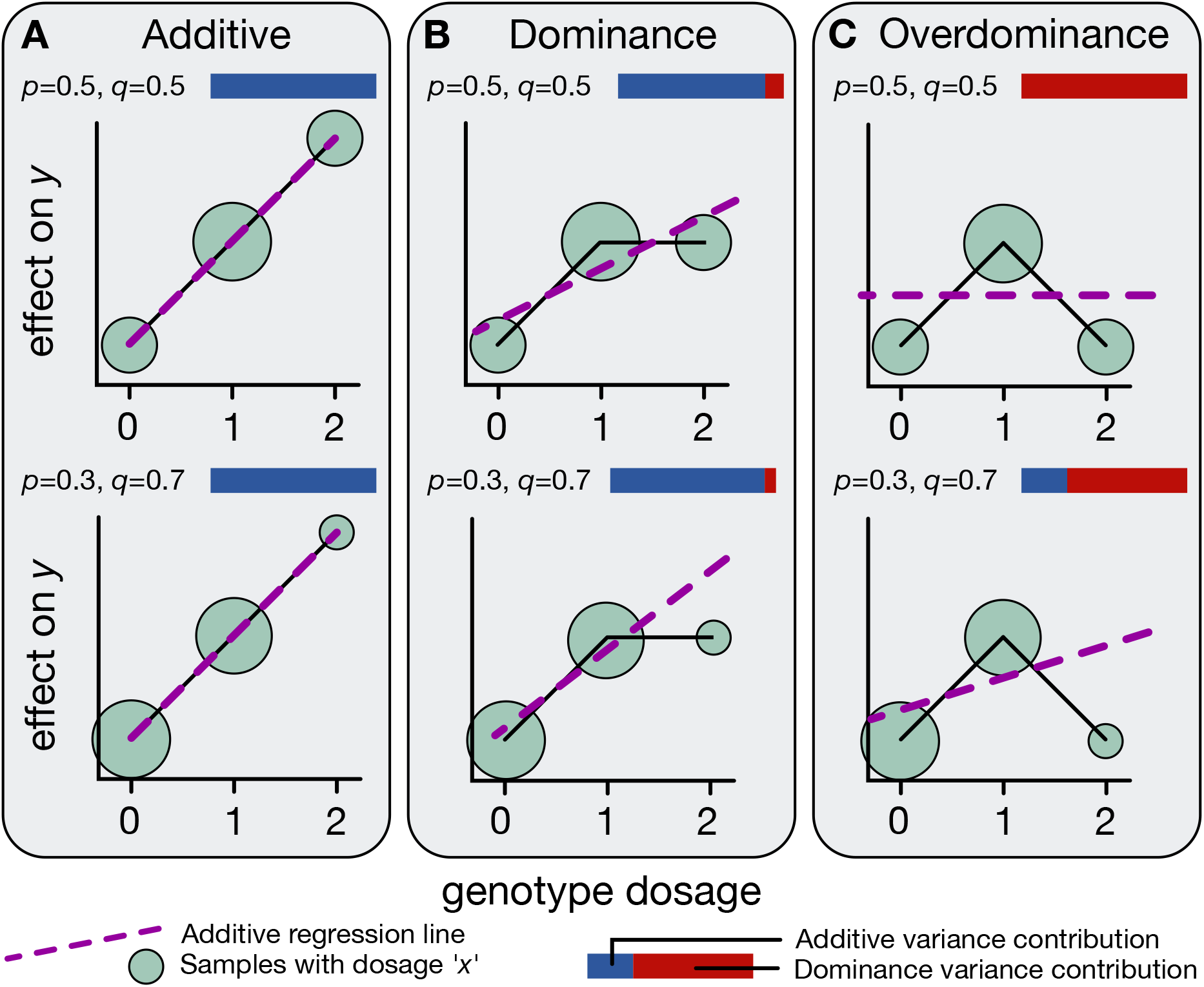
Examples of inheritance patterns at different MAFs. The effect sizes under a collection of genetic architectures is shown by the black line. The expected proportion of individuals with each of the three genotypes is shown by a green circle under the alternative allelic dosage (0, 1, 2); the area of the circle scales with the expected proportion of samples with that genotype. The extra variance captured by deviations from that additive fit (dashed purple line) is non-additive or dominance contribution to phenotypic variance. A. Purely additive genetic architecture at the SNP, no deviation of the truth (black line) from the additive fit (dashed purple line), so no extra variance is explained by non-additive effects, independent of MAF (*p* = 0.5; top row, p = 0.3; bottom row, variance contribution for both MAFs is entirely blue; representing 100% additive variance contribution at this site). B. Biological dominance architecture at a very common SNP (p = 0.5; top row) and at a common SNP (p = 0.3; bottom row). Despite being the canonical ‘dominance’ architecture, note that additivity explains a large portion of the variance (the dashed purple line is not horizontal and the variance contribution of additive effects is high: see the relative length of the blue bar to the red bar), but there is an appreciable amount of variation which cannot be explained by a purely additive model. This contribution decreases as MAF decreases from 0.5. I.e. the allele frequency of the SNP matters. Due to the rarity of the homozygous alternate genotype (shown by a smaller green circle beneath the black line at ‘2’), the additive model explains a larger portion of the total variance at the SNP. The variance contribution of the recessive contribution is equivalent to the biological dominance encoding of the other allele, and amounts to swapping the alternative allele. C. Overdominance at a very common SNP (p = 0.5; top row) and at a common SNP (p = 0.3; bottom row). When p = 0.5 (top row), half the sample is expected to be heterozygous, which completely balances the homozygous individuals so that the additive model explains none of the variance (the dashed purple line is horizontal, the variance contribution is entirely red) for this genetic architecture at a SNP with p = 0.5. However, overdominance architecture with any other MAF will contain an additive contribution, for example the bottom row, where p = 0.3.

Using a re-coding of the genotypes to capture the dominance deviation, referred to hereafter as the dominance encoding (see Theory: parametrisation of non-additivity at a locus, for our reasoning), we then assess within locus non-additive effects using a series of dominance GWAS scans and estimate the relative contribution of additive and non-additive genetic variation to phenotypic variance at top loci.

### Theory: Parametrisation of non-additivity at a locus

The dominance encoding amounts to mapping the genotypes [0,1,2] to 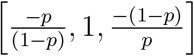 for each SNP, where *p* is the MAF of the SNP (Supplementary note: LD-score regression with dominance: Eqns (33)-(45), (*13*)). The model incorporating both an additive effect and dominance deviation site-by-site is then:

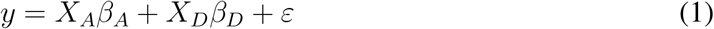

where *y* is a phenotype vector across the samples, *X_A_* and *X_D_* are matrices of the standardised additive and dominance encoding of the genotypes, *β_A_* and *β_D_* are column vectors of causal effect sizes, and *ε* is a vector of contributions to the phenotype through environmental contributions and higher order genetic contributions. Re-coding the sample-by-variant genotype matrix in this way means that the additive contribution to the phenotype (*X_A_β_A_*) is unchanged following the addition of dominance effects (*X_D_β_D_*). Not only is each column of our dominance encoded genotype matrix uncorrelated with its additive counterpart assuming Hardy-Weinberg Equilibrium (HWE), but also with every other additively encoded SNP (Supplementary note: LD-score regression with dominance: Eqns (72)-(75)).

In light of this orthogonality between additive and dominance encodings, we extend the standard GWAS marginal regression framework to perform simple marginal regression of phenotype on both *X_j,A_* and *X_j,D_* for each variant *j*, yielding GWAS coefficients 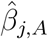 and 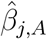 respectively. Using Fishers dominance deviation parametrisation serves several of purposes: it enables us to be agnostic to the marginal diploid genotype-phenotype pattern, leads to the same estimator for marginal additive effects 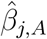 as in the usual additive-only model, and provides a straightforward partitioning of phenotypic variance explained by genetics.

Fig. 1 displays examples of inheritance patterns at different MAFs. In each panel, the nonadditive variance captured by deviations from the best linear fit (shown by the dashed purple line) is the contribution to phenotypic variance that our method will estimate. The largest contributions to non-additive variation at a site will occur if dominance and overdominance inheritance patterns are widespread at high MAFs. See Fig. S2 for examples of the additive and dominance encoding at varying MAF and effect size, followed by their combined contribution to the phenotype, to help guide intuition for how the additive and dominance effects combine to represent any inheritance pattern at a SNP.

It is relevant that linkage disequilibrium (LD) behaves differently under the dominance encoding: in particular, dominance LD decays much more rapidly than additive LD as one moves away from an index SNP (the former is in fact the square of the latter: Fig. S1B, Supplementary note: LD-score regression with dominance: Eqns (72)-(75), (*13, 21*)). One may ask whether this behavior can be exploited in fine-mapping a locus; we explore this idea by developing a modification of the SuSiE method (*22*) to ‘dominance fine-map’ a collection of leading dominance loci.

### Application: Association studies in the UK Biobank

Following careful curation of the phenotypic and genotypic data (361,194 samples, 13.7 million variants, 1,060 well-covered phenotypes (Materials and methods: phenotype curation, sample and genotype curation, Fig. S3-4)), we ran additive and dominance GWAS using *age, age*^2^, *sex, age* × *sex, age*^2^ × *sex*, and the first 20 principal components as covariates (Materials and methods: evaluation of additive and dominance marginal effect sizes). To avoid potential bias arising due to ancestry differences, we considered the individuals with British and Irish ancestry (*23*). For continuous traits, we analysed the inverse-rank normal transformation (IRNT) of the raw phenotype. This was to avoid false positive dominance associations which can arise due to heteroskedasticity in the noise term unaccounted for in Eqn. (1), (Materials and methods: checking for artefactual dominance: heteroskedastic noise, Box: the impact of scale). We note that this does not remove all potential sources of heteroskedasticity, but works well in removing spurious inflation in dominance test statistics empirically. Restricting to SNPs with MAF > 0.05, we found 175 genome-wide significant (using a conservative Bonferroni cutoff: 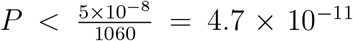) loci: 128 in continuous traits, 47 in case-control traits. Loci were defined by considering 500kb windows around significant SNPs; merging loci where windows overlapped. To check for potential sequencing artefacts, we examined the distribution of imputation quality score (*24*), P-HWE, inbreeding *F*-statistic, and cluster confidence (*25*) at the closest genotyped variant across MAF bins (Fig. S5-7). Dominance loci were more confidently called and have greater cluster confidence than random loci. As expected, dominance hits were enriched for lower P-HWE. To guard against incorrect calls, we excluded variants with P-HWE < 10^−6^. A summary of the dominance and additive GWAS results are shown in Fig. 2 and Fig. S8. We verified empirically and via simulation that these results were not due to deviations from HWE (Materials and methods: checking for artefactual dominance: deviations from Hardy-Weinberg equilibrium, Fig. S9-S13). At lead SNPs in loci exhibiting a genomewide significant dominance association, estimates of the underlying genetic architecture were enriched for monotonic functions of dosage: of the 175 genome-wide significant dominance loci, we observed monotonic functions of dosage at the lead dominance SNP in 132 (75.4%).

**Figure 2:**
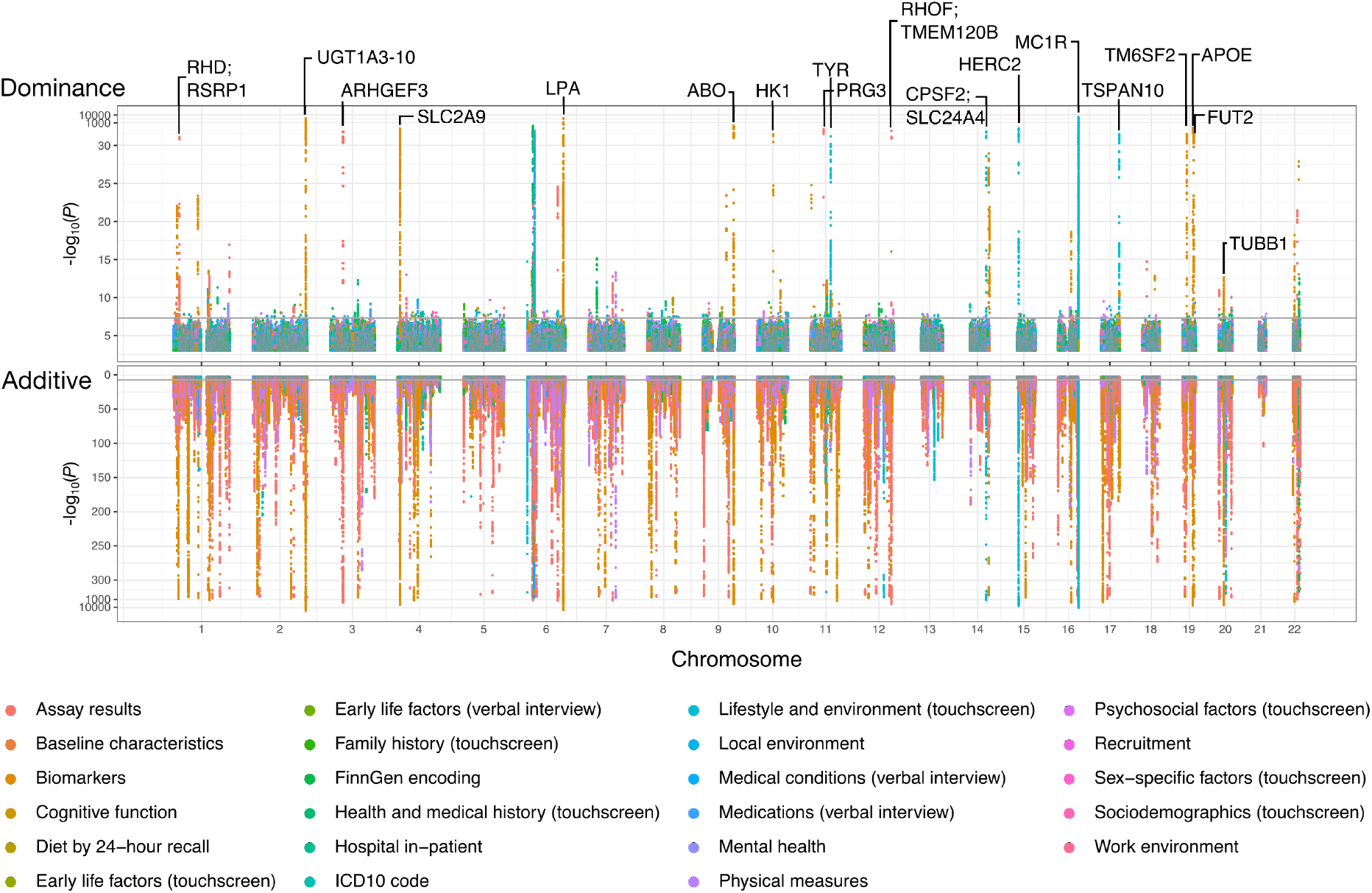
Aggregate Manhattan plots for additive and non-additive marginal effect sizes. We determine the most extreme t-statistic for marginal effect sizes across all well covered traits at each site with MAF > 0.05, and plot the associated – log_10_(*P*) value for the non-additive (top) and reflect the additive – log_10_ (*P*) value (bottom). Traits are grouped into broad categories as defined in the UK Biobank data showcase and coloured according to the legend. To aid presentation, in panel A (B respectively) the y-axis is on the – log_10_ scale up to 30 (300 respectively), after which it switches to a – log_10_(– log_10_(*P*)) scale. Plots on the – log^10^(*P*) scale are shown in Fig. S7.

We replicated known non-additive effects, including rs1805007 in *MC1R* for hair colour (*26,27*) (*P* < 1 × 10^−5000^ for red hair, Fig. S14), an intronic variant of *HERC2* which functions as an enhancer regulating *OCA2* expression for hair and skin colour (*28,29*) (rs12913832, P = 2.87 × 10^−56^ and *P* = 5.33 × 10^−191^ for blonde hair and skin colour respectively, Fig. S15-16), and a non-additive signal tagging *ILDR1* for hearing difficulty which is stronger than the additive signal at the locus (additive *P* = 4.26 × 10^−8^, dominance *P* = 5.79 × 10^−13^, Fig. S17). *ILDR1* is a known Mendelian hearing loss gene (30–32). The stronger dominance signal reflects the high MAF and putative overdominant contribution to the phenotype. We also observed a genomewide significant non-additive association with red blood cell distribution width (rs67002563, *P* = 2.99 × 10^−11^, Fig. S18), in tight linkage-disequilibrium with *ITPA.* Notably, this locus did not reach genome-wide significance under additivity (*P* = 0.000152, Fig. S18). *ITPA* has been implicated in red blood cell disorders through an autosomal recessive mode of inheritance (*33–36*). A strong non-additive association was also observed for the distribution width of platelets at an intronic eQTL (rs1354034, *P* = 5.80 × 10^−90^, Fig. S19) associated with *ARHGEF3* expression in platelets (*37*). *ARHGEF3* displays a regulatory role in myeloid differentiation in zebrafish (*38*). The highly significant additive association for platelet count and volume at this locus was completely ablated for distribution width (*P* = 0.825, Fig. S19). Throughout, we ran linear regression to identify associations to enable cost effective examination of thousands of traits. In the presence of small effect sizes and MAF > 0.05, we expect P-values for linear and logistic regression to be extremely similar (Supplementary note: similarity of logistic and linear regression with small effect sizes). We assessed whether this choice of model materially affected the results by running logistic regression at all dominance loci which were genomewide significant under a standard linear model following the dominance encoding. P-values were highly correlated at dominance loci (mean *r* = 0.993), with the median difference in log(*P*) of 0.02 across all SNPs within 500kb of the lead SNP.

### Application: The relative variance contribution of additive and dominance effects at top loci

To probe the relative variance explained by the additive and non-additive contributions, we first examined their relative contributions across the top loci. We annotated variants with MAF > 0.05 in the full dataset using ANNOVAR (*39*), and took the top additive and dominance associations across unique cytobands for each phenotype. Effect sizes were rescaled such that the squared effect size represents an estimate of the variance explained by the locus, and summed across the top five additive and dominance associations respectively. We then plot the relative contribution to variance explained by the additive and uncorrelated non-additive components in Fig. 3A. The median ratio of the two variance components is 20.9. If we further enforce that the additive and dominance associations must have a signal of association *P* < 1 × 10^−6^, the median ratio of the variance components is 28.0; displayed in Fig. 3B. Therefore, assuming the same distribution of genetic architectures across MAF and effect size, we expect to require millions of samples (~ 7,500,000) to be powered to detect non-additive effects to the same level of significance as those currently reported for additive effects (*40*). Furthermore, this is a best-case scenario for our ability to detect non-additive effects for any locus-by-locus: given that there is much less dominance variance for any particular locus, the expected amount of noise variance in the marginal dominance effect size contribution is greater than the additive noise variance.

**Figure 3:**
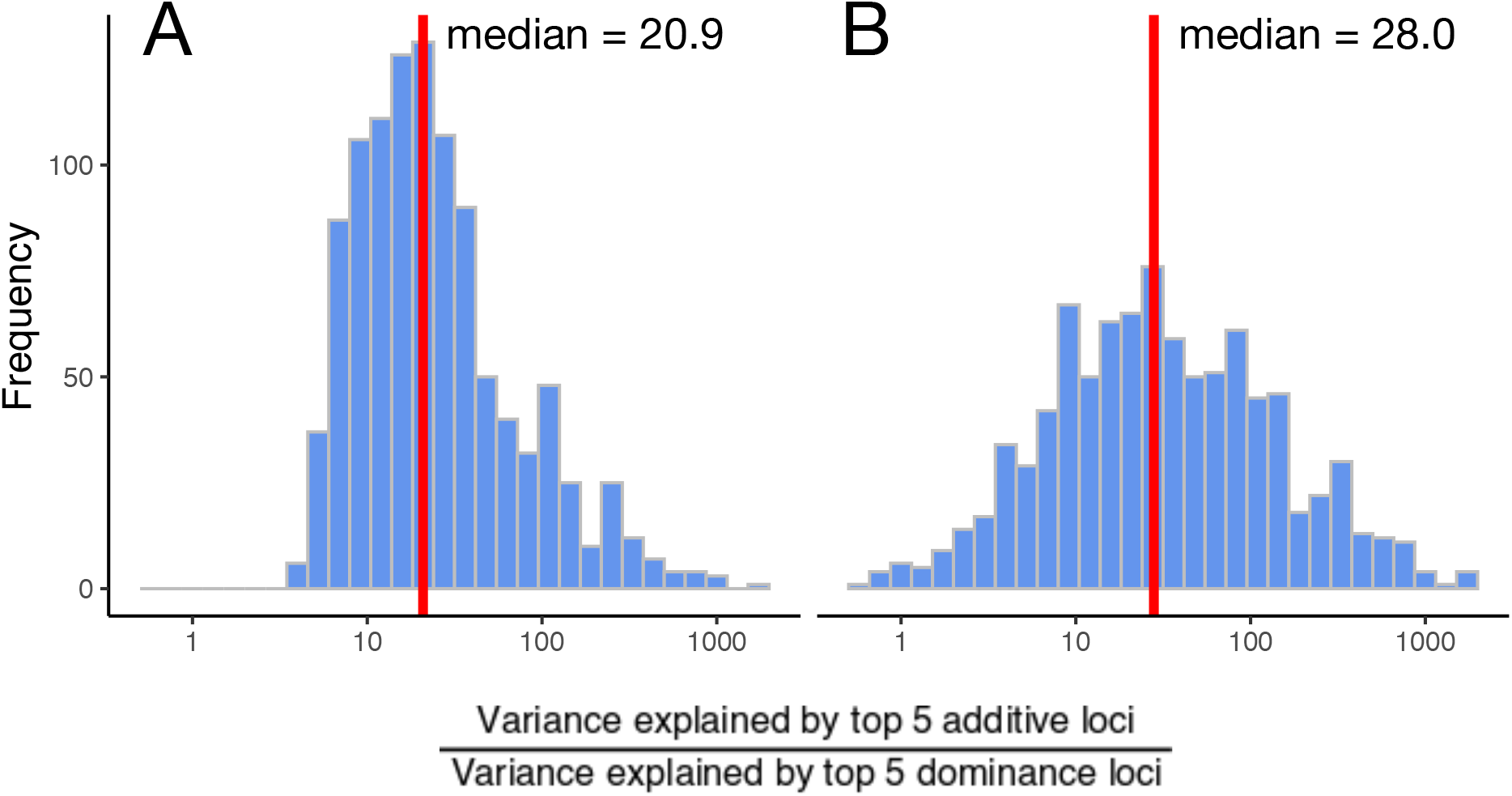
Relative power to detect dominance associations. We show the distribution across phenotypes of the ratio of the variance explained by the top 5 additive loci to the variance explained by the top 5 dominance loci. The x-axis is on the log scale. In A, we place no P-value restriction for inclusion of the most significant association in the cytoband. In B, we enforce that each association must have *P* < 1 × 10^−6^.

### Theory: Fine-mapping dominance loci

By combining the observations that not all dominance GWAS are devoid of signal (Fig. 2) and that the dominance flavour of LD decays at the square of additive LD (Fig. 1B), we sought to determine whether we could fine-map these non-additive signals. By doing so, we looked to distinguish between distinct causal variants within a locus when there is a differential contribution of dominance signals at these causal sites. We used a straightforward method to perform dominance fine-mapping that can be naturally applied due to the partitioning of effects at each SNP into uncorrelated additive and dominance effects: we can simply take the dominance LD matrix (an entry-wise squared additive LD matrix) and marginal dominance effect size esti-mates as input into existing fine-mapping software that act on summary statistics to fine-map dominance effects (Materials and methods: fine-mapping).

### Application: Fine-mapping dominance loci

We used SuSiE (*22*) to fine-map the dominance signal at nominally significant (*P* < 5 × 10^−8^) dominance loci. We then fine-mapped the additive effects at these loci and compared the results. A summary is displayed in Fig. 4. While acknowledging the likelihood of ‘winner’s curse’ in these results (we restricted attention to significant dominance loci), we observed differences in fine-mapping. Of the variant-phenotype pairs with additive posterior inclusion probability (PIP) > 0.2 and more evidence of causal signal under the additive encoding (bottom right, *n* = 710), 22.6% are distinct additive signals without a dominance component lying in the same gene as a confidently fine-mapped dominance association. Of particular interest are those variants lying above the *y* = *x* line (top left, n = 330 with dominance PIP > 0.2). These are putatively causal variants that were more confidently fine-mapped via the dominance association. Table S1 summarises the collection of exonic variants that are more confidently dominance fine-mapped. The non-additive signal in the *ITPA* locus associated with red blood cell distribution width was fine-mapped to rs1127354, a non-synonymous single nucleotide variant which is known to predict drug induced anaemia among chronic HCV patients (*41–45*). The genome-wide significant non-additive association with hearing difficulty/problems was fine-mapped to rs2877561, a synonymous change in *ILDR1* associated with age-related hearing impairment (*46*) but which did not reach genome-wide significance in that study. We note that rs2877561 is an eQTL and sQTL for a large number of genes and tissues in GTEx (*47*).

**Figure 4:**
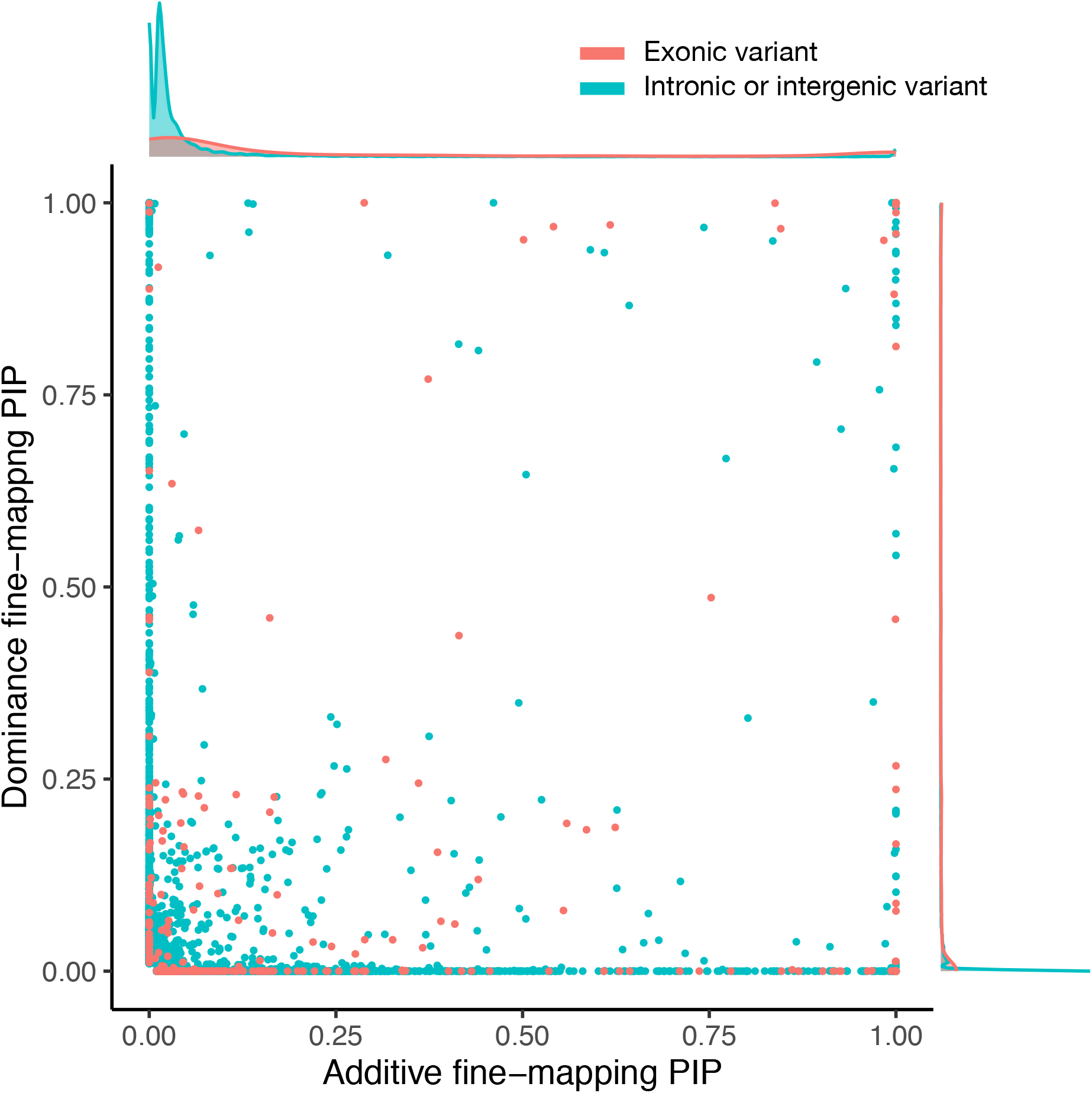
Fine-mapping dominance loci using SuSiE. We took the collection of genome wide significant dominance loci tagged by SNPs with MAF > 0.05, and fine-mapped using SuSiE ((*22*), Materials and methods: fine-mapping). This amounted to passing dominance effect sizes and within sample dominance LD. We then plot the additive and dominance posterior inclusion probabilities against each other for all dominance loci across all phenotypes. Red points are in the exome, teal points are intronic or intergenic.

### Genome-wide dominance – a matter of LD genome-wide

Another question that presents itself following our dominance scans is what proportion of phenotypic variance can be explained by dominance effects? Some existing papers have suggested via theory and empirically that this dominance heritability hD is likely small in human complex traits (*13*). More recently, methods used to estimate the dominance contribution (which rely on the same partitioning of variance that we apply here) were extended and applied to 70 complex traits in the UK Biobank (*48*). Others have relaxed the assumptions of the model and applied it to a selection of 50 quantitative traits in the UK Biobank (*49*). These works find zero or marginal evidence of non-additive effects contributing meaningfully to phenotypic variance genome-wide.

By extending the LD-score software, we are able to estimate the dominance contribution to the variance of traits extremely rapidly. Following a dominance GWAS and generation of dominance LD-scores (in the sample or an ancestrally matched reference panel, Materials and methods: evaluation of additive and dominance heritability), dominance SNP heritability estimates can be obtained as quickly as additive SNP heritability estimates (*50*). This efficiency allowed us to estimate the non-additive variance contribution to all 1,060 curated phenotypes in the UK Biobank at low time and economic cost.

### Theory: Dominance LD-score regression

To estimate the common variant contribution of dominance effects to phenotypic variation hD, we have constructed a dominance LD-score regression model, which we call d-ldsc (see Materials and methods: LD-score dominance extension for a summary and Supplementary note: LD-score regression with dominance for a detailed derivation). Briefly, the dominance encoding of genotypes naturally leads to a new flavour of LD:

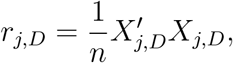

which can then be used in a new LD-score regression in which we regress the chi-squared statistics of our dominance GWAS on dominance LD-scores (Fig. S1B):

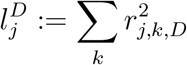

The key insight is that HWE assumption (Fig. S1A) leads to several simplifications, ultimately yielding a simple, decoupled analogue of the standard additive ldsc result (*18*):

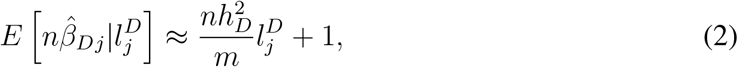

where *n* is the number of samples, *m* is the number of SNPs, 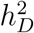 is the dominance heritability, and 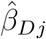 is the marginal effect of SNP *j* under the dominance encoding.

As in the original, we may estimate the dominance LD-scores using an ancestry matched reference panel. With access to dominance summary statistics and LD-scores, we can estimate the gradient of the linear model in Eq. (2) to obtain an estimate of 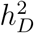. We use a block jackknife (*18*) to obtain standard errors on our estimates and determine their statistical significance. To ensure d-ldsc is an appropriately conditioned statistical model, we performed an extensive collection of simulation studies (Materials and methods: simulation studies). We considered two simulation scenarios: fully simulated genotypes and real genotypes (a random subset of the UK Biobank samples which we define as having British or Irish ancestry). We then simulated phenotype data according to the model in Eq. (1) under infinitesimal (all SNPs are causal) and spike and slab (10% of SNPs are causal) genetic architectures with varying sample sizes (10,000, 50,000, 100,000). We tested the accuracy of our estimator on both continuous and case-control phenotypes (under a liability threshold model) with varying levels of case-control ascertainment, and magnitudes of 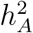 and 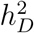 (0, 0.05, 0.2). Throughout, estimates are unbiased, and well calibrated under all simulation scenarios (Fig. S20-28, summarised in Tables S2-3).

#### Box: The impact of phenotype scale

As part of the phenotype curation process we compared and contrasted the impact of inverse-rank normal transformation (IRNT) of continuous phenotypes in our results as in the curation in the additive heritability browser (*51*). We find high concordance between evidence of association across the raw and IRNT flavours of continuous phenotypes (median Pearson correlation = 0.913), but a minority showed poorly calibrated QQ plots with an abundance of ‘hits’ and deflation at higher P-values. Similarly, a subset of ordinal phenotypes displayed likely spurious signals. An example of one of the most extreme of these was high light reticulocyte percentage on the raw scale (GC = 0.862, Fig. S29). Following IRNT, these artefacts were removed, with the resultant Manhattan plot displaying a few dominance ‘towers’, and well calibrated QQ plot at high P-values (GC = 1.02, Fig. S29). Due to the distribution of phenotypes containing this signature of likely spurious hits, we hypothesised that this error mode could be partially a result of heteroskedasticity of the noise term. For example, perhaps larger phenotypic values were measured with increased measurement error on the raw scale. To investigate, we performed a simple simulation study by simulating true effects under an infinitesimal additive model and adding a heteroskedastic noise term that increases in variance as the phenotype increases (Materials and methods: checking for artefactual dominance: heteroskedastic noise). We then performed the same simulation study with spike and slab architecture in the true effects. In each case, heteroskedastic noise did indeed give rise to spurious significant dominance signals and reduced GC which was then resolved through the IRNT (GC = 0.74 and 0.96 before and after IRNT for infinitesimal architecture, and GC = 0.78 and 0.99 for 1% causal spike and slab before and after IRNT respectively (Fig. S30-31)). As a consequence, we conservatively removed all raw and ordinal phenotypes from the rest of our analysis (but we make all raw and ordinal additive and dominance GWAS results publicly available (see Data and Materials Availability). Given the observation that a subset of variants in a collection of traits displayed artifactual dominance, we were concerned that similar effects could accumulate over small scale perturbations genome-wide, giving rise to significant but spurious dominance heritability for some traits. More generally, we were concerned that a simple change in scale could induce significant dominance heritability. To test the impact of phenotype scale or parametrisation both at the level of the locus and genome-wide, we applied a collection of non-linear transformations to standing height in the UK Biobank and re-ran GWAS and heritability analysis (Table S4-5). Second order and higher polynomial transformations induced non-additive effects as expected. Exponentiation of height did not, due to the small effect size polygenic nature of height and accuracy of the linear approximation under this regime (exp(*y*) ≈ 1 + *y* for *y* ≪ 1). Under all transformation scenarios, non-additive heritability was not induced, and we conclude that false positive nonadditive heritability estimation is unlikely. This is likely due to the transformations inducing heteroskedasticity which lead to both genome wide significant hits and deflation in dominance chi-squared statistics for non-significant variants, which effectively cancel each other out. Due to the better calibrated dominance QQ plots following the IRNT we also restrict to IRNT continuous phenotypes and remove ordinal phenotypes from our dominance heritability analyses.

### Application: Dominance heritability of traits in the UK Biobank

Applying additive and dominance LD-score regression after filtering (Fig. S32) to the 1,060 traits with dense phenotyping we found strong evidence of significant additive heritability as expected, but very little evidence of dominance heritability (Fig. 5, Table S6-7). This contrast was present in both continuous and case-control traits. For case-control phenotypes, we see increased evidence for variance explained by additive genetic architecture as case-count increases. This trend was not apparent for non-additive heritability tagged by common variation (MAF > 0.05); hD. These results support existing evidence for extremely modest additional contributions to phenotypic variance that a model incorporating non-additive effects provides over a purely additive model tagged by common variation (*13,48, 49*): across all 1,060 phenotypes we did not find evidence of a non-zero relative contribution of dominance heritability estimates to additive heritability estimates (gradient = 0.0024, *P* = 0.10; Fig 5A). This does not preclude the possibility that non-additivity is widespread throughout the genome, or that non-additive effects should be disregarded. Finally, power to detect deviations from additivity is weakest precisely where we expect the largest effects: rarer variation.

**Figure 5:**
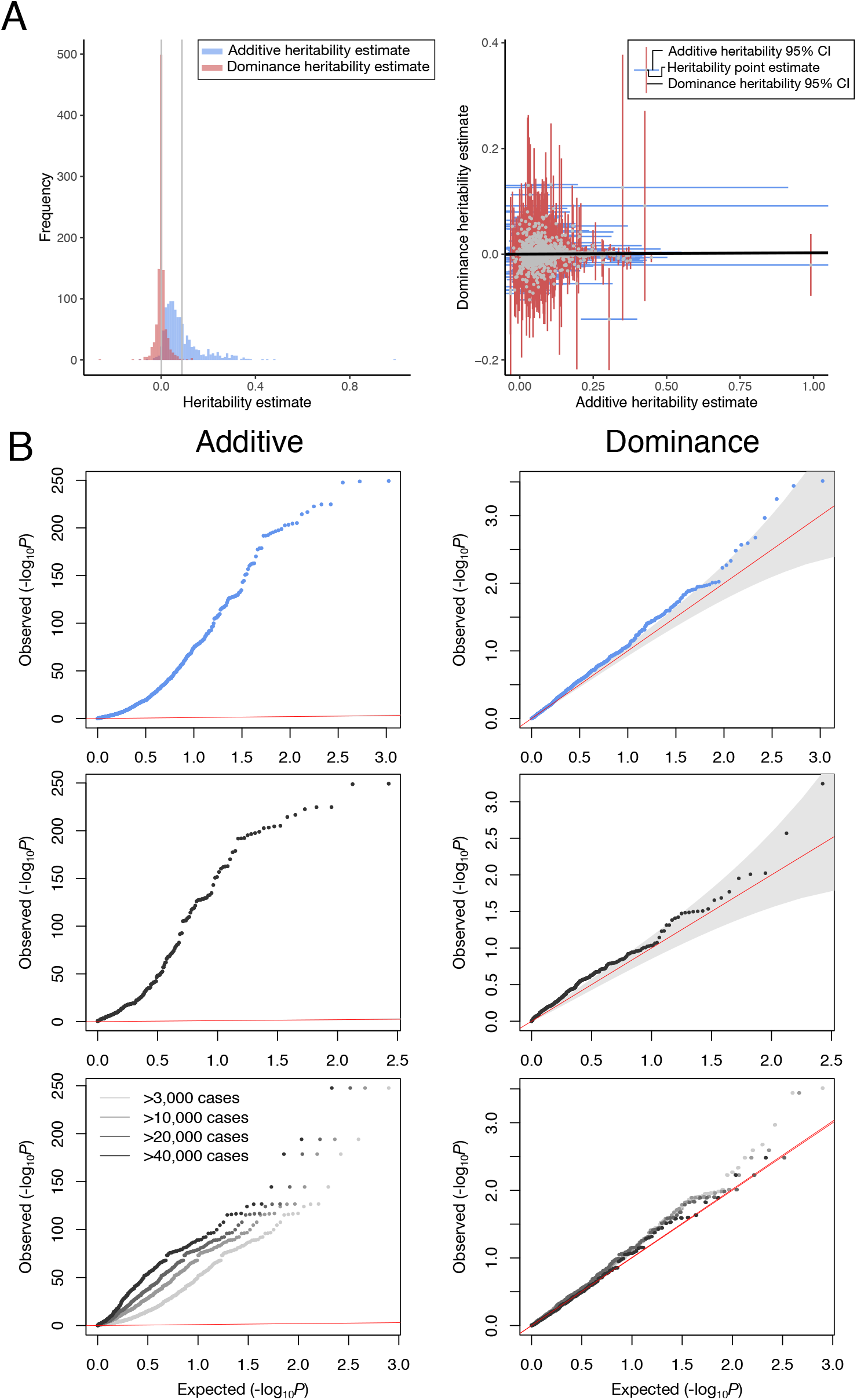
Summary of heritability analysis. In panel A, the first column displays histograms of LD-score based estimates of additive and dominance SNP heritability shown in blue and red respectively. Mean heritability estimates are shown by the grey lines. The second column displays the paired results for each phenotype, coloured according to the key. The York regression best fit line is displayed in black (intercept = 0.00026, gradient = 0.0024). In panel B, the first and second columns display QQ plots of observed against expected *P*-values evaluated using block jack-knife standard errors to test if 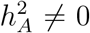 and 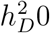 respectively. The first row includes all traits with over 50,000 data points if continuous or ordinal, and over 3,000 ‘cases’ if binary. In the second row, we restrict to the continuous and ordinal traits. Finally, in the third row, we restrict to the binary traits. Points in the final row are coloured according to the legend.

## Discussion

We have performed the largest and most comprehensive dominance scan and heritability analysis of phenotypes to date. We analysed 1,060 GWAS scans following careful phenotype curation, identifying 175 loci at genome-wide significance. These loci consisted of many well known associations in phenotypes with dominant and recessive patterns of inheritance (for example, hair colour). Qualitatively, we observed stronger non-additive effects in more tissuespecific phenotypes such as blood, hair colour traits, biomarkers, and in genes with strong additive associations such as APOE and HPE. For most traits, far more samples are likely to be required to capture evidence of dominance effects to genome-wide significance. Extrapolating from the lead five loci in each trait, we estimate that on the order of millions of samples will be required to obtain marginal dominance effect size estimates to a level of significance similar to that currently observed in additive GWAS.

Despite increased power over existing studies (*13, 14*), we still find limited evidence of a substantial dominance contribution to phenotypic variance. We hypothesise that the distribution of estimated dominance heritability being close to null remains a power issue, coupled with the low relative magnitude of the dominance variance to the additive variance enforced by the uncorrelated parametrisation of dominance (*52*). More generally, we find further evidence supporting the robustness of the linear model for human complex traits, reflecting that common variant effects are largely small perturbations of continuous latent biological processes aggregated by a mean-field approximation.

We introduce dominance fine-mapping to attempt more accurate estimation of causal variants in the presence of a dominance signal. Using the dominance effect sizes and the dominance LD matrix we are able to use existing software (*22*) to fine-map dominance loci. With the same strength of association, a dominance signal would fine-map more readily. However, gains in fine-mapping accuracy due to the far more rapid decay of dominance LD are generally outweighed by the increased effect size of the additive signal of association at the locus. It would be natural to explore using all information jointly to gain power to increase fine-mapping precision and distinguish distinct putatively causal variants in a locus.

We highlight two caveats and limitations. Throughout, we have assumed HWE in the determination of the dominance encoding and subsequent evaluation of dominance effect sizes and heritability. If this is not the case, additive effects will be pulled into the dominance encoding and manifest as non-zero dominance effect sizes. To counter this effect, we impose a stringent P-HWE cutoff for inclusion in our results. In doing so, we are removing a subset of SNPs of putatively large effect under selection that we are most well powered to detect. However, given we are focused on common variation, we expect that variants exhibiting this behavior are far more likely to be due to errors in genotyping rather than true effects of selection. Parametrisations exist that do not require an assumption of HWE (*53, 54*). We can also determine an orthogonal contribution within-sample empirically to avoid potentially capturing additive effects in the non-additive dominance contribution to phenotypic variation. Either of these extensions would help get around the issue of selection for or against a variant coinciding with departures from HWE leading to a spurious dominance association.

Secondly, in our dominance scans we assume the same cutoff for genome-wide significance as for additive GWAS. However, given the increased number of effectively independent sites in the genome implied by a more rapidly decaying LD structure, this assumption should be challenged. Under the approximation of a Poisson process for recombination, the effective number of independent SNPs is approximately double the number of independent additive markers. The benefit dominance LD provides for fine-mapping is a drawback for GWAS and detection of phenotypic variance explained: dominance LD tagging does not extend as far in the genome. This reduced tagging, in combination with the observation that we are most underpowered to detect associations at rare variants where we expect to see the strongest effect sizes, suggests that we should look to bottlenecked populations such as Finland and Iceland as their enrichment for rare variants and longer haplotypes may offer enhanced power to detect non-additive effects at a locus.

In this work, we estimated dominance heritability tagged by common SNPs genome-wide and did not consider partitions of the genome due to the general paucity of dominance variance explained. However, given a non-zero dominance variance, the entirety of the LD-score toolkit including partitioned heritability estimation (*20*) and genetic correlation estimation (*19*) can be applied to dominance effects. In addition to site-by-site dominance effects, LD-score regression is readily extendable to test gene-by-environment interactions; an extension to pairwise epistasis would be interesting but more challenging.

## Acknowledgements

We thank all members of the Neale lab for comments and suggestions, and thank the remaining members of the Hail team for their code suggestions, and continued improvements to the Hail codebase. We thank the participants and leadership of the UK Biobank: this work was carried out under UK Biobank application 31063. Data and materials availability Dominance summary statistics and heritability estimates are available for download from Amazon web services at s3://broad-ukb-sumstats-us-east-1/round2/dominance-tsvs/. We also performed sex-specific analyses, curating phenotypes according to the same pipeline (Materials and methods: phenotype curation), associated summary statistics files are also available at s3://broad-ukb-sumstats-us-east-1/round2/dominance-tsvs/. Code implementing the d-ldsc framework is available at https://github.com/astheeggeggs/d-ldsc. Code implementing the msprime simulation framework is available at https://github.com/astheeggeggs/msprime_sim.

## Author contributions

DSP, AB and BMN developed the theory, DSP implemented the d-ldsc pipeline, LA and DSP implemented the dominance GWAS pipelines, MK and DSP implemented the dominance fine-mapping pipeline. DSP and WZ performed all analyses and produced the figures. CS, TP, and DK implemented computational tools to enable the analyses. AB and BMN supervised the project. DSP and NB implemented and ran simulation studies. DSP and AB wrote the manuscript. BMN, MK, and CC provided valuable edits.

