## Supplementary materials for "Analysis of genetic dominance in the UK Biobank"

### Supplementary materials for
### Analysis of genetic dominance in the UK Biobank

Duncan S. Palmer[1,2,*], Wei Zhou[1,2], Liam Abbott[1,2,3],
Nikolas Baya[1,2,3], Claire Churchhouse[1,2,3], Cotton Seed[2,3], Tim Poterba[2,3],
Daniel King[2,3], Masahiro Kanai[1,2,3], Alex Bloemendal[1,2,3,†] and Benjamin M. Neale[1,2,3,†,*]

[1] Analytical and Translational Genetics Unit, Department of Medicine,
Massachusetts General Hospital, Boston, Massachusetts, USA.
[2]Stanley Center for Psychiatric Research,
Broad Institute of MIT and Harvard, Cambridge, Massachusetts, USA.
[3]Program in Medical and Population Genetics,
Broad Institute of MIT and Harvard, Cambridge, Massachusetts, USA.
[†]These authors contributed equally.

**This PDF file includes:**

Materials and Methods

Supplementary Note

Figs. S1 to S32

| Phenotype class | Initial count | $h^2$ browser filtering | Dominance filtering |
|---|---|---|---|
| Binary | 2,346 | 2,305 | 641 |
| Categorial | 1,192 | 1,039 | 152 |
| Continuous (IRNT) | 305 | 305 | 267 |
| Continuous (raw) | 305 | 0 | 0 |
| Ordinal | 271 | 260 | 0 |
| Total | 4,419 | 3,909 | 1,060 |

**Table** S1: Phenotype counts by phenotype category before and after phenotype curation filters.

### Materials and Methods

#### Phenotype curation

UK Biobank phenotypes were re-coded and curated. Briefly we use a modified version of the PHESANT software (*56*), enabling the generation of meaningful, coherent phenotypes for our downstream analysis. Curation of primary ICD10 codes truncated to two digits was carried out separately for computational efficiency. A summary of the PHESANT pipeline is displayed in Fig. S3. We also incorporated phenotypes curated by FinnGen, described elsewhere (*57*). The breakdown following this initial curation was:

Following initial phenotype curation using PHESANT, and incorporation of 633 ICD10 and 559 FinnGen phenotypes, we performed a further collection of filtering steps. These steps are described in detail in the methods portion of the Neale lab heritability browser (*51*). Briefly, we remove redundant FinnGen phenotypes (58 examples), and sex-specific phenotypes from the analysis. To do this we use a series of cutoffs. If over 97% of the total sample size for a phenotype comes from a single sex (51 such phenotypes), it is removed from our primary analysis (in which we analyse across the sexes). Further, for boolean phenotypes, if over 99.7% of the cases or controls comes from a single sex (enough to separate sex-specific phenotypes from strongly sex-biased biomedical phenotypes), it is removed from our primary analysis,

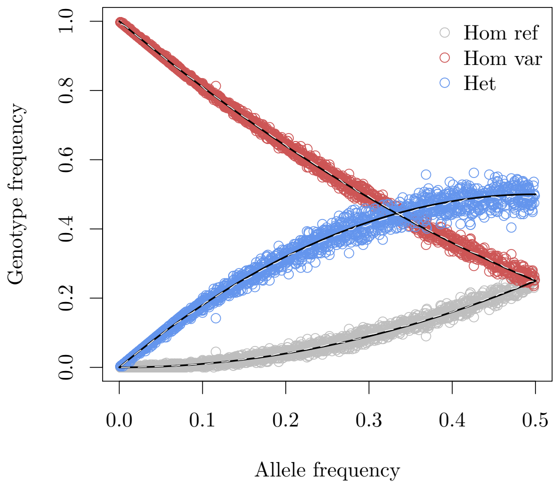


(A) Hardy-Weinberg equilibrium approximation.

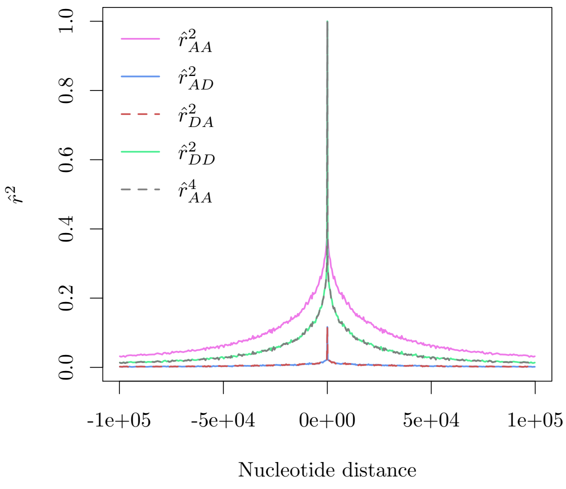


(B) Average value of $\widehat{r}^2$ moving away from a SNP.

**Figure** S1: HWE and decay of flavours of LD. Fig. S1A gives an indication of how prevalences of genotypes conform to Hardy-Weinberg equilibrium in European populations. Here, we plot the allele frequency against genotype frequencies for European chromosome 22 samples in the 1000 Genomes data-set (*55*). In black, we show the analytic Hardy-Weinberg proportions, and in dashed white we overlay the mean of European samples in the 1000 Genomes data (*55*). To calculate means, we split the data into allele frequency bins of width 0.005. In Fig. S1B we show the average values of our estimates of $r^2$ statistics as we move away from a SNP. To obtain the plot we take mean values within bins containing at least 5,000 estimates, with a minimum bin width of 400 base-pairs. We then plot out to a maximum distance of 100,000 nucleotides in each direction. Lines are coloured according to the legend. Where we expect similarity between estimates ($\widehat{r}^{AA^4}$ versus $\widehat{r}^{DD^2}$, and $\widehat{r}^{AD^2}$ versus $\widehat{r}^{DA^2}$) we dash one of the lines and overlay it.

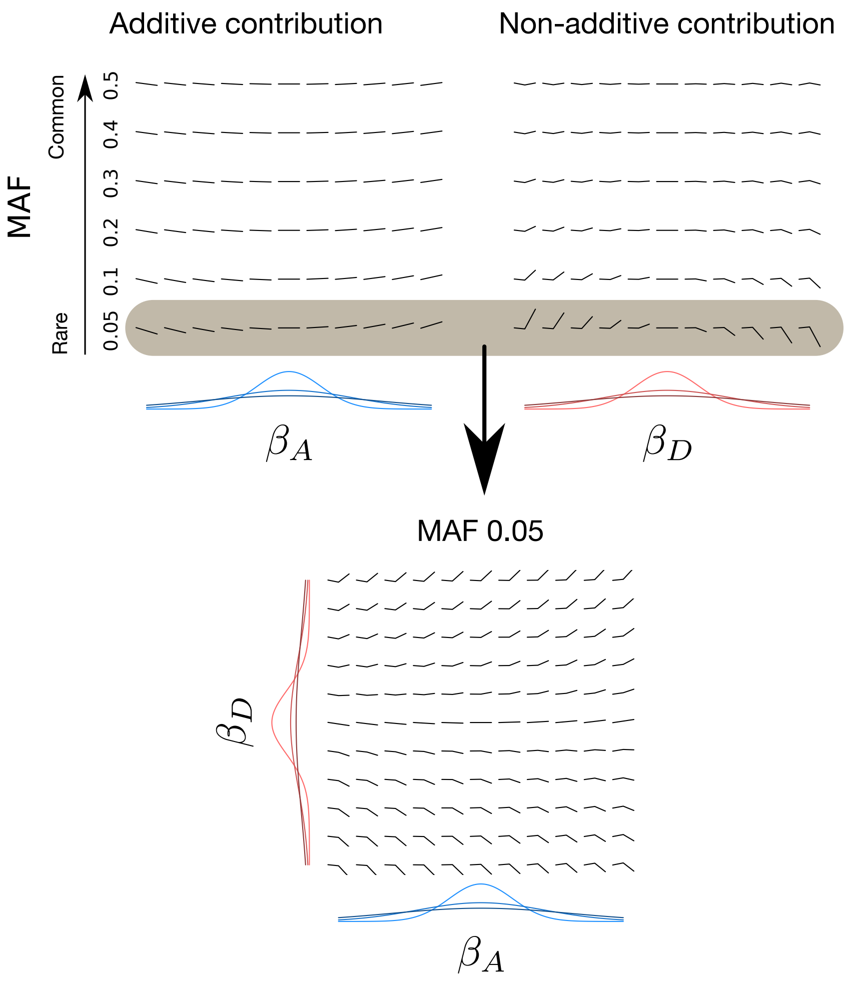


**Figure** S2: Plots to inform intuition. The top two plots display the uncorrelated true additive and dominance contributions of the causal effect of a SNP on a trait for varying values of $\beta_A$ and $\beta_D$ respectively for a given MAF at each genotype dosage (shown in each mini-subplot). Values of $\beta_A$ and $\beta_D$ range from -1 to 1 in equally spaced intervals), and MAF is varied on the y-axis (0.05, 0.1, 0.2, 0.3, 0.4, and 0.5). Moving from dark to light, distributions at the foot of the top two plots are the sampling distributions for $\beta_A$ and $\beta_D$ when $h_A^2$ and $h_D^2$ are 0.05, 0.2 and 0.4. In the lower plot we display the summation of the additive and dominance contributions as we vary $(\beta_A$ and $\beta_D)$ over [-1,1][-1,1] at a site with the MAF highlighted in grey in the upper plots (0.05)

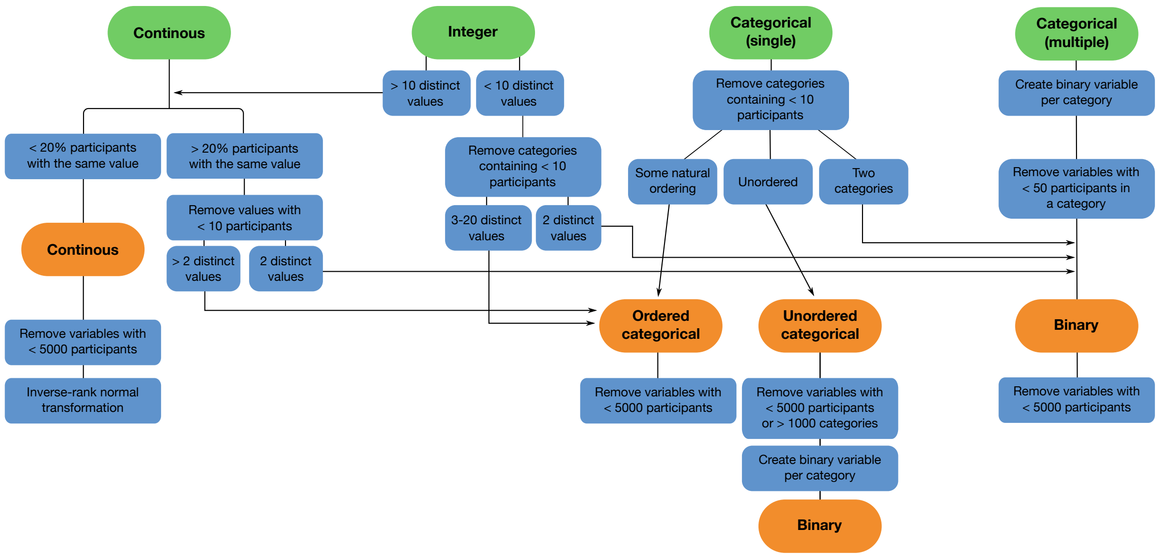


**Figure** S3: Phenotype curation using an adapted version of PHESANT. The flowchart summarises the filtering and transformation steps used to parse the raw UK biobank phenotype data. Figure adapted from (*58*).

unless it is a job code (96 such phenotypes).

We then remove the raw encoding for continuous phenotypes, and filter on the case/control count to ensure that binary and categorical phenotypes have at least 3,000 cases (or members of the category) and controls. This was to ensure some power to detect dominance effects at a MAF of at least 0.05, as it results in at least $\sim 8$ samples who are homozygous alternate at every site above MAF 0.05. We include only continuous phenotypes with at least 50,000 non-missing data points. Finally, we exclude all ordinal variables to avoid non-linear phenotypic encodings. The resultant breakdown of phenotypes by category is displayed in the third column of Table S1.

#### Sample and genotype curation

##### Sample QC

Beginning with the 487,409 individuals with phased and imputed genotype data, we restricted to the collection of unrelated individuals with low autosomal missingness rates used for principal components analysis (PCA) by Bycroft *et al.* (*16*). Using the first six principal components provided by the UK Biobank genetics team (UK biobank phenotype ID 22009, we then filtered individuals of European ancestry individuals. In particular, we define a six-dimensional ellipse of the top six PCs by centering on the mean among individuals who self-identified as white and British. We then remove samples that lie outside seven standard deviations on any of these six PC axes. Standard deviations were computed using those individuals reporting as white British. We also remove individuals who self-identify as an ethnicity other than white from our GWAS analyses. See (*23*) for reasoning and further details. This definition resulted in a moderate increase in sample size when compared to the white British subset defined in Bycroft *et al.* (*16*), predominantly due to the inclusion of individuals reporting as Irish or another white ethnicity.

Finally, following our ancestry based filtering regime, we remove samples who withdrew from UK Biobank participation as well as those individuals who were omitted from phasing and imputation. Following all of these sample QC filtering steps, 361,194 positive individuals were retained for the dominance GWAS.

##### Genotype QC

Over 92 million imputed variants across the autosomes and chromosome X are available for analysis. As a starting point for our initial collection of GWASes, we subset to variants with

MAF $> 0.1\%$ in the subset of individuals defined in the sample QC procedure, Hardy-Weinberg $P$-value $> 1 \times 10^{-10}$ and an info score $> 0.8$ from the UK Biobank SNP manifest file. The exception being variants annotated using Ensembl VEP (*59*) consequence as having protein-truncating or missense consequences: at these variants we loosened our MAF filter to $> 1 \times 10^{-6}$. Following this collection of filtering steps, $13.7$ million variants were retained for our GWASes. A summary of the sample and genotype QC is displayed in Fig. S4.

#### Evaluation of additive and dominance marginal effect sizes

Additive genome-wide association studies were carried out using hail version 0.1 (*60, 61*) using linear regression with the `linreg3` function. Dominance genome-wide association studies were also carried out in hail version 0.1 using a custom alteration to the scala code to recode genotypes under the dominance encoding on the fly (*62*). For each phenotype (where appropriate) we run a both sex, male, and female GWAS. For the both sex analysis, we include $age$, $age^2$, $sex$, $age \times sex$, $age^2 \times sex$, and the first 20 principal components as covariates. In both the male and female specific GWAS, we include $age$, $age^2$, and the first 20 principal components as covariates.

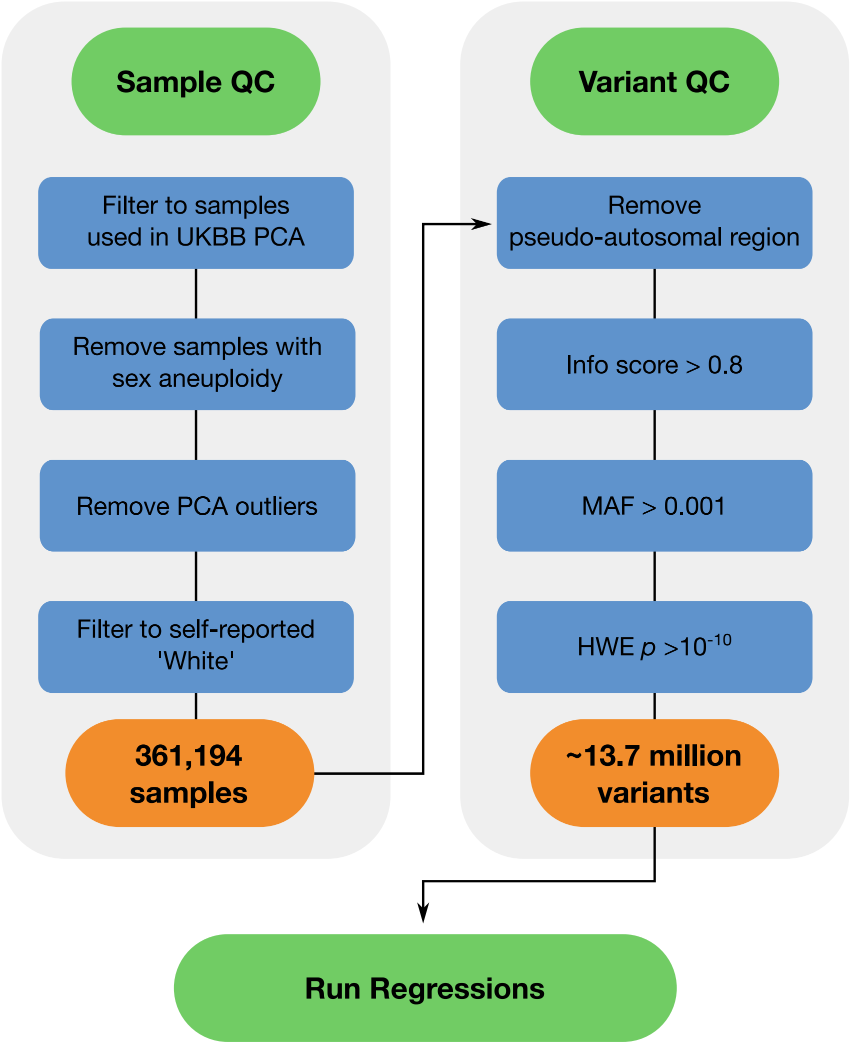


**Figure** S4: Sample and variant QC summary. Flowchart summarising sample and variant QC steps prior to performing GWAS on each curated trait. Following sample and variant QC, 361,194 samples and 13.7 million variants remain.

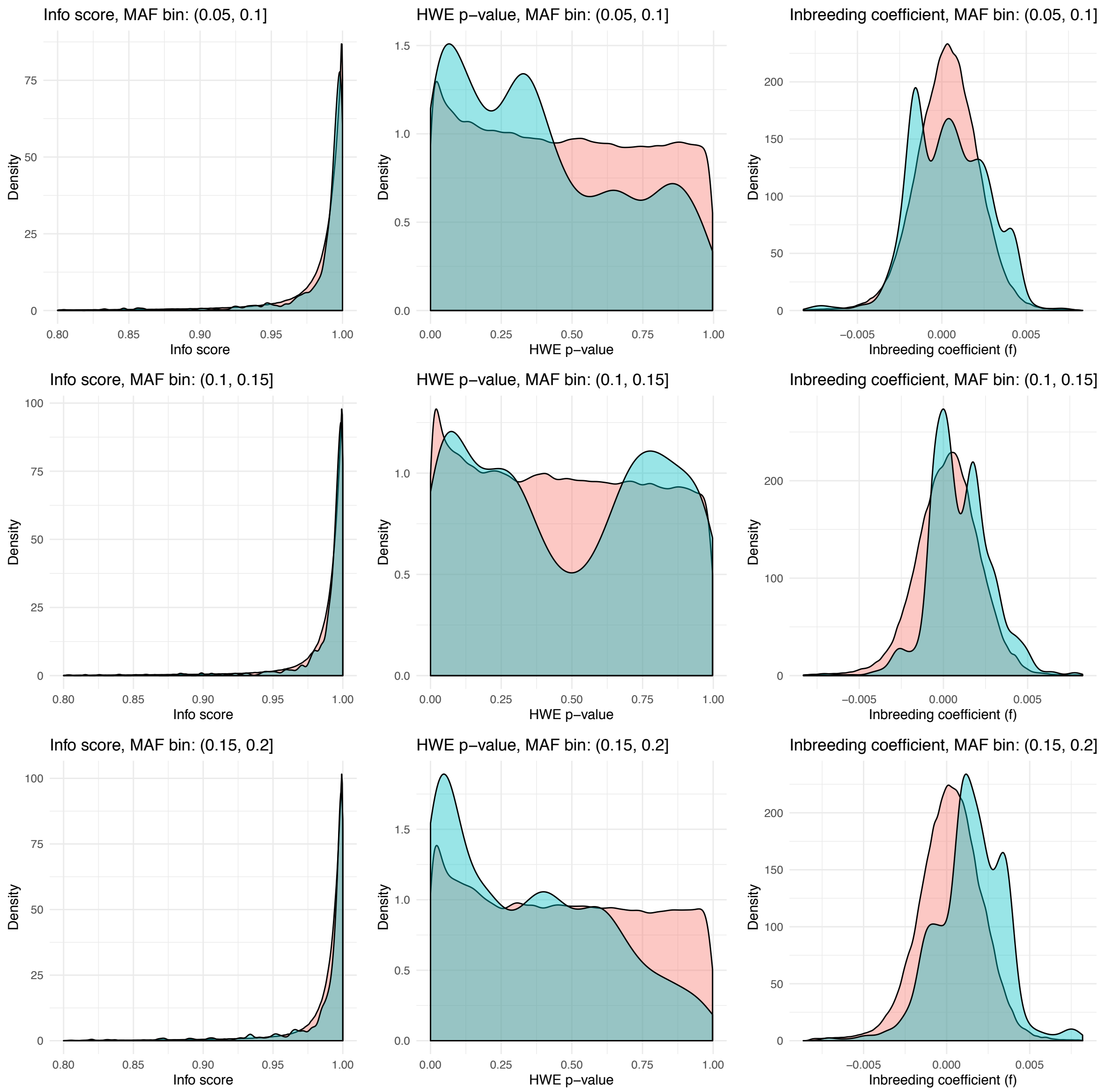


**Figure** S5: Comparing the distribution of info score, $P$-HWE, and $f$-statistic between sites with and without a genome-wide significant ($P < 5 \times 10^{-8}$) non-additive signal: MAF $\in [0.05, 0.10), [0.10, 0.15), [0.15, 0.20)$. We display the distribution of info score, $P$-HWE, and $f$-statistic in each column respectively. Variants were grouped into 5% MAF bins in each row. In each panel, teal and pink displays the distribution of the metric at sites with and without genome-wide significant non-additive associations respectively.

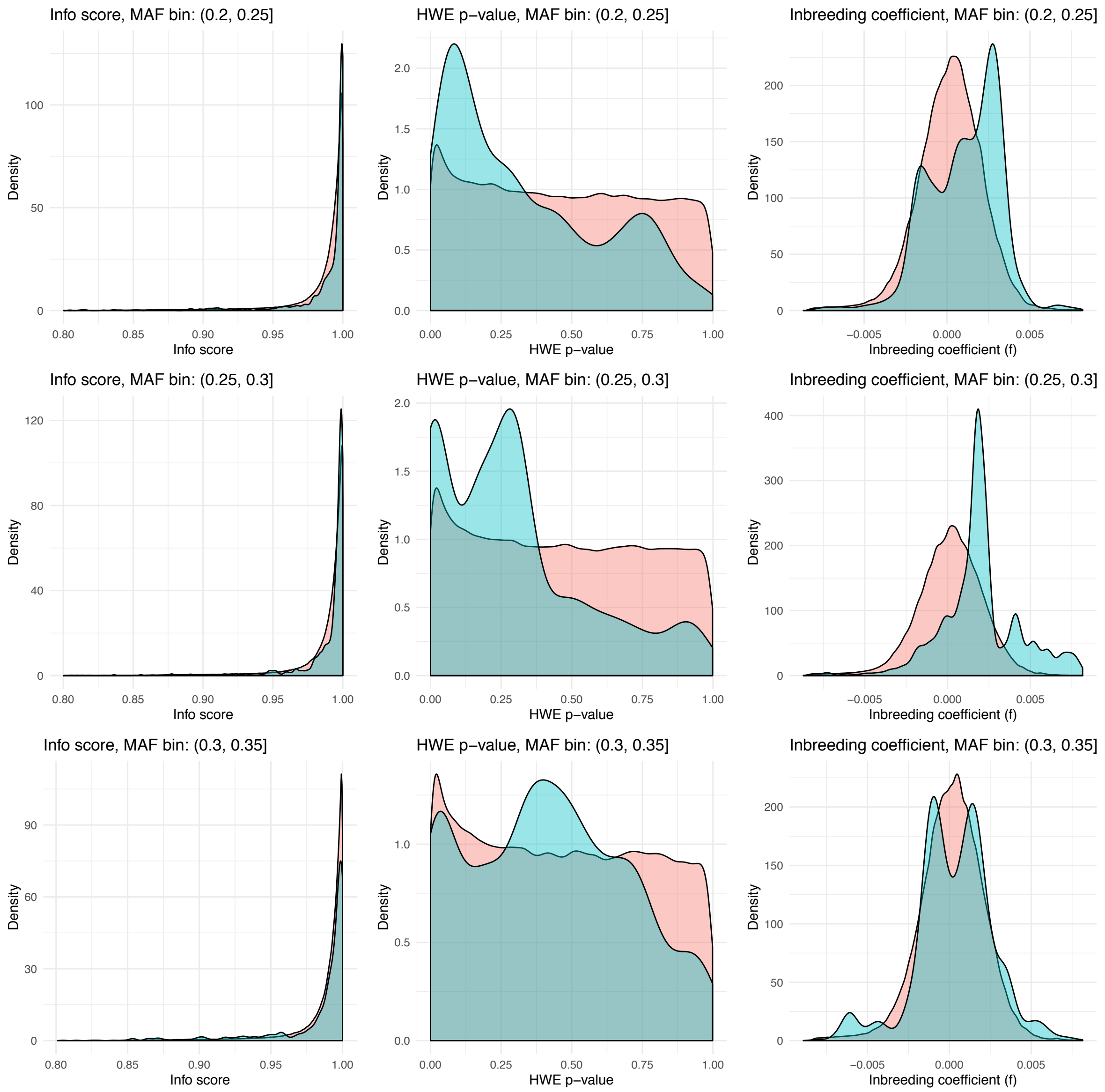


**Figure** S6: Comparing the distribution of info score, $P$-HWE, and $f$-statistic between sites with and without a genome-wide significant ($P < 5 \times 10^{-8}$) non-additive signal: MAF $\in [0.20, 0.25), [0.25, 0.30), [0.30, 0.35)$. We display the distribution of info score, $P$-HWE, and $f$-statistic in each column respectively. Variants were grouped into 5% MAF bins in each row. In each panel, teal and pink displays the distribution of the metric at sites with and without genome-wide significant non-additive associations respectively.

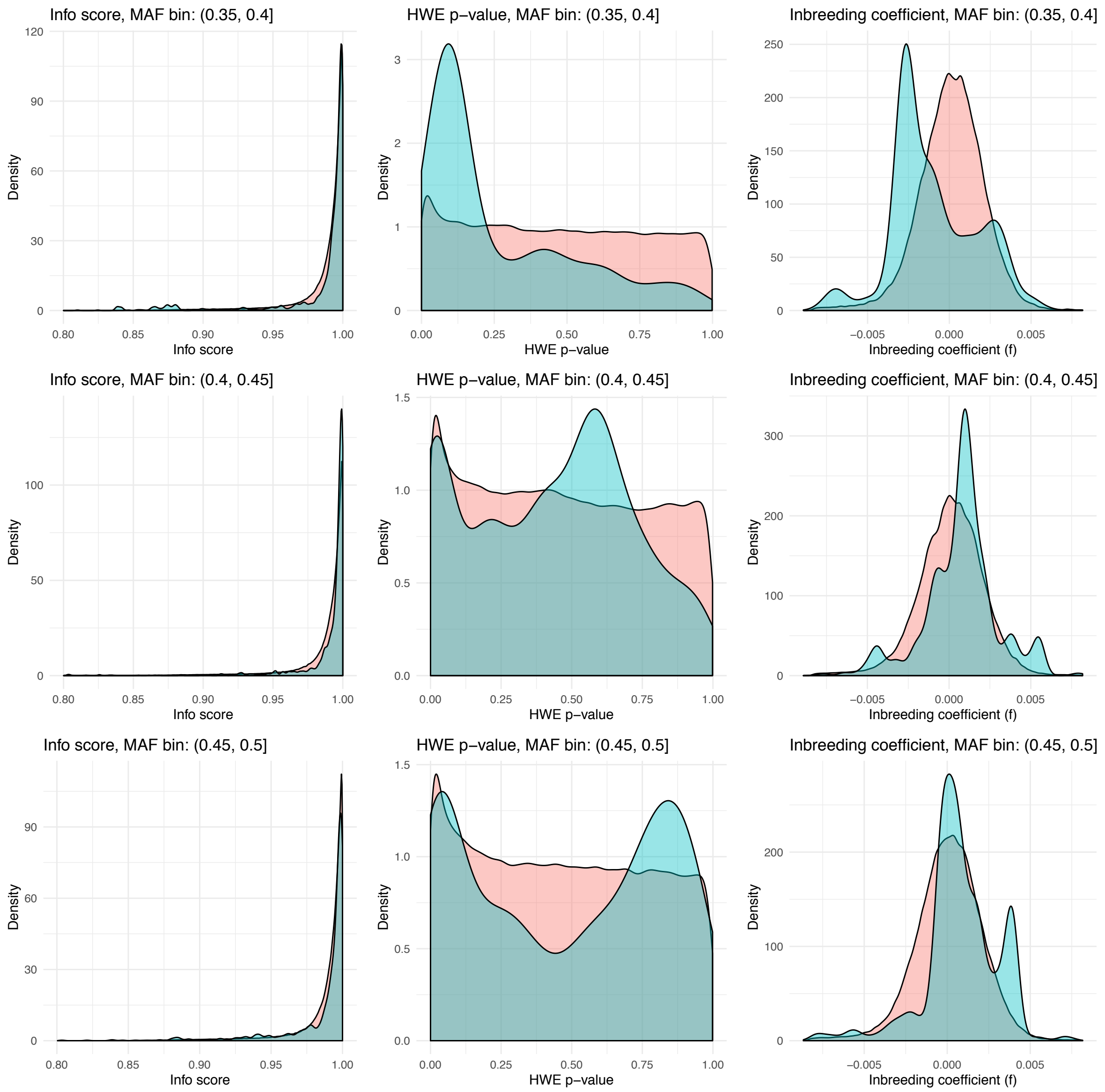


**Figure** S7: Comparing the distribution of info score, $P$-HWE, and $f$-statistic between sites with and without a genome-wide significant ($P < 5 \times 10^{-8}$) non-additive signal: MAF $\in [0.35, 0.40), [0.40, 0.45), [0.45, 0.50)$. We display the distribution of info score, $P$-HWE, and $f$-statistic in each column respectively. Variants were grouped into 5% MAF bins in each row. In each panel, teal and pink displays the distribution of the metric at sites with and without genome-wide significant non-additive associations respectively.

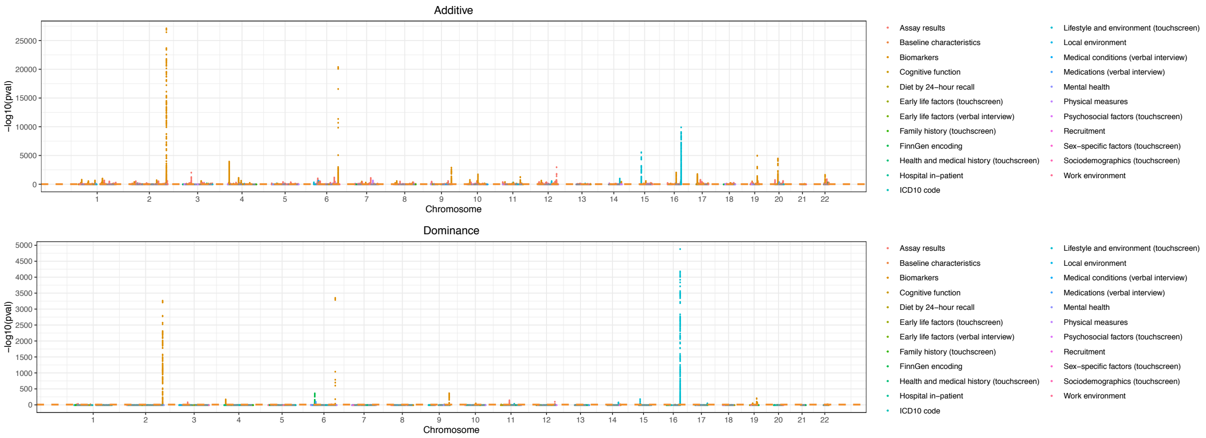


**Figure** S8: Aggregate Manhattan plots for additive and non-additive marginal effect sizes. We determine the most extreme t-statistic for marginal effect sizes across all well covered traits at each site with MAF $> 0.05$, and plot the associated $-\log_{10}(p)$ value. Traits are grouped into broad categories as defined in the UK Biobank data showcase and coloured according to the legend.

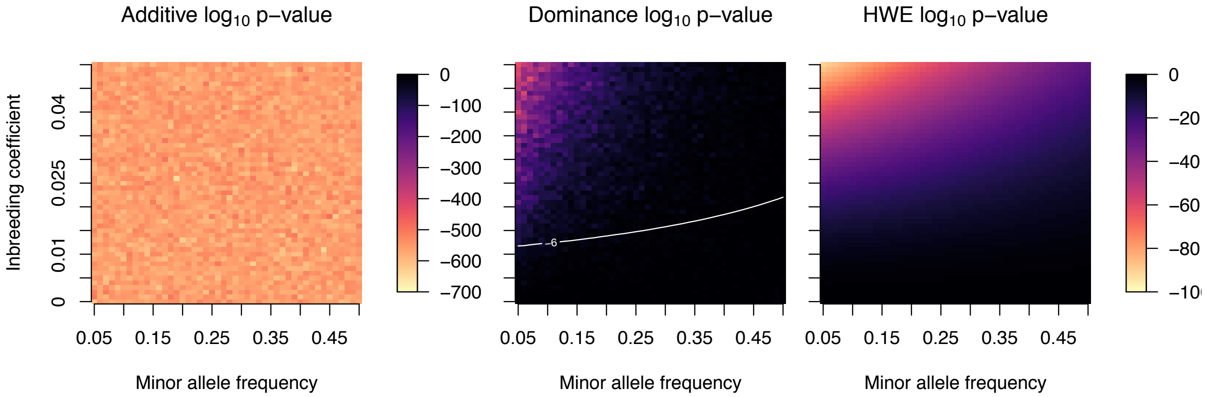


**Figure** S9: $P$-values for dominance effect sizes under an additive model with genotype frequencies out of HWE due to genotyping error. A proportion $\phi$ of heterogygote calls are moved into the homozygous alternate cluster. Additive, dominance, and HWE $P$-values are displayed for varying MAF and error proportion given $n = 50,000$ samples and a true effect size $\beta = 0.05$ on a standardised phenotype. The additive $P$-value heat map is coloured according to the first legend, the second two heat maps are coloured according to the second legend.

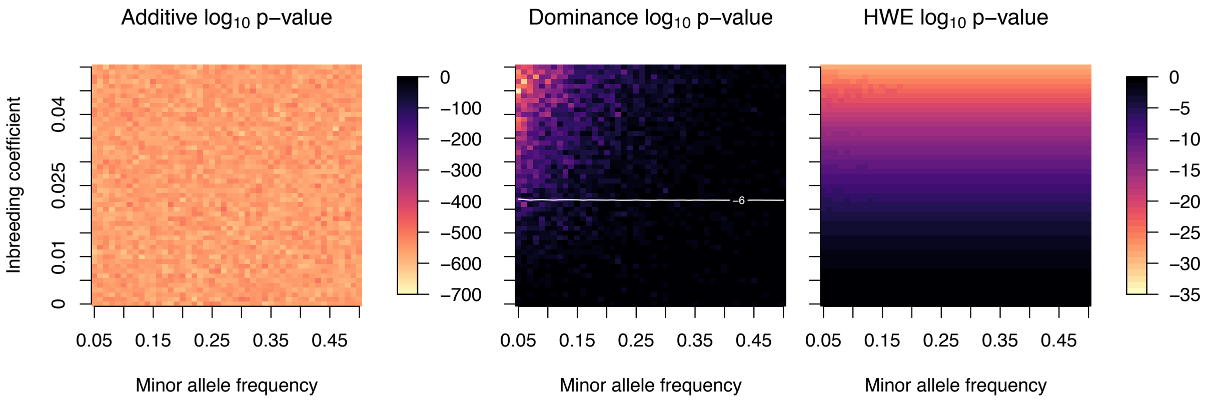


**Figure** S10: $P$-values for dominance effect sizes under an additive model with genotype frequencies out of HWE. Additive, dominance, and HWE $P$-values are displayed for varying MAF and inbreeding coefficient $(f)$ given $n = 50,000$ samples and a true effect size $\beta = 0.05$ on a standardised phenotype. Plots display heat maps of the additive and dominance $P$-values for association, and HWE $P$-values, respectively. The additive $P$-value heat map is coloured according to the first legend, the second two heat maps are coloured according to the second legend.

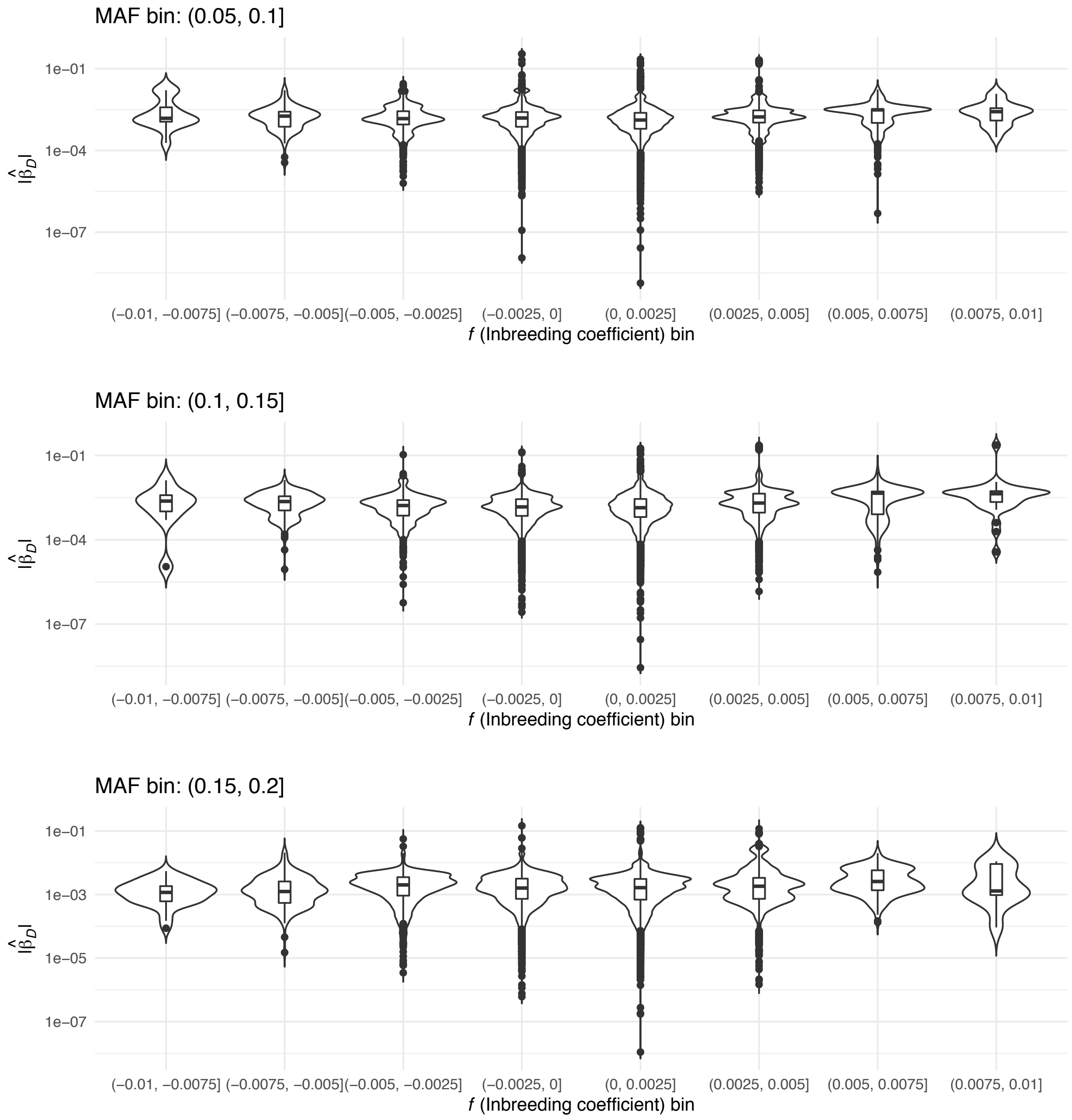


**Figure** S11: Marginal non-additive effect size estimates at extremely significant additive associations $(p < 10^{-30})$: MAF $\in (0.05, 0.1], (0.1, 0.15], (0.15, 0.2]$. At each site we determined the most significant additive associations across all well covered phenotypes. We then examined the distribution of $|\hat{\beta}_D|$ binning by MAF and inbreeding coefficient $(f)$. If deviations from HWE leading to genome wide significant effects are widespread, we expect to see a trend between deviation of $f$-statistic from 0 and $|\hat{\beta}_D|$ which should be particularly apparent at low MAF.

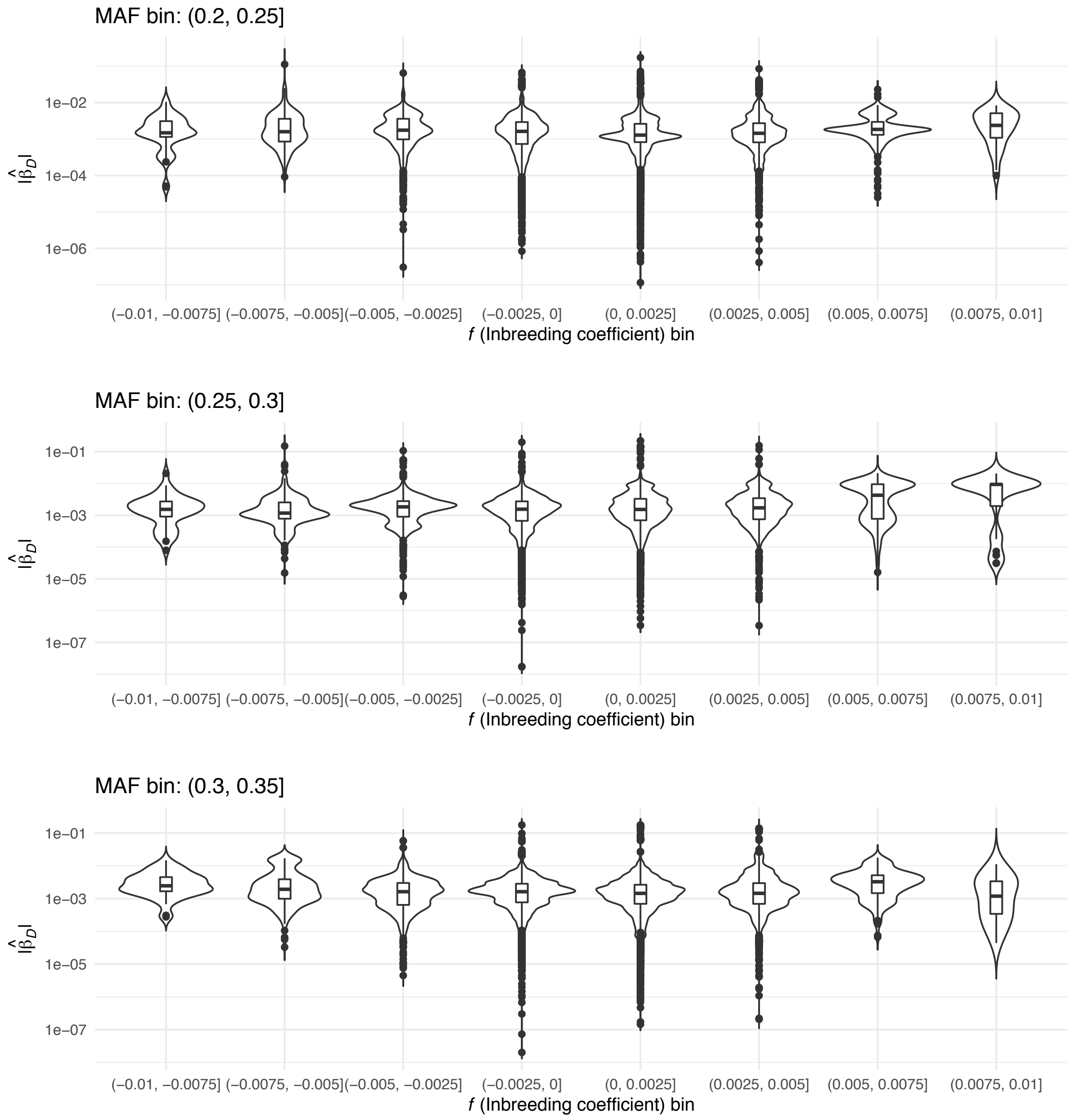


**Figure** S12: Marginal non-additive effect size estimates at extremely significant additive associations ($p < 10^{-30}$): MAF $\in (0.2, 0.25], (0.25, 0.3], (0.3, 0.35]$. At each site we determined the most significant additive associations across all well covered phenotypes. We then examined the distribution of $|\hat{\beta}_D|$ binning by MAF and inbreeding coefficient $(f)$. If deviations from HWE leading to genome wide significant effects are widespread, we expect to see a trend between deviation of $f$-statistic from 0 and $|\hat{\beta}_D|$ which should be particularly apparent at low MAF.

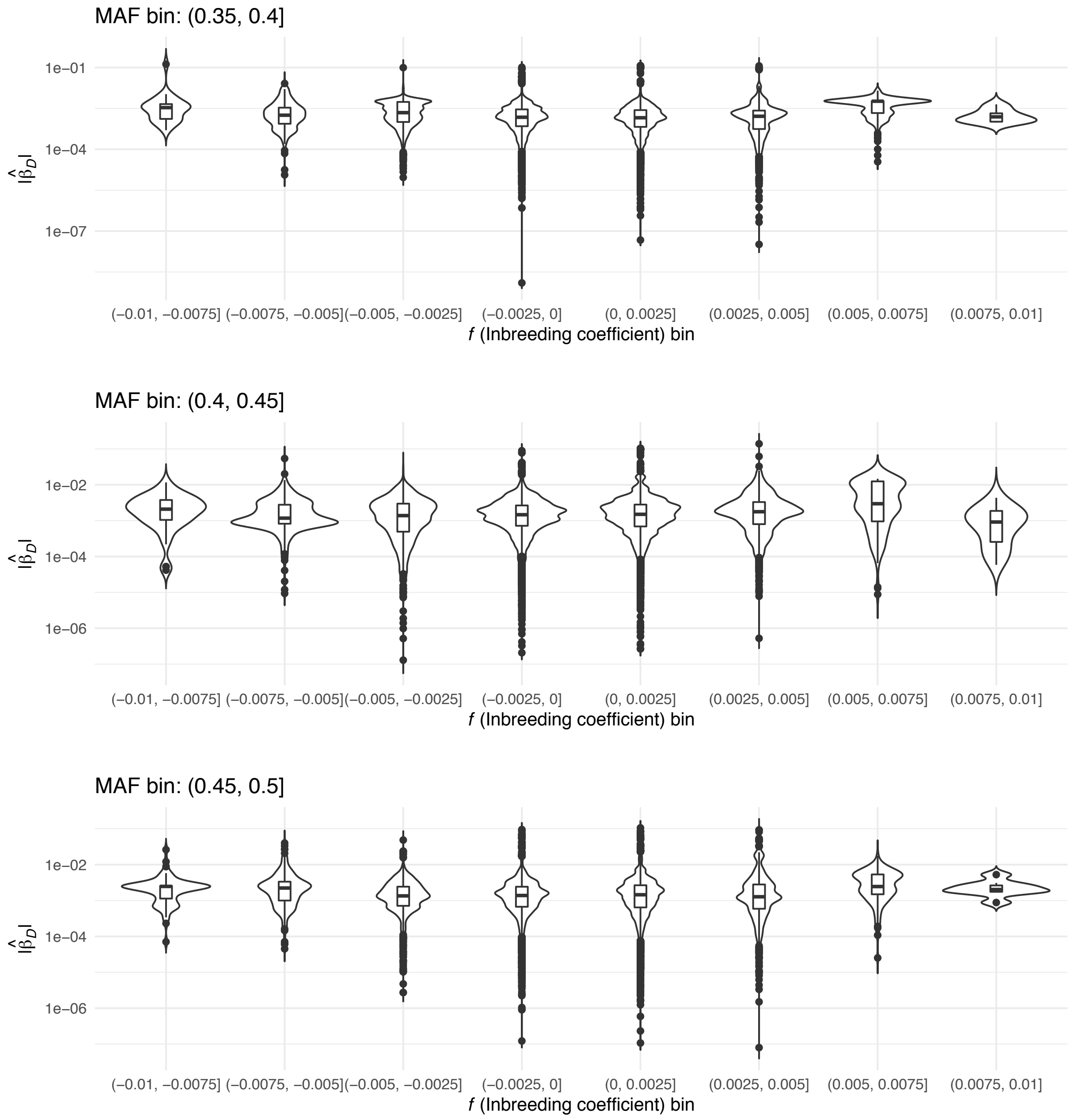


**Figure** S13: Marginal non-additive effect size estimates at extremely significant additive associations $(p < 10^{-30})$: MAF $\in (0.35, 0.4], (0.4, 0.45], (0.45, 0.5]$. At each site we determined the most significant additive associations across all well covered phenotypes. We then examined the distribution of $|\hat{\beta}_D|$ binning by MAF and inbreeding coefficient $(f)$. If deviations from HWE leading to genome wide significant effects are widespread, we expect to see a trend between deviation of $f$-statistic from 0 and $|\hat{\beta}_D|$ which should be particularly apparent at low MAF.

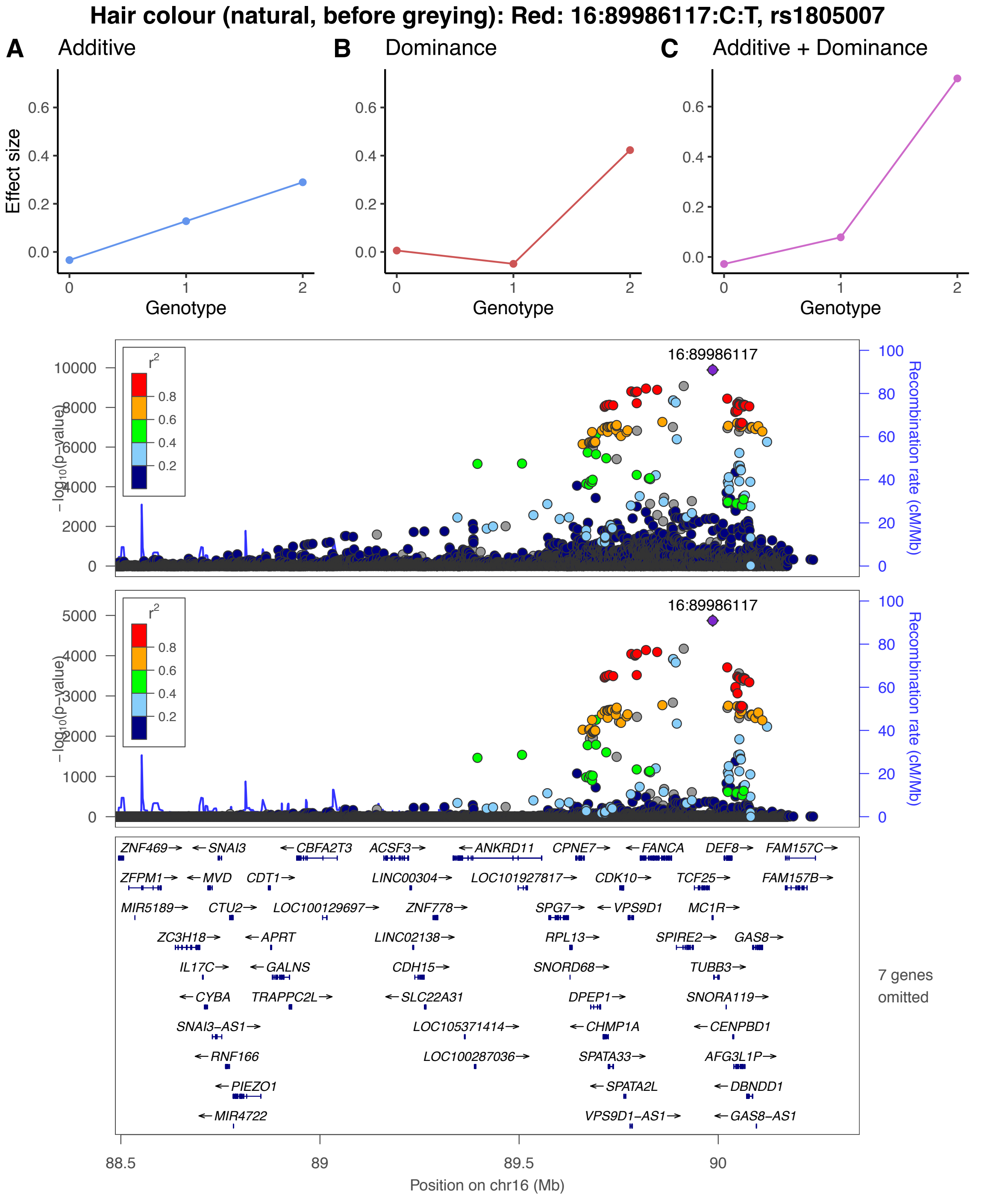


**Figure** S14: Red hair (UK biobank phenotype code 1747), MC1R locus. Locus-zoom plots of $-\log_{10}(p)$ values associated to the marginal additive and dominance effect sizes in the upper and lower panels respectively. In each plot the legend refers to additive LD from the lead SNP in the region.

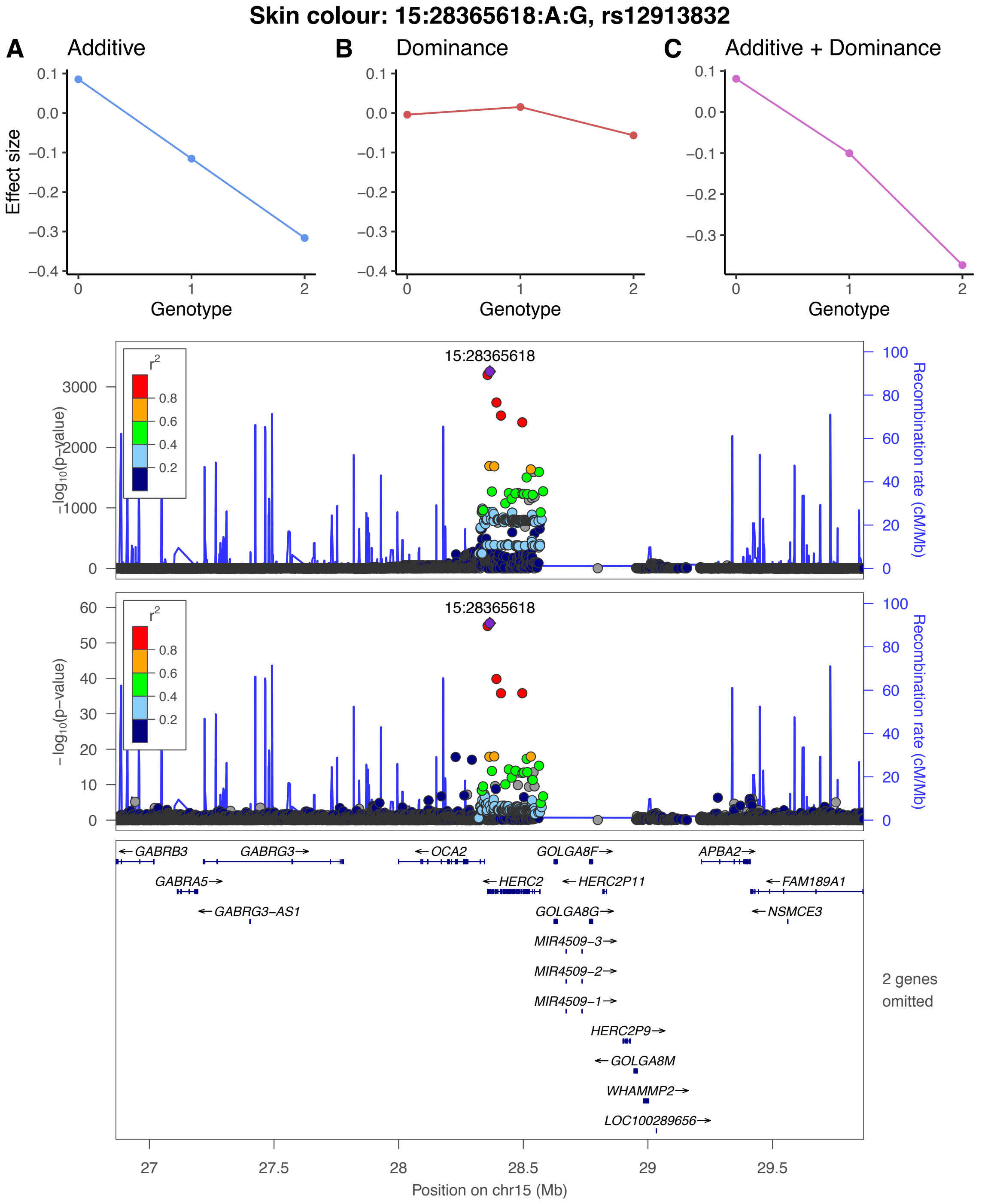


**Figure** S15: Skin Colour (UK Biobank phenotype code 1717), HERC2 locus. Locus-zoom plots of $-\log_{10}(p)$ values associated to the marginal additive and dominance effect sizes in the upper and lower panels respectively. In each plot the legend refers to additive LD from the lead SNP in the region.

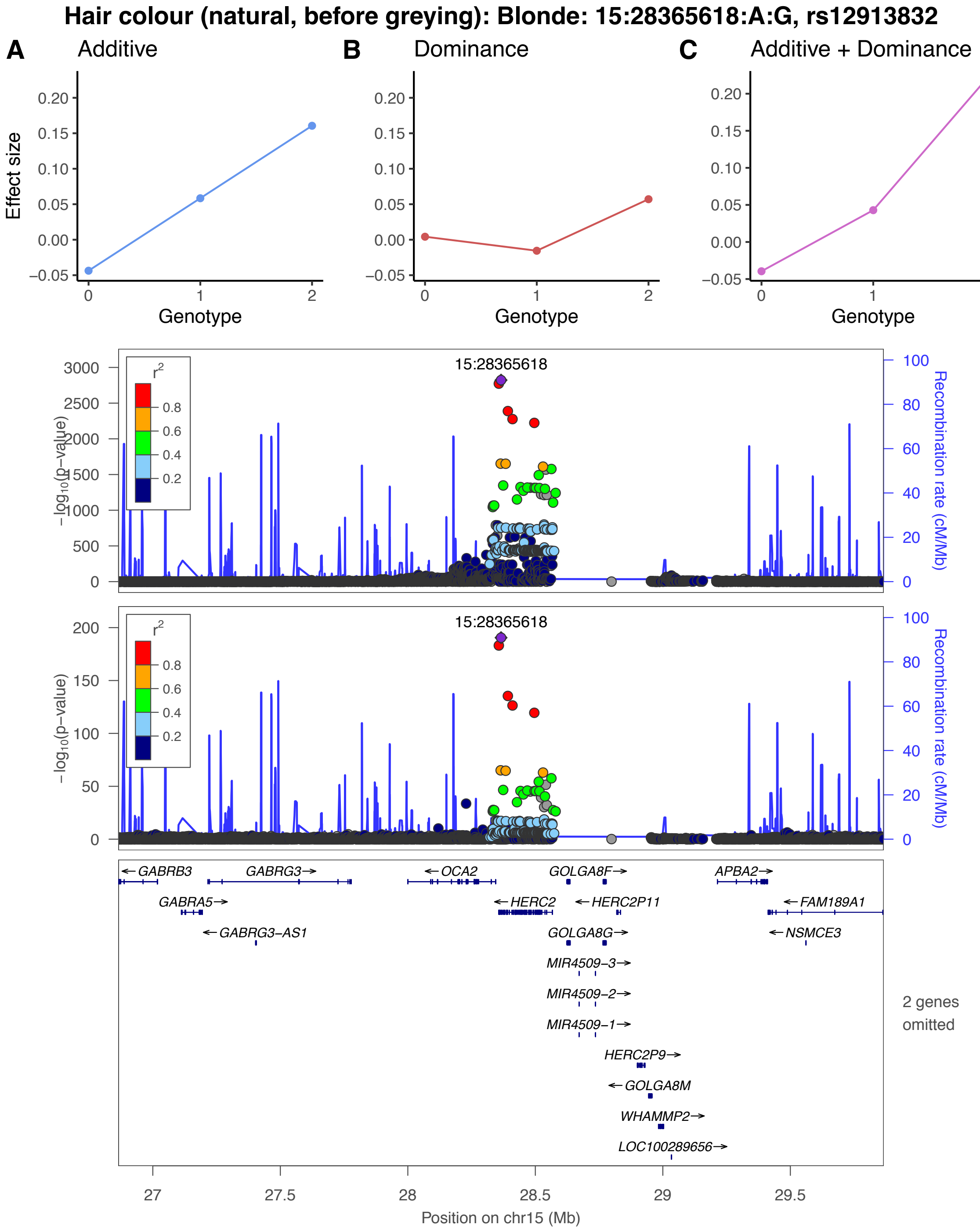


**Figure** S16: Blonde hair (UK Biobank phenotype code 1747), HERC2 locus. Locus-zoom plots of $-\log_{10}(p)$ values associated to the marginal additive and dominance effect sizes in the upper and lower panels respectively. In each plot the legend refers to additive LD from the lead SNP in the region.

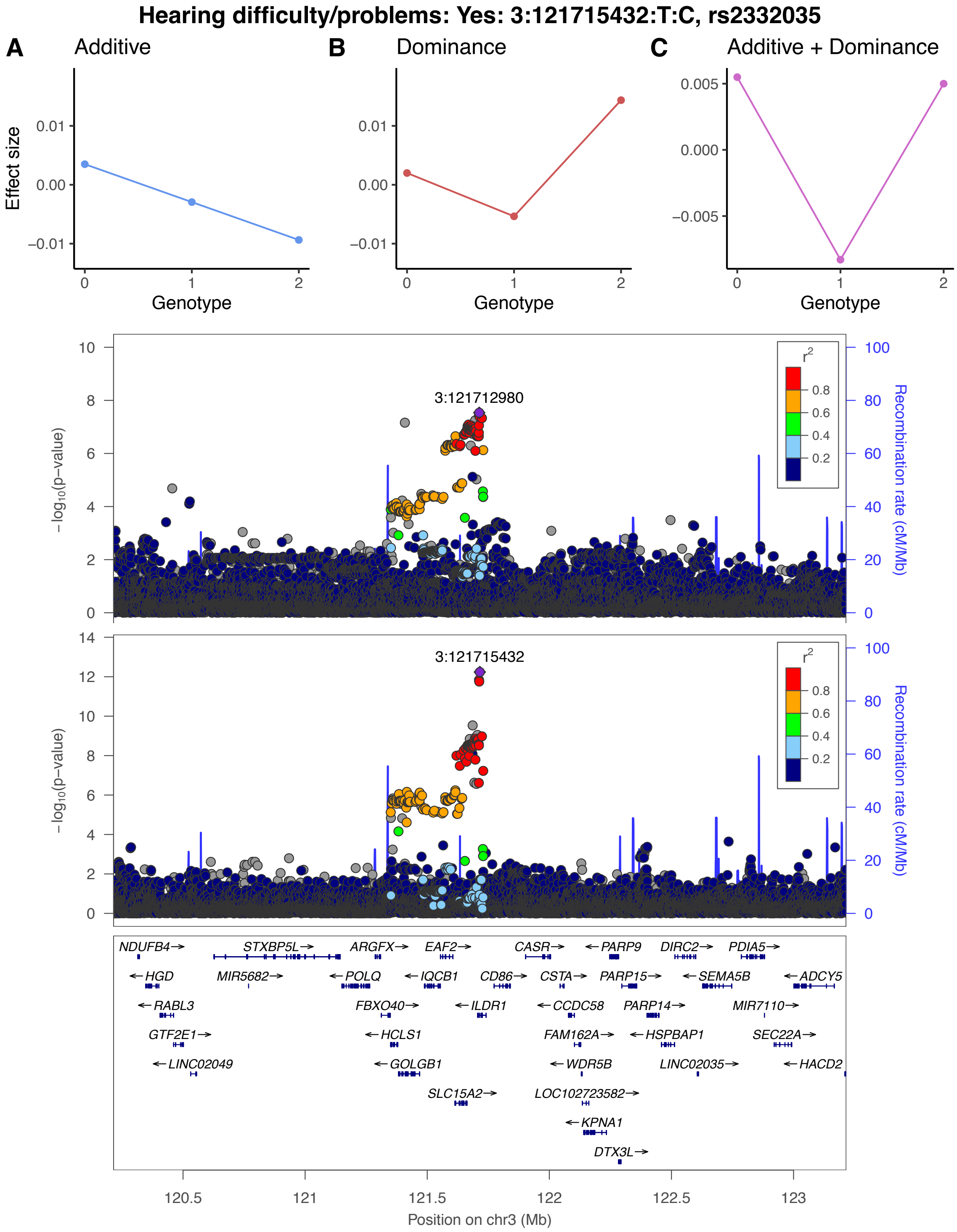


**Figure** S17: Hearing difficulty/problems (UK Biobank phenotype code 2247), ILDR1 locus. Locus-zoom plots of $-\log_{10}(p)$ values associated to the marginal additive and dominance effect sizes in the upper and lower panels respectively. In each plot the legend refers to additive LD from the lead SNP in the region.

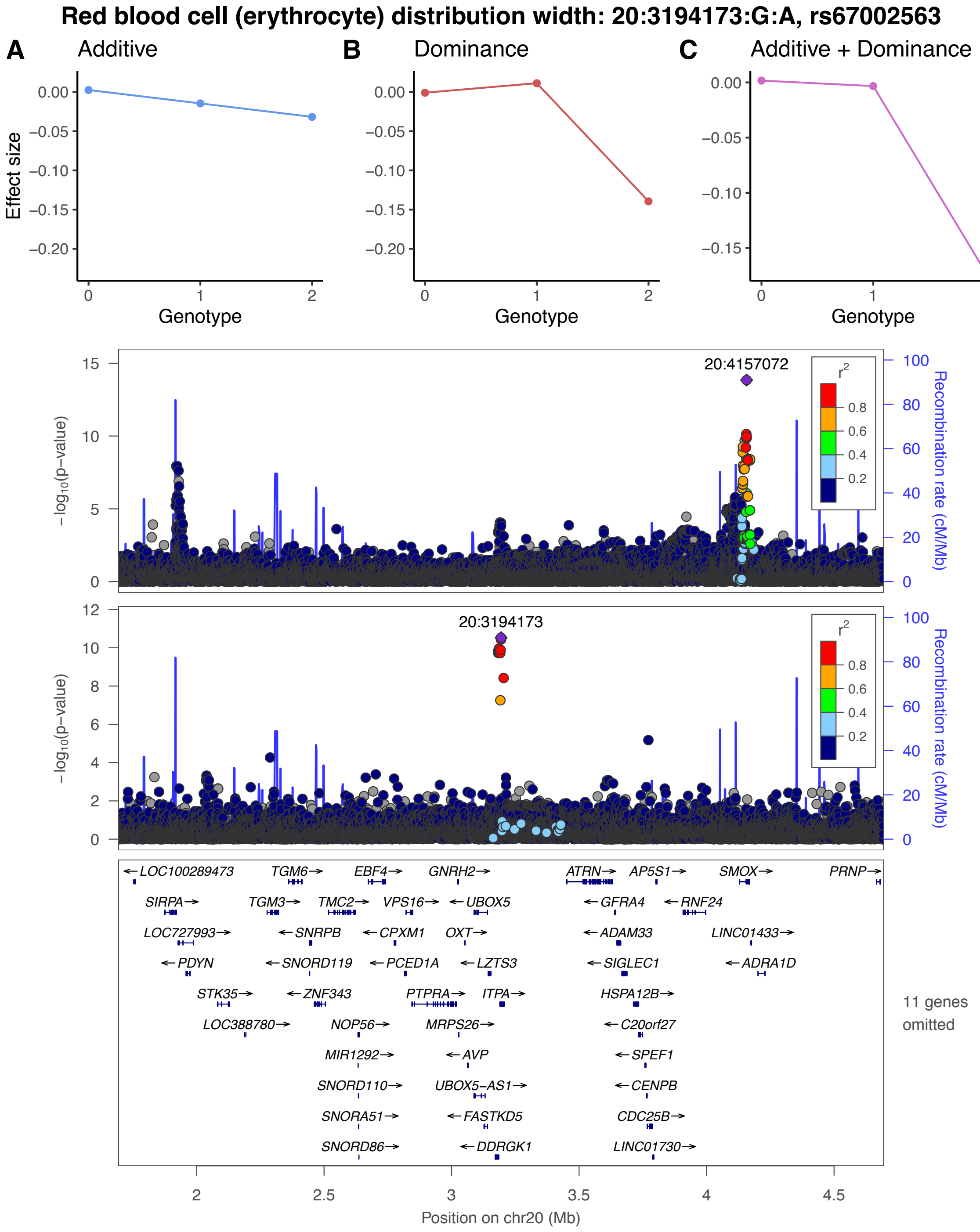


**Figure** S18: Red blood cell distribution width (UK Biobank phenotype code 30070), ITPA locus. Locus-zoom plots of $-\log_{10}(p)$ values associated to the marginal additive and dominance effect sizes in the upper and lower panels respectively. In each plot the legend refers to additive LD from the lead SNP in the region.

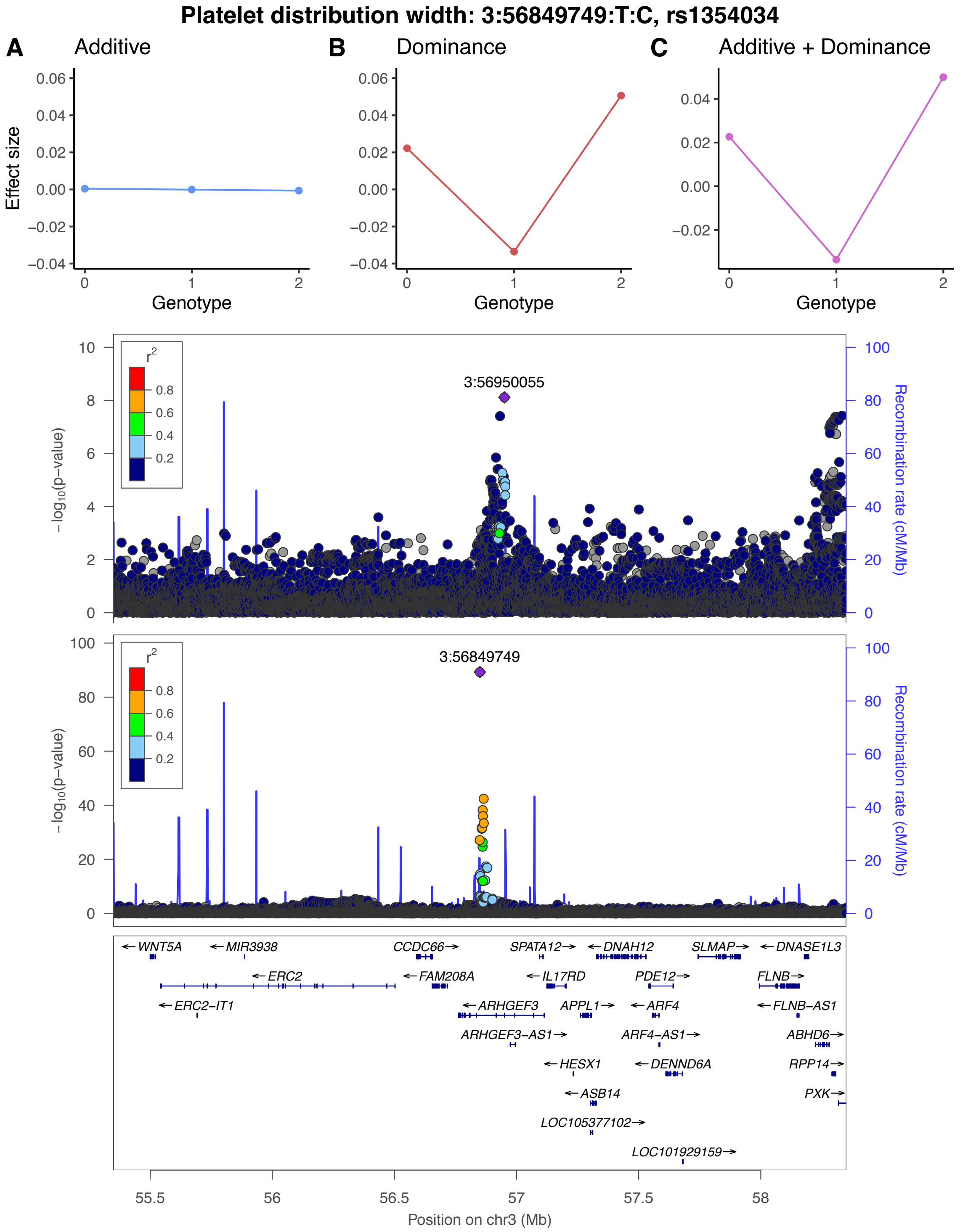


**Figure** S19: Platelet distribution width (UK Biobank phenotype code 30110), ARHGEF3 locus. Locus-zoom plots of $-\log_{10}(p)$ values associated to the marginal additive and dominance effect sizes in the upper and lower panels respectively. In each plot the legend refers to additive LD from the lead SNP in the region.

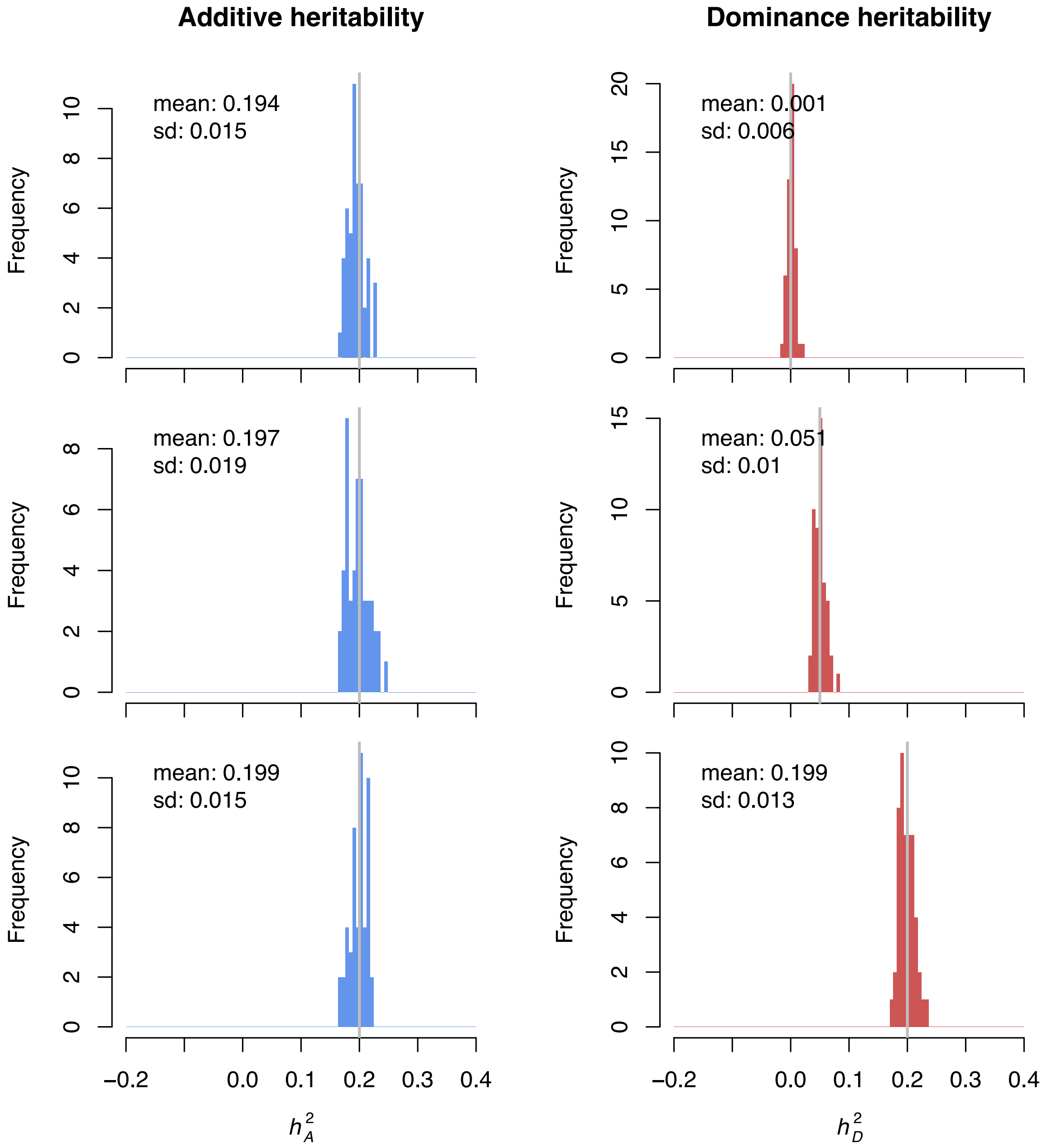


**Figure** S20: Msprime simulation study: Continuous phenotype, all SNPs causal. Phenotypes are simulated under an infinitesimal architecture. Histograms of additive and dominance heritability estimates across the 50 independent phenotype simulations are shown in the left and right columns respectively. The grey line in the true underlying heritability, mean and standard deviations of the point estimates across runs are inset in each panel.

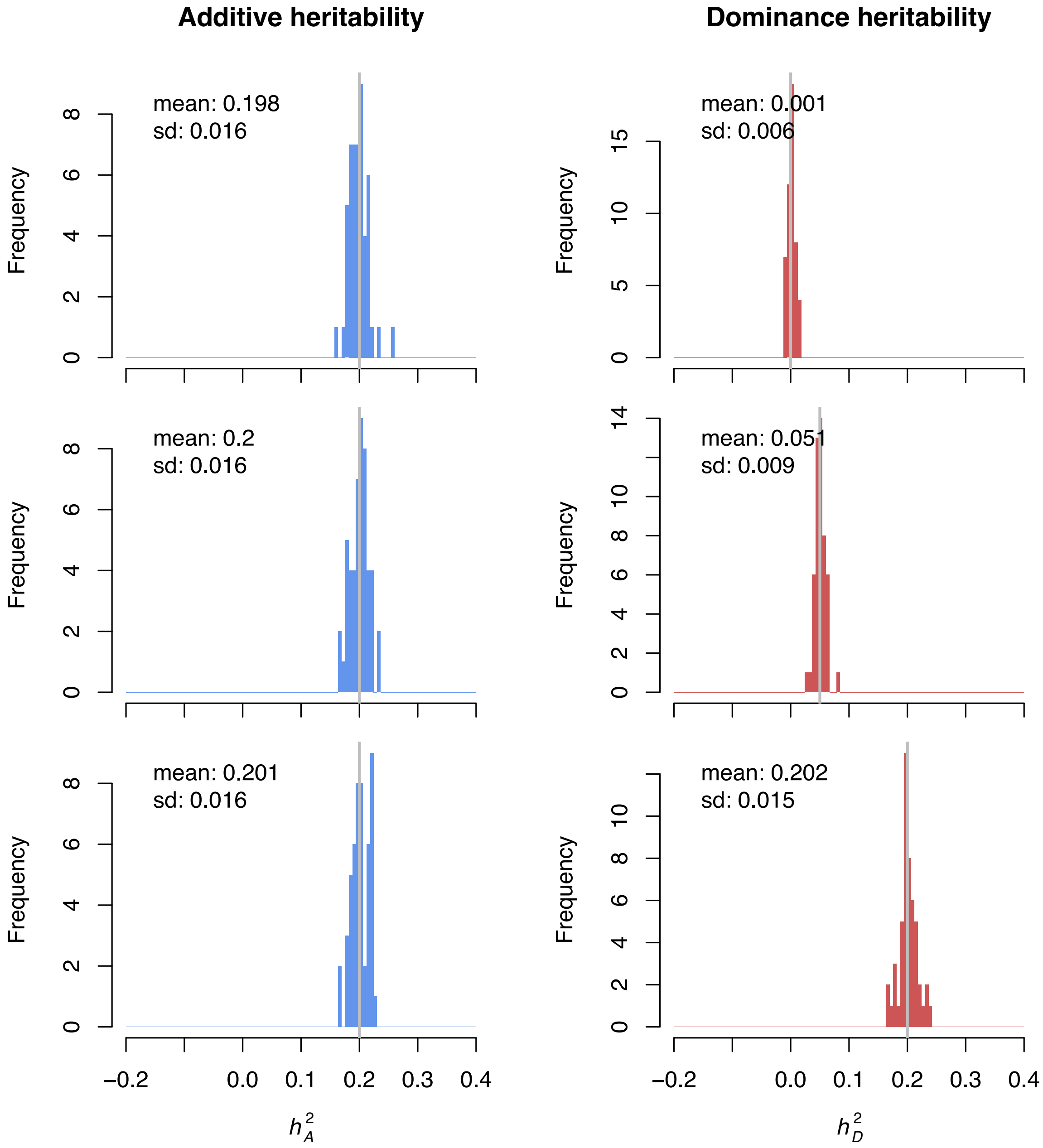


**Figure** S21: Msprime simulation study: Continuous phenotype, 10% SNPs causal. Phenotypes are simulated under a spike and slab architecture with 10% causal sites. Histograms of additive and dominance heritability estimates across the 50 independent phenotype simulations are shown in the left and right columns respectively. The grey line in the true underlying heritability, mean and standard deviations of the point estimates across runs are inset in each panel.

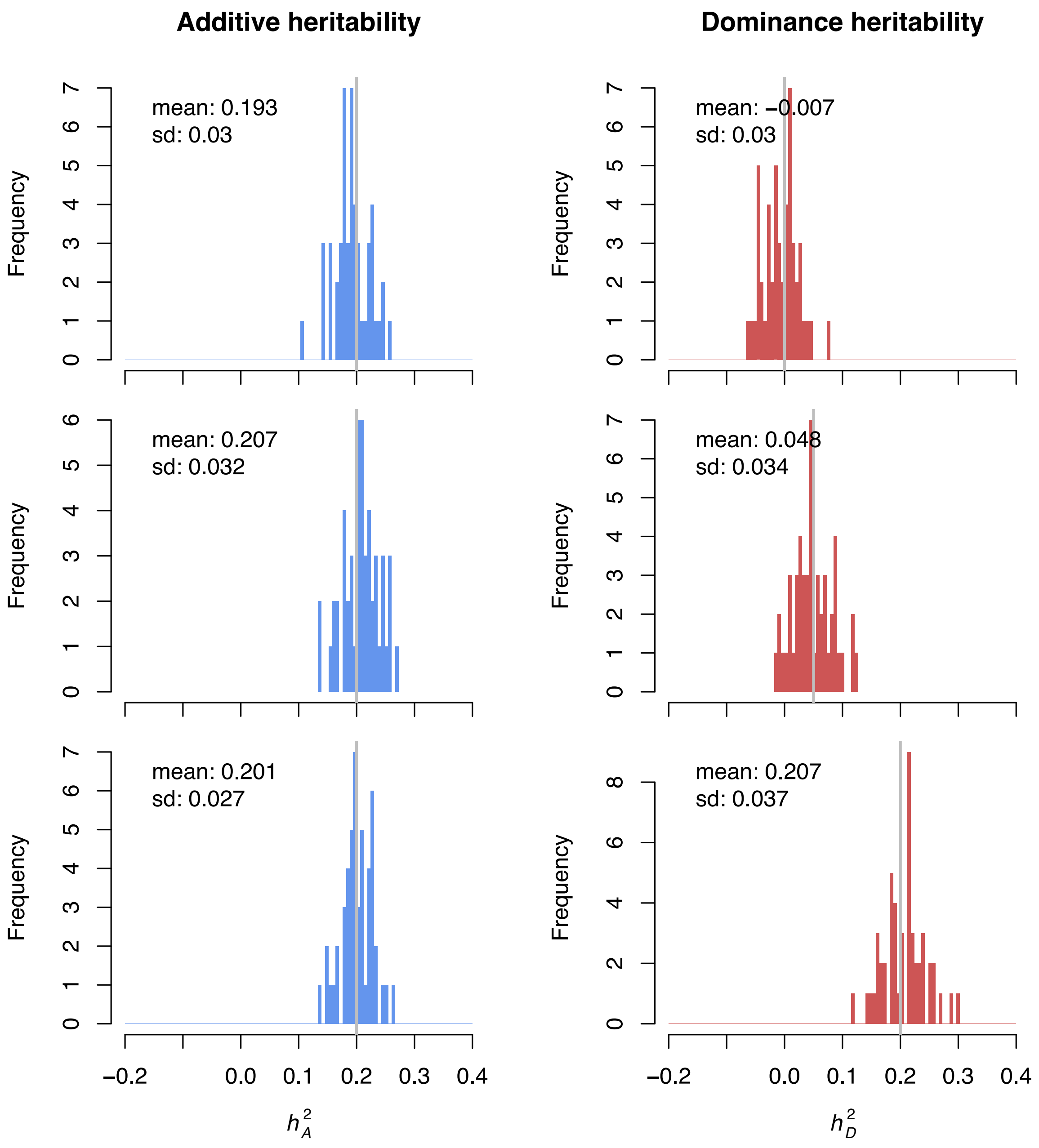


**Figure** S22: Msprime simulation study: Case-control phenotype with 6% prevalence, no case ascertainment, all SNPs causal. Phenotypes are simulated under an infinitesimal architecture. Histograms of additive and dominance heritability estimates across the 50 independent phenotype simulations are shown in the left and right columns respectively. The grey line in the true underlying heritability, mean and standard deviations of the point estimates across runs are inset in each panel.

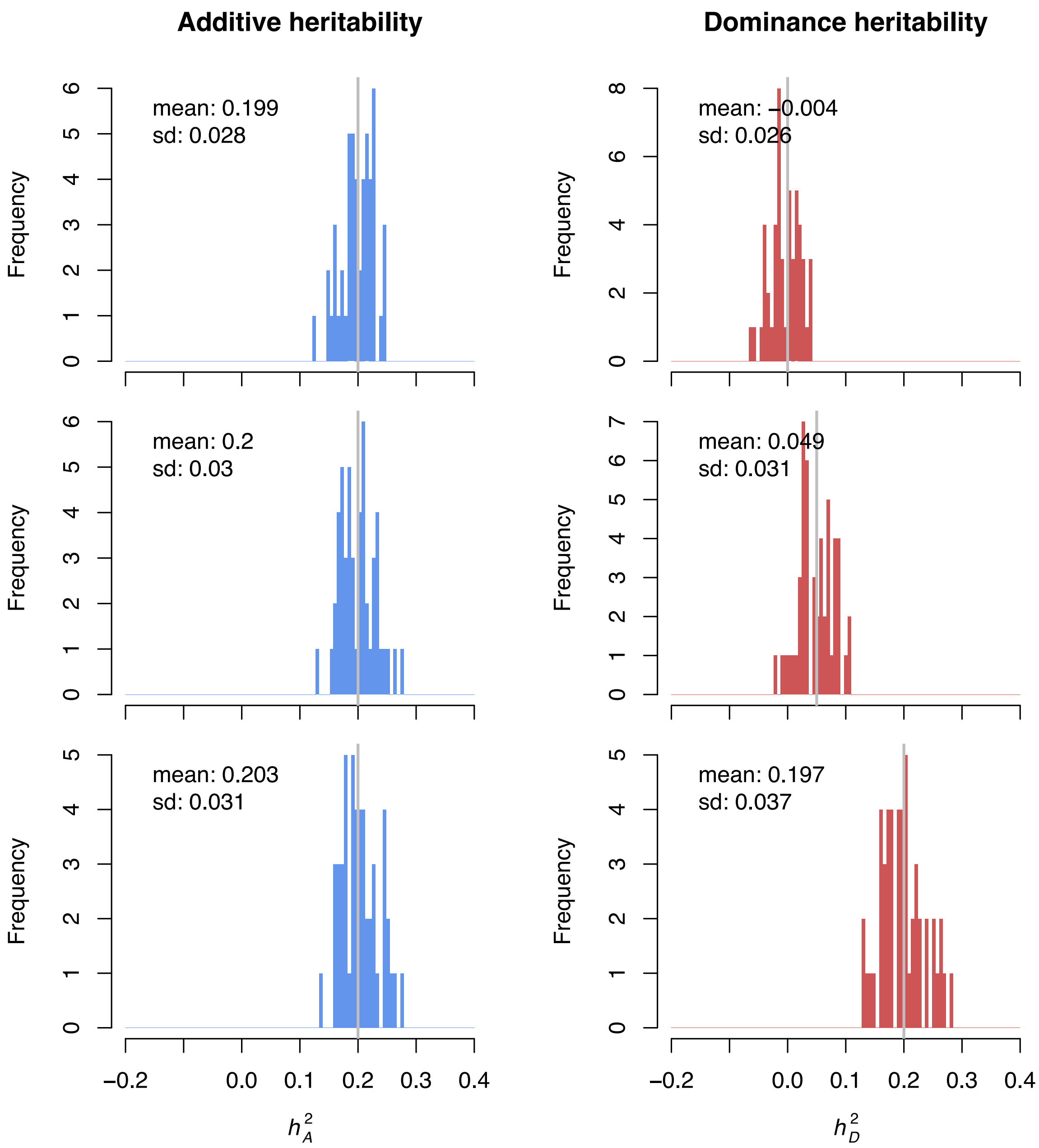


**Figure** S23: Msprime simulation study: Case-control phenotype with 6% prevalence, no case ascertainment, 10% SNPs causal. Phenotypes are simulated under a spike and slab architecture with 10% causal sites. Histograms of additive and dominance heritability estimates across the 50 independent phenotype simulations are shown in the left and right columns respectively. The grey line in the true underlying heritability, mean and standard deviations of the point estimates across runs are inset in each panel.

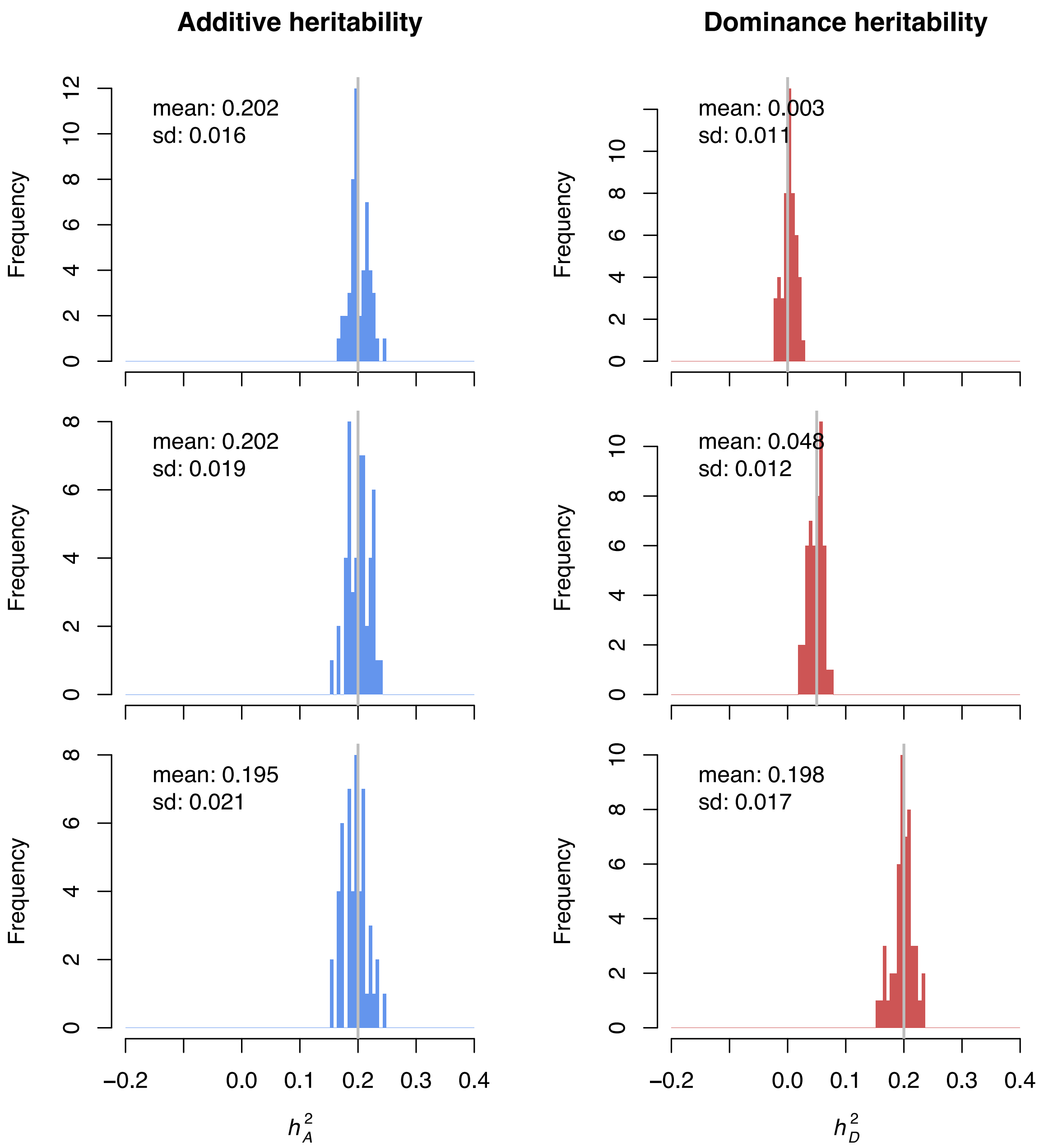


**Figure** S24: Msprime simulation study: Case-control phenotype with $6\%$ prevalence, $18\%$ ascertained sample prevalence, all SNPs causal. Phenotypes are simulated under an infinitesimal architecture. Histograms of additive and dominance heritability estimates across the $50$ independent phenotype simulations are shown in the left and right columns respectively. The grey line in the true underlying heritability, mean and standard deviations of the point estimates across runs are inset in each panel.

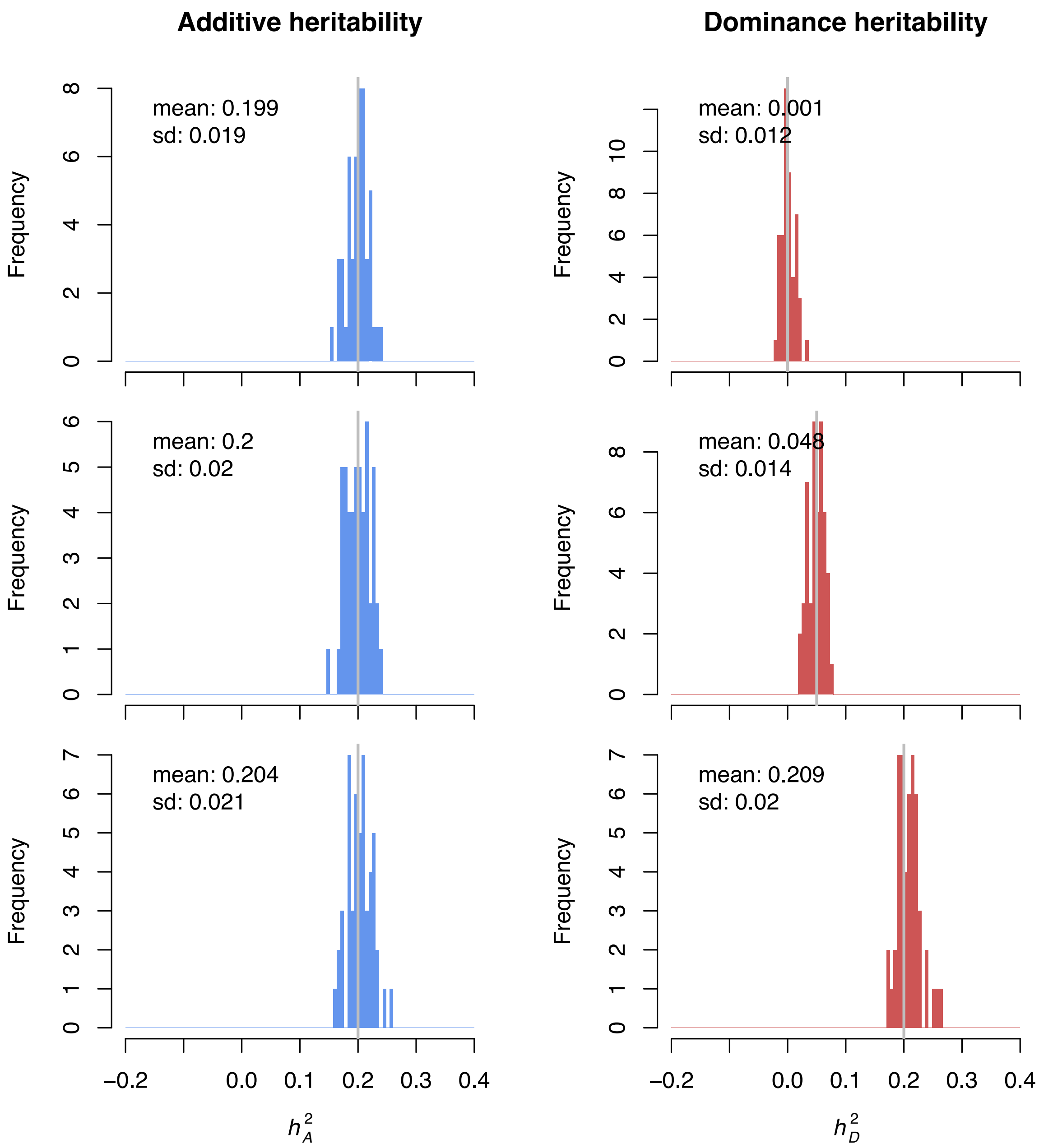


**Figure** S25: Msprime simulation study: Case-control phenotype with 6% prevalence, 18% ascertained sample prevalence, 10% SNPs causal. Phenotypes are simulated under a spike and slab architecture with 10% causal sites. Histograms of additive and dominance heritability estimates across the 50 independent phenotype simulations are shown in the left and right columns respectively. The grey line in the true underlying heritability, mean and standard deviations of the point estimates across runs are inset in each panel.

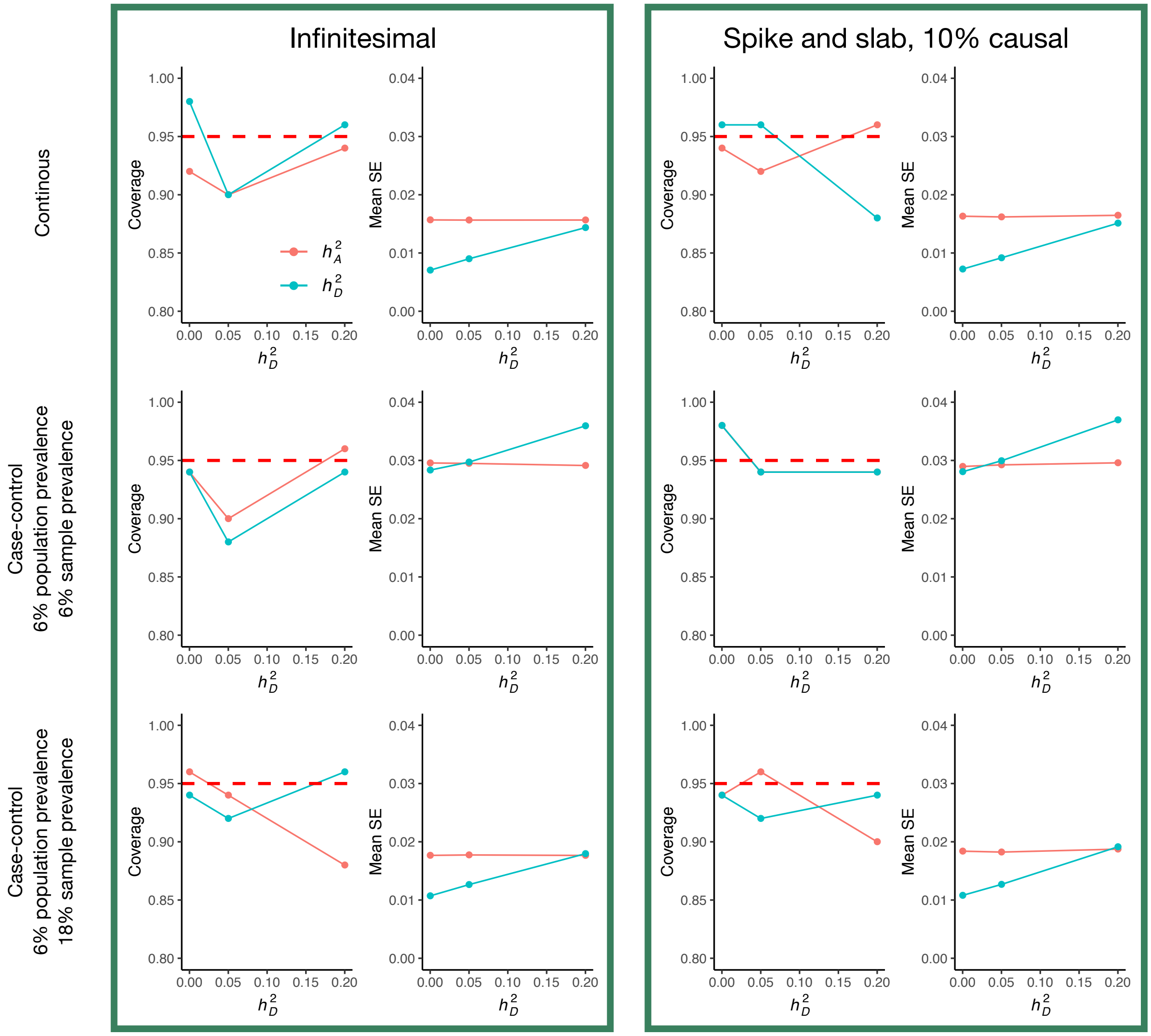


**Figure** S26: Msprime simulation study: Coverage and standard errors of heritability point estimates. The first and second column display the $95\%$ coverage (proportion of the $50$ runs with the true heritability lying within the $95\%$ confidence interval) and mean standard error respectively for the infinitesimal model. Columns three and four show the corresponding metrics for a spike and slab architecture with $10\%$ causal SNPs. Collected phenotypes change as we move through the rows according the label to the left of the panels.

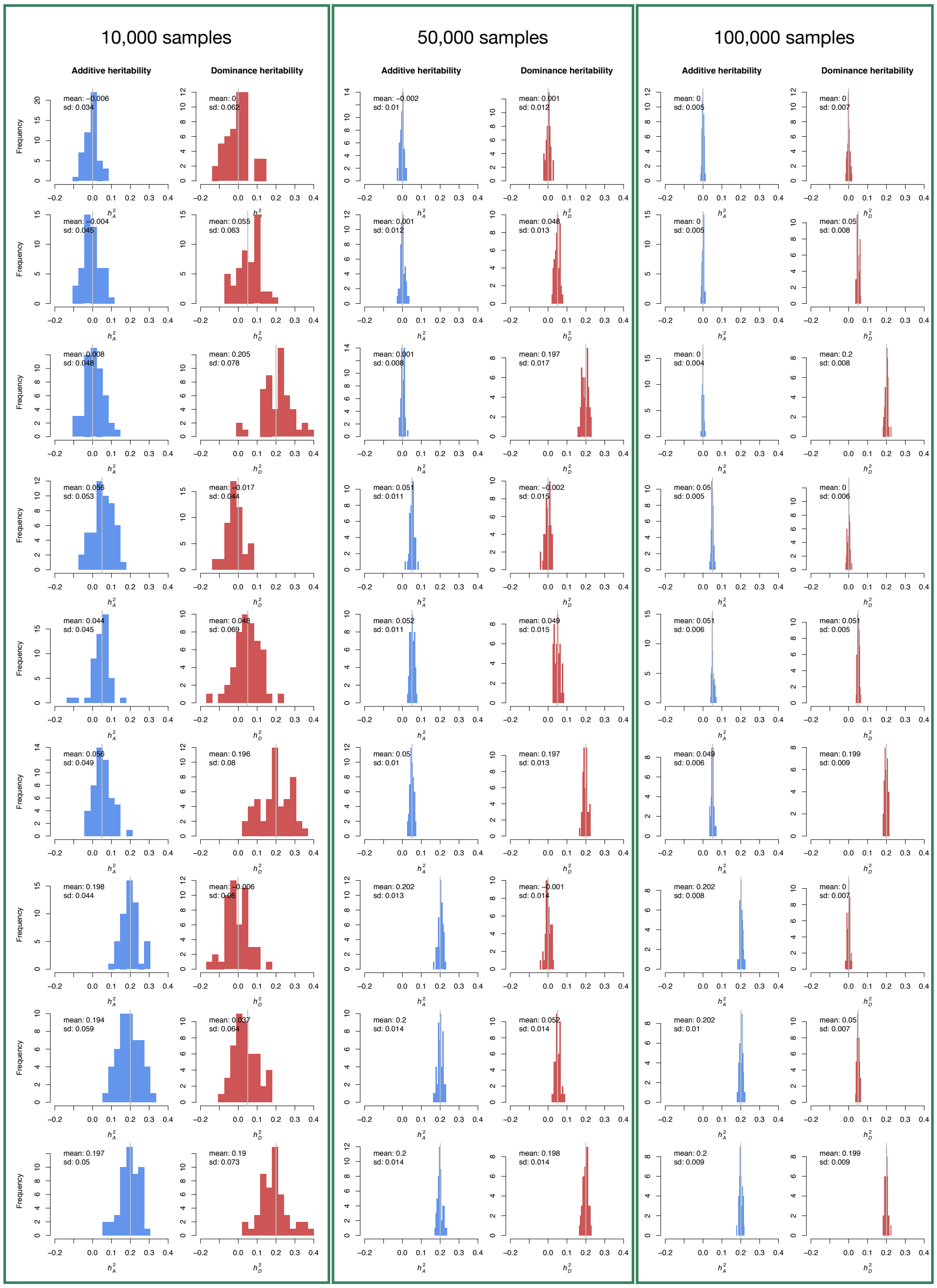


**Figure** S27: UK Biobank simulation study: Distributions of heritability point estimates. Histograms of additive and dominance heritability estimates across the 50 independent phenotype simulations are shown in blue and red respectively. The grey line in the true underlying heritability, mean and standard deviations of the point estimates across runs are inset in each panel. Columns outlined in green delineate runs with $10,000$ samples, $50,000$ samples, and $100,000$ samples respectively.

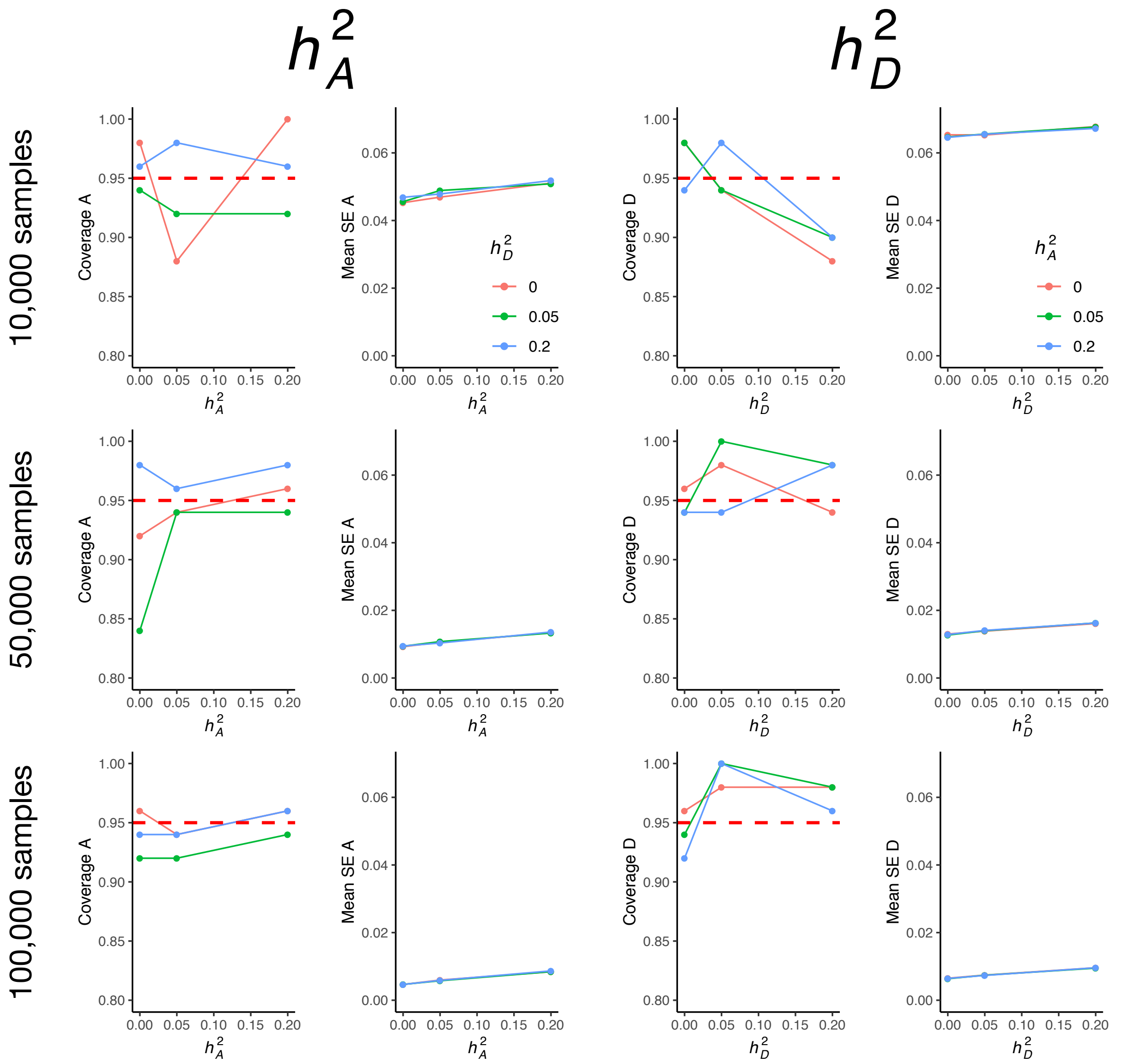


**Figure** S28: UKBB simulation study: Coverage and standard errors of heritability point estimates. The first and second column display the $95\%$ coverage (proportion of the 50 runs with the true heritability lying within the $95\%$ confidence interval) and mean standard error respectively for the additive heritability. Columns three and four show the corresponding metrics for dominance heritability estimates. Sample size of our simulations increases moving through the rows, from $10,000$ to $100,000$. Line colourings are coloured according to the keys, specific the the first two and second two columns respectively.

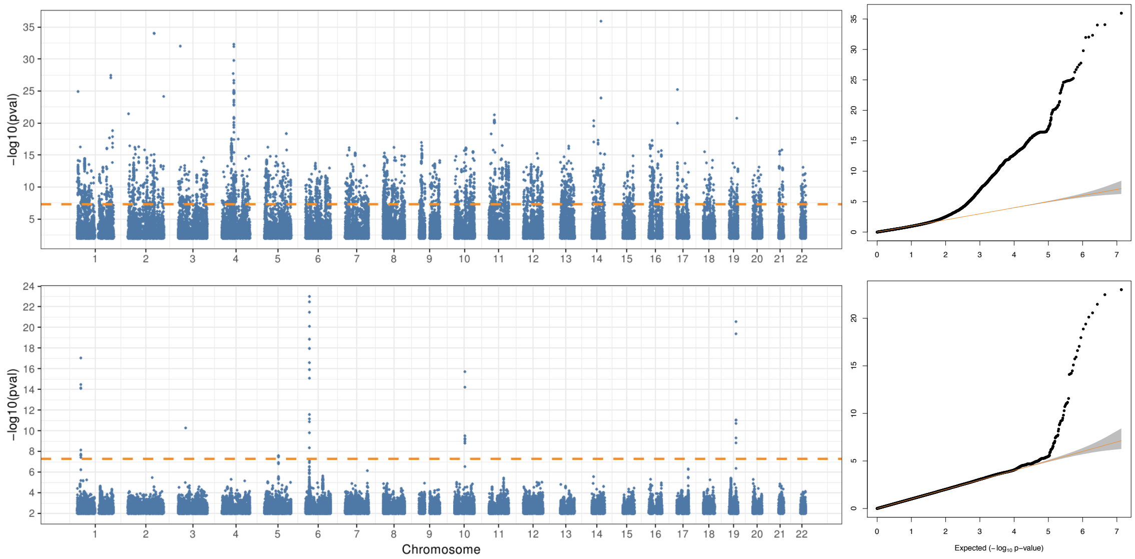


**Figure** S29: Impact of IRNT, high light scatter reticulocyte percentage. Here we show the most extreme effect of the IRNT transformation on the observed dominance signal across the genome. The first and second rows display the dominance Manhattan and QQ-plots for high light scatter reticulocyte percentage (UK Biobank phenotype ID 30290) before and after inverse-rank normalisation. The dotted orange line in the Manhattan plots indicate genome wide significance ($P = 5 \times 10^{-8}$).

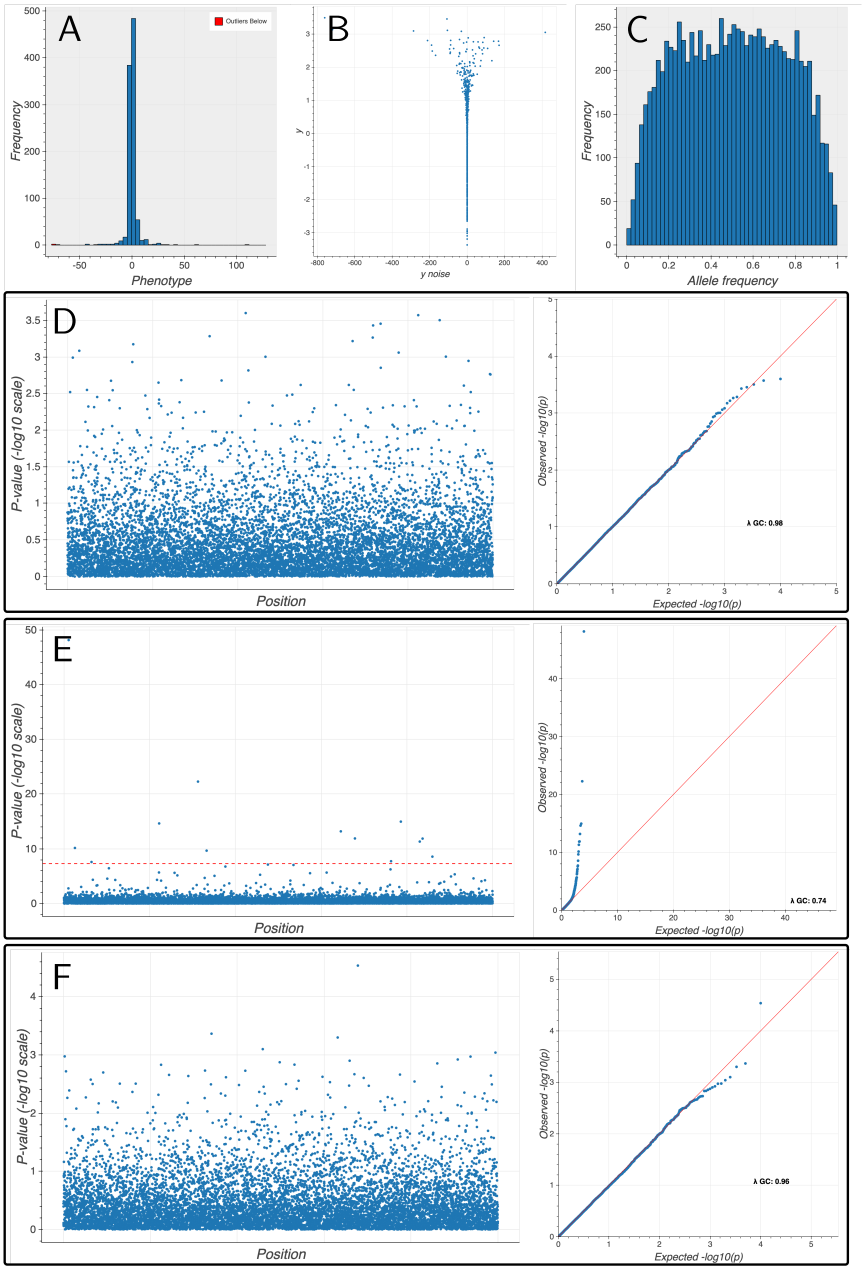


**Figure** S30: Results of simulation study with a heteroskedastic noise term under the infinitesimal model with 10,000 sites and 1,000 individuals. Genotypes are simulated under the Balding-Nichols model with two populations and $F_{ST}$ of 0.1 to the ancestral population in each. Simulation of a heteroskedastic noise term defined in (6) with $\alpha = 10$. A displays the distribution of the GWASed phenotype. B shows the relationship between true $y$ and the heteroskedastic measured $y$. C displays the allele frequency distribution of the genotype data. D-F display Manhattan (though sites here are unlinked) and associated QQ plots for the additive, dominance before, and dominance after inverse rank normalisation respectively.

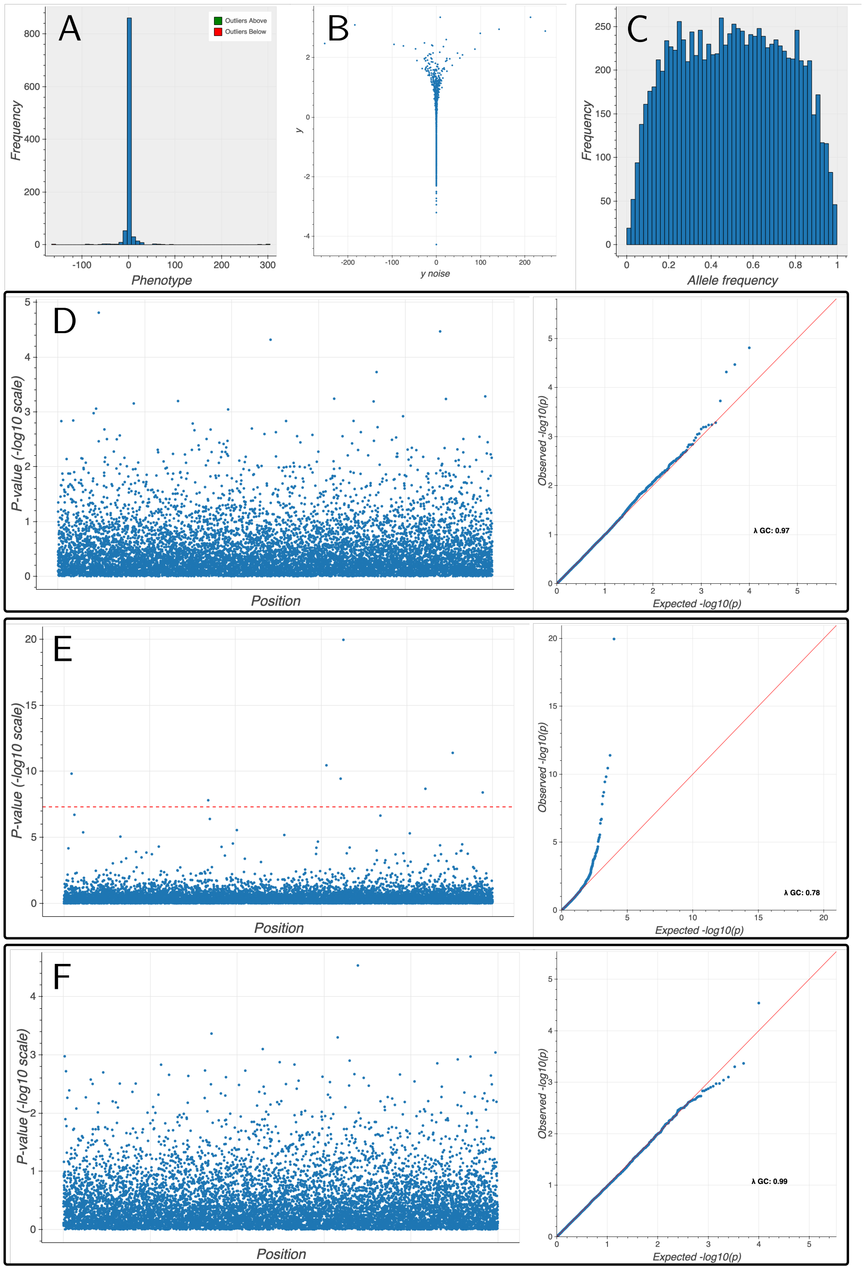


**Figure** S31: Results of simulation study with a heteroskedastic noise term under the spike and slab model; $\pi = 0.01$, with 10,000 sites and 1,000 individuals. Genotypes are simulated under the Balding-Nichols model with two populations and $F_{ST}$ of 0.1 to the ancestral population in each. Simulation of a heteroskedastic noise term defined in (6) with $\alpha = 10$. A displays the distribution of the GWASed phenotype. B shows the relationship between true $y$ and the heteroskedastic measured $y$. C displays the allele frequency distribution of the genotype data. D-F display Manhattan (though sites here are unlinked) and associated QQ plots for the additive, dominance before, and dominance after inverse rank normalisation respectively.

| Phenotype | Gene | Variant | rsid | Exonic function | $\widehat{\beta}_A$ | $\log_{10}(p_A)$ | Add PIP | $\widehat{\beta}_D$ | $\log_{10}(p_D)$ | Dom PIP |
|---|---|---|---|---|---|---|---|---|---|---|
| Natural hair colour: Blonde | TCHH | 1:152083325:A:T | rs11803731 | non-syn SNV | -0.00221 | -1.66 | $1.83 \times 10^{-5}$ | -0.00355 | -10.8 | 0.218 |
| C-reactive protein | IL6R | 1:154426970:A:C | rs2228145 | non-syn SNV | -0.0935 | $< -308$ | 0.00525 | 0.00958 | -7.97 | 0.0889 |
| Mean reticulocyte volume | SPTA1 | 1:158582646:T:C | rs2251969 | syn SNV | 0.0401 | -61.8 | $1.5 \times 10^{-5}$ | -0.00666 | -4.09 | 0.113 |
| Mean sphered cell volume | SPTA1 | 1:158582646:T:C | rs2251969 | syn SNV | 0.0522 | -104 | $3.16 \times 10^{-5}$ | -0.00722 | -4.7 | 0.104 |
| Glycated haemoglobin | SPTA1 | 1:158582646:T:C | rs2251969 | syn SNV | 0.0457 | -86.5 | $3.22 \times 10^{-5}$ | -0.00509 | -2.75 | 0.0995 |
| Mean reticulocyte volume | SPTA1 | 1:158584091:A:G | rs952094 | non-syn SNV | 0.0402 | -62.2 | $1.49 \times 10^{-5}$ | -0.00666 | -4.08 | 0.0911 |
| Mean sphered cell volume | SPTA1 | 1:158584091:A:G | rs952094 | non-syn SNV | 0.0524 | -105 | $3.3 \times 10^{-5}$ | -0.00724 | -4.72 | 0.0979 |
| Glycated haemoglobin | SPTA1 | 1:158584091:A:G | rs952094 | non-syn SNV | 0.0454 | -85.7 | $2.94 \times 10^{-5}$ | -0.00531 | -2.96 | 0.226 |
| Glycated haemoglobin | SPTA1 | 1:158626378:C:T | rs857691 | syn SNV | 0.076 | -178 | $1.16 \times 10^{-5}$ | 0.0113 | -10.6 | 0.239 |
| Mean platelet (thrombocyte) volume | TRIM58 | 1:248039294:G:A | rs1339847 | non-syn SNV | -0.102 | -156 | $5.26 \times 10^{-5}$ | -0.0125 | -13 | 0.651 |
| Ever had eye surgery | ATL2 | 2:38537579:T:C | rs34873284 | non-syn SNV | -0.00114 | -0.16 | 0 | -0.00495 | -7.9 | 0.888 |
| Factors influencing health status | FAM178B | 2:97559759:G:A | rs11677797 | syn SNV | 0.000137 | -0.0489 | 0 | -0.00318 | -8.01 | 0.11 |
| Hearing difficulty/problems: Yes | ILDR1 | 3:121712051:A:C | rs2877561 | syn SNV | -0.00645 | -7.43 | 0.0458 | -0.00525 | -11.8 | 0.231 |
| Hayfever, allergic rhinitis or eczema | TLR1 | 4:38798935:C:T | rs5743614 | syn SNV | -0.024 | -82.1 | $1.66 \times 10^{-6}$ | -0.00374 | -7.24 | 0.0641 |
| Hearing difficulty/problems: Yes | CHMP4C | 8:82670771:G:A | rs35094336 | non-syn SNV | 0.00985 | -5.91 | 0.109 | -0.00453 | -9.09 | 0.134 |
| Alkaline phosphatase | ABO | 9:136131461:G:A | rs8176741 | unknown | 0.0473 | -21.3 | $3.22 \times 10^{-5}$ | 0.00253 | -0.695 | 0.988 |
| Air pollution (pm2.5); 2010 | KNDC1 | 10:134999602:G:C | rs3008393 | syn SNV | -0.000343 | -0.0326 | 0 | -0.00973 | -7.57 | 0.158 |
| Eosinophill percentage | PRG3 | 11:57146225:A:G | rs34108746 | non-syn SNV | -0.0647 | -47.6 | 1 | 0.0439 | -153 | 1 |
| Natural hair colour: Black | TYR | 11:89017961:G:A | rs1126809 | non-syn SNV | -0.00339 | -10.3 | 0.501 | 0.0022 | -10.2 | 0.952 |
| Alanine aminotransferase | PANX1 | 11:93862493:A:C | rs1138800 | non-syn SNV | 0.00497 | -1.5 | $1.07 \times 10^{-5}$ | 0.0104 | -10.4 | 0.389 |
| Yogurt/ice-cream consumers | CLEC12A | 12:10137557:A:C | rs479499 | non-syn SNV | -0.00447 | -0.803 | 0 | -0.0115 | -7.04 | 0.0589 |
| Natural hair colour: Blonde | MC1R | 16:89986117:C:T | rs1805007 | non-syn SNV | 0.0295 | -129 | $2.76 \times 10^{-7}$ | 0.00874 | -63.9 | 0.999 |
| Natural hair colour: Light brown | MC1R | 16:89986117:C:T | rs1805007 | non-syn SNV | -0.028 | -49.3 | 0.288 | 0.0291 | -292 | 1 |
| Natural hair colour: Light brown | MC1R | 16:89986144:C:T | rs1805008 | non-syn SNV | -0.00275 | -0.752 | $1.35 \times 10^{-7}$ | 0.00842 | -24.4 | 0.457 |
| Natural hair colour: Light brown | MC1R | 16:89986546:G:C | rs1805009 | non-syn SNV | -0.0277 | -12.8 | 0.838 | 0.0086 | -17.6 | 0.999 |
| Natural hair colour: Light brown | CENPBD1 | 16:90037828:G:A | rs4785755 | non-syn SNV | -0.00282 | -1.59 | $1.27 \times 10^{-7}$ | -0.000599 | -0.334 | 0.163 |
| Worry too long after embarrassment | DPH1 | 17:1943880:C:G | rs35394823 | non-syn SNV | 0.00393 | -1.25 | $1.87 \times 10^{-6}$ | 0.00422 | -6.37 | 0.0619 |
| Natural hair colour: Light brown | TSPAN10 | 17:79612397:A:G | rs6420484 | unknown | -0.0185 | -52.1 | 0.0167 | -0.00928 | -29.1 | 0.0533 |
| Natural hair colour: Dark brown | TSPAN10 | 17:79612397:A:G | rs6420484 | unknown | 0.0203 | -64.8 | 0.000139 | 0.0128 | -56 | 0.0762 |
| logMAR, final (right) | TSPAN10 | 17:79612397:A:G | rs6420484 | unknown | -0.0443 | -17.5 | 0.0434 | -0.0221 | -9.81 | 0.134 |
| logMAR, final (left) | TSPAN10 | 17:79612397:A:G | rs6420484 | unknown | -0.0537 | -25.3 | 0.0422 | -0.0271 | -14.4 | 0.193 |
| Alanine aminotransferase | LRRC45 | 17:79986156:C:T | rs72861736 | non-syn SNV | 0.0129 | -3.48 | 0.00252 | -0.01 | -10.3 | 0.121 |
| Apoliprotein B | TM6SF2 | 19:19379549:C:T | rs58542926 | non-syn SNV | -0.0956 | -99.6 | 0.541 | 0.0275 | -60.4 | 0.969 |
| Cholesterol | TM6SF2 | 19:19379549:C:T | rs58542926 | non-syn SNV | -0.116 | -154 | 0.846 | 0.021 | -37.4 | 0.966 |
| LDL direct | TM6SF2 | 19:19379549:C:T | rs58542926 | non-syn SNV | -0.108 | -129 | 0.617 | 0.0228 | -42.5 | 0.971 |
| SHBG | TM6SF2 | 19:19379549:C:T | rs58542926 | non-syn SNV | -0.00317 | -0.337 | 0.000234 | -0.00975 | -9.14 | 0.222 |
| Self-reported: high cholesterol | APOE | 19:45412079:C:T | rs7412 | non-syn SNV | -0.037 | -158 | 0.0121 | -0.00599 | -28.4 | 0.916 |
| Medication code: simvastatin | APOE | 19:45412079:C:T | rs7412 | non-syn SNV | -0.0248 | -76.8 | 0.0304 | -0.003 | -8.24 | 0.634 |
| Medication code: atorvastatin | APOE | 19:45412079:C:T | rs7412 | non-syn SNV | -0.00933 | -36.5 | 0.374 | -0.0018 | -9.65 | 0.77 |
| Mean corpuscular haemoglobin | APOE | 19:45412079:C:T | rs7412 | non-syn SNV | -0.0221 | -6.46 | 0.0661 | 0.0102 | -8.84 | 0.574 |
| Lipoprotein A | APOE | 19:45412079:C:T | rs7412 | non-syn SNV | -0.108 | -103 | 0.0673 | 0.0131 | -10.6 | 0.111 |
| Cholesterol | FUT2 | 19:49206417:A:G | rs492602 | syn SNV | 0.0289 | -34.7 | 0.016 | -0.0128 | -14.3 | 0.0996 |
| Gamma glutamyltransferase | FUT2 | 19:49206417:A:G | rs492602 | syn SNV | 0.0448 | -89.1 | 0.0259 | -0.0113 | -12.3 | 0.0657 |
| Glycated haemoglobin | FUT2 | 19:49206417:A:G | rs492602 | syn SNV | -0.00722 | -2.76 | 0.00125 | 0.00888 | -7.34 | 0.198 |
| LDL direct | FUT2 | 19:49206417:A:G | rs492602 | syn SNV | 0.0286 | -33 | 0.092 | -0.012 | -12.2 | 0.101 |
| Total protein | FUT2 | 19:49206417:A:G | rs492602 | syn SNV | 0.0375 | -49.3 | 0.0464 | -0.0109 | -9.07 | 0.162 |

| | | | | | | | | | | |
|---|---|---|---|---|---|---|---|---|---|---|
| Alkaline phosphatase | FUT2 | 19:49206462:C:T | rs681343 | syn SNV | -0.119 | $< -308$ | 0.167 | 0.0329 | -88.2 | 0.227 |
| Apolipoprotein B | FUT2 | 19:49206462:C:T | rs681343 | syn SNV | 0.029 | -33.4 | 0.161 | -0.0113 | -10.7 | 0.207 |
| Cholesterol | FUT2 | 19:49206462:C:T | rs681343 | syn SNV | 0.029 | -34.9 | 0.0217 | -0.013 | -14.7 | 0.223 |
| Gamma glutamyltransferase | FUT2 | 19:49206462:C:T | rs681343 | syn SNV | 0.0447 | -89 | 0.0244 | -0.0113 | -12.2 | 0.0582 |
| Glycated haemoglobin | FUT2 | 19:49206462:C:T | rs681343 | syn SNV | -0.00714 | -2.7 | 0.00113 | 0.00887 | -7.32 | 0.19 |
| LDL direct | FUT2 | 19:49206462:C:T | rs681343 | syn SNV | 0.0287 | -33.1 | 0.116 | -0.0122 | -12.6 | 0.23 |
| Total protein | FUT2 | 19:49206462:C:T | rs681343 | syn SNV | 0.0373 | -48.8 | 0.0174 | -0.0109 | -9.09 | 0.169 |
| Apolipoprotein B | FUT2 | 19:49206674:G:A | rs601338 | stopgain | 0.0289 | -33.1 | 0.0738 | -0.0113 | -10.7 | 0.213 |
| Cholesterol | FUT2 | 19:49206674:G:A | rs601338 | stopgain | 0.0289 | -34.6 | 0.0128 | -0.013 | -14.7 | 0.203 |
| Gamma glutamyltransferase | FUT2 | 19:49206674:G:A | rs601338 | stopgain | 0.0449 | -89.4 | 0.0594 | -0.0114 | -12.3 | 0.0798 |
| Glycated haemoglobin | FUT2 | 19:49206674:G:A | rs601338 | stopgain | -0.00709 | -2.67 | 0.00107 | 0.00883 | -7.26 | 0.167 |
| LDL direct | FUT2 | 19:49206674:G:A | rs601338 | stopgain | 0.0285 | -32.8 | 0.0659 | -0.0122 | -12.6 | 0.228 |
| Total protein | FUT2 | 19:49206674:G:A | rs601338 | stopgain | 0.0373 | -48.9 | 0.0184 | -0.0109 | -9.13 | 0.183 |
| Red blood cell (erythrocyte) distribution width | ITPA | 20:3193842:C:A | rs1127354 | non-syn SNV | -0.0177 | -4.03 | 0.00848 | 0.0112 | -10.4 | 0.245 |
| Mean sphered cell volume | ITPA | 20:3193842:C:A | rs1127354 | non-syn SNV | -0.00551 | -0.643 | 0.000181 | 0.0116 | -11 | 0.215 |
| Mean corpuscular volume | TMPRSS6 | 22:37469591:G:A | rs4820268 | syn SNV | 0.125 | $< -308$ | $3.22 \times 10^{-10}$ | 0.0128 | -13.7 | 0.306 |
| Mean corpuscular haemoglobin | TMPRSS6 | 22:37469591:G:A | rs4820268 | syn SNV | 0.145 | $< -308$ | $1.66 \times 10^{-8}$ | 0.0162 | -21.4 | 0.461 |
| Alanine aminotransferase | PNPLA3 | 22:44324727:C:G | rs738409 | non-syn SNV | 0.0992 | -293 | 0.415 | -0.0175 | -27.9 | 0.437 |
| Alanine aminotransferase | PNPLA3 | 22:44324730:C:T | rs738408 | syn SNV | 0.0991 | -293 | 0.161 | -0.0175 | -27.9 | 0.46 |

**Table** S2: Fine-mapped exonic variants with larger dominance posterior probability. Ordered by position in the genome, $p_A$ and $p_D$ denote the $P$-value associated to the marginal additive and dominance effect size being non-zero respectively. Add PIP and dom PIP are the additive and dominance posterior inclusion probability respectively.

| Phenotype | Prop. sites causal | $h_A^2$ | $h_D^2$ | Mean $\widehat{h}_A^2$ | Mean $\widehat{h}_A^2$ SE | $\widehat{h}_A^2$ 95% CI coverage | Mean $\widehat{h}_D^2$ | Mean $\widehat{h}_D^2$ SE | $\widehat{h}_D^2$ 95% CI coverage |
|---|---|---|---|---|---|---|---|---|---|
| Continuous | 100% | 0.20 | 0.00 | 0.194 | 0.0157 | 0.92 | 0.001 | 0.0071 | 0.98 |
| | | 0.20 | 0.05 | 0.197 | 0.0157 | 0.90 | 0.051 | 0.0090 | 0.90 |
| | | 0.20 | 0.20 | 0.199 | 0.0157 | 0.94 | 0.199 | 0.0144 | 0.96 |
| | 10% | 0.20 | 0.00 | 0.198 | 0.0163 | 0.94 | 0.001 | 0.0072 | 0.96 |
| | | 0.20 | 0.05 | 0.200 | 0.0162 | 0.92 | 0.051 | 0.0092 | 0.96 |
| | | 0.20 | 0.20 | 0.210 | 0.0165 | 0.96 | 0.202 | 0.0151 | 0.88 |
| Case control: 6% prevalence, no ascertainment | 100% | 0.20 | 0.00 | 0.193 | 0.0296 | 0.94 | -0.007 | 0.0283 | 0.94 |
| | | 0.20 | 0.05 | 0.207 | 0.0295 | 0.90 | 0.048 | 0.0297 | 0.88 |
| | | 0.20 | 0.20 | 0.201 | 0.0291 | 0.96 | 0.207 | 0.0359 | 0.94 |
| | 10% | 0.20 | 0.00 | 0.199 | 0.0290 | 0.98 | -0.004 | 0.0281 | 0.98 |
| | | 0.20 | 0.05 | 0.200 | 0.0292 | 0.94 | 0.049 | 0.0299 | 0.94 |
| | | 0.20 | 0.20 | 0.203 | 0.0296 | 0.94 | 0.197 | 0.0370 | 0.94 |
| Case control: 6% prevalence, 18% sample prevalence | 100% | 0.20 | 0.00 | 0.202 | 0.0177 | 0.96 | 0.003 | 0.0107 | 0.94 |
| | | 0.20 | 0.05 | 0.202 | 0.0177 | 0.94 | 0.048 | 0.0126 | 0.92 |
| | | 0.20 | 0.20 | 0.195 | 0.0177 | 0.88 | 0.198 | 0.0180 | 0.96 |
| | 10% | 0.20 | 0.00 | 0.199 | 0.0184 | 0.94 | 0.001 | 0.0108 | 0.94 |
| | | 0.20 | 0.05 | 0.200 | 0.0182 | 0.96 | 0.048 | 0.0127 | 0.92 |
| | | 0.20 | 0.20 | 0.204 | 0.0187 | 0.90 | 0.209 | 0.0191 | 0.94 |

**Table** S3: Summary of msprime simulation results. Means of point estimates and standard errors are taken across the $50$ independent phenotype simulations for each $(h_A^2, h_D^2)$ pair. The coverage is the proportion of the $50$ runs that the true underlying heritability lies within the $95\%$ confidence interval as defined by the estimated standard error.

#### Evaluation of additive and dominance heritability

We first determine the additive and dominance LD-scores to use for the heritability analyses. We filter to HapMap3 SNPs (*63*) and evaluate additive and dominance LD-scores within a 1cM window. These additive and dominance LD-scores were used in the regressions and as weights in the respective heritability analyses. We then evaluate additive and dominance heritability estimates by looping over phenotype dominance summary statistics files.

#### LD-score dominance extension

In order to estimate the contribution of within locus non-additive effects across the genome to phenotypic variation, we extend the infinitesimal model underlying LD-score regression to in-

| $h^2_A$ | $h^2_D$ | Mean $h^2_A$ | Mean $h^2_A$ SE | $h^2_A$ 95% CI coverage | Mean $h^2_D$ | Mean $h^2_D$ SE | $h^2_D$ 95% CI coverage |
|---|---|---|---|---|---|---|---|
| 0.00 | 0.00 | -0.002 | 0.0093 | 0.92 | 0.001 | 0.0130 | 0.96 |
| 0.00 | 0.05 | 0.001 | 0.0094 | 0.94 | 0.048 | 0.0138 | 0.98 |
| 0.00 | 0.20 | 0.001 | 0.0095 | 0.96 | 0.197 | 0.0161 | 0.94 |
| 0.05 | 0.00 | 0.051 | 0.0105 | 0.84 | -0.002 | 0.0127 | 0.94 |
| 0.05 | 0.05 | 0.052 | 0.0108 | 0.94 | 0.049 | 0.0140 | 1.00 |
| 0.05 | 0.20 | 0.050 | 0.0104 | 0.94 | 0.197 | 0.0163 | 0.98 |
| 0.20 | 0.00 | 0.202 | 0.0134 | 0.98 | -0.001 | 0.0128 | 0.94 |
| 0.20 | 0.05 | 0.200 | 0.0133 | 0.96 | 0.052 | 0.0141 | 0.94 |
| 0.20 | 0.20 | 0.200 | 0.0136 | 0.98 | 0.198 | 0.0163 | 0.98 |

**Table** S4: Summary of UK Biobank simulation results. We summarise the results for our sample size $50,000$ simulations. Means of point estimates and standard errors are taken across the $50$ independent phenotype simulations for each $(h^2_A, h^2_D)$ pair. The coverage is the proportion of the $50$ runs that the true underlying heritability lies within the $95\%$ confidence interval as defined by the estimated standard error.

| Phenotype | mean $\chi^2_A$ | $\lambda^A_{GC}$ | int $A$ | SE int $A$ | $P$ int $A$ | $h^2_A$ | SE $h^2_A$ | $P$ $h^2_A$ |
|---|---|---|---|---|---|---|---|---|
| $h$ | 2.77 | 1.84 | 1.10 | 0.0334 | 0.00157 | 0.493 | 0.0266 | $4.40 \times 10^{-77}$ |
| $\mathrm{relu}\,(h)$ | 2.34 | 1.70 | 1.06 | 0.0254 | 0.00558 | 0.381 | 0.0207 | 4.68-76 |
| $h^2$ | 1.01 | 1.02 | 1.00 | 0.00643 | 0.508 | 0.00439 | 0.00278 | 0.0568 |
| $\exp\,(h)$ | 1.96 | 1.54 | 1.05 | 0.0201 | 0.00546 | 0.270 | 0.0159 | $3.86 \times 10^{-65}$ |
| $\exp\left(-h^2\right)$ | 1.01 | 1.02 | 0.999 | 0.00677 | 0.536 | 0.00413 | 0.0031 | 0.0913 |
| $\exp\left(-100h^2\right)$ | 0.990 | 0.992 | 0.990 | 0.00608 | 0.944 | $3.4 \times 10^{-6}$ | 0.00267 | 0.499 |

**Table** S5: Additive heritability statistics for non-linear re-scalings of height. $h$ is normalised height, we then apply a collection of non-linear rescalings and examine the impact on additive heritability estimates.

| Phenotype | mean $\chi^2_D$ | $\lambda^D_{GC}$ | int $D$ | SE int $D$ | $p$ int $D$ | $h^2_D$ | SE $h^2_D$ | $p$ $h^2_D$ |
|---|---|---|---|---|---|---|---|---|
| $h$ | 1.00 | 1.00 | 1.00 | 0.00387 | 0.279 | $-0.000141$ | 0.00391 | 0.514 |
| $\mathrm{relu}\,(h)$ | 1.00 | 1.00 | 0.999 | 0.00407 | 0.626 | 0.00232 | 0.00382 | 0.272 |
| $h^2$ | 0.999 | 0.967 | 0.979 | 0.00576 | 1.00 | 0.0123 | 0.00539 | 0.0110 |
| $\exp\,(h)$ | 1.00 | 0.999 | 0.997 | 0.00439 | 0.721 | 0.00194 | 0.00368 | 0.299 |
| $\exp\left(-h^2\right)$ | 0.999 | 1.00 | 0.994 | 0.00363 | 0.957 | 0.00327 | 0.00340 | 0.168 |
| $\exp\left(-100h^2\right)$ | 1.01 | 1.01 | 0.996 | 0.00395 | 0.855 | 0.00603 | 0.00356 | 0.0449 |

**Table** S6: Dominance heritability statistics for non-linear re-scalings of height. $h$ is normalised height, we then apply a collection of non-linear rescalings and examine the impact on additive heritability estimates.

| Phenotype | $h^2_A$ | SE $h^2_A$ | $P$ $h^2_A$ |
|---|---|---|---|
| Leg fat percentage | 0.224 | 0.00663 | 5.39e-250 |
| Qualifications: College or University degree | 0.295 | 0.00877 | 2.45e-248 |
| Body mass index (BMI) | 0.252 | 0.00802 | 2.6e-217 |
| Leg fat mass | 0.237 | 0.00776 | 7.58e-206 |
| Impedance of arm | 0.24 | 0.00797 | 1.37e-199 |
| Qualifications: None of the above | 0.236 | 0.00794 | 7.95e-195 |
| Waist circumference | 0.206 | 0.00698 | 1.18e-192 |
| Duration to first press of snap-button in each round | 0.0864 | 0.00292 | 1.64e-192 |
| Qualifications: A levels/AS levels or equivalent | 0.171 | 0.00599 | 2.01e-179 |
| Age first had sexual intercourse | 0.162 | 0.00571 | 3.98e-178 |
| Mean time to correctly identify matches | 0.084 | 0.00302 | 8.28e-171 |
| Weight | 0.27 | 0.00992 | 2.88e-163 |
| Hand grip strength (left) | 0.117 | 0.00435 | 5.84e-161 |
| Sodium in urine | 0.0741 | 0.00283 | 1.88e-151 |
| Smoking status: Never | 0.156 | 0.0061 | 3.2e-145 |
| Hip circumference | 0.223 | 0.00874 | 7.6e-144 |
| Systolic blood pressure, automated reading | 0.149 | 0.00603 | 2.06e-135 |
| Leg predicted mass | 0.282 | 0.0115 | 1.93e-133 |
| Leg fat-free mass | 0.283 | 0.0116 | 9.23e-133 |
| Basal metabolic rate | 0.299 | 0.0124 | 2.56e-128 |
| Age at first live birth | 0.18 | 0.00748 | 2.64e-128 |
| Bread type: White | 0.111 | 0.00462 | 2.1e-127 |
| Vascular/heart problems diagnosed by doctor: None of the above | 0.217 | 0.00904 | 3.44e-127 |
| Snoring | 0.0995 | 0.00419 | 9.25e-125 |
| Creatinine (enzymatic) in urine | 0.0687 | 0.00292 | 2.94e-122 |
| Diastolic blood pressure, automated reading | 0.14 | 0.00601 | 1.28e-119 |
| Ever smoked | 0.123 | 0.00535 | 1.52e-117 |
| Coffee type: Ground coffee (include espresso, filter etc) | 0.132 | 0.00574 | 1.4e-116 |
| Non-cancer illness code, self-reported: hypertension | 0.225 | 0.00986 | 5.26e-116 |
| Forced expiratory volume in 1-second (FEV1) | 0.195 | 0.00855 | 7.17e-115 |
| Time to complete round | 0.0902 | 0.00404 | 2.1e-110 |
| Fluid intelligence score | 0.23 | 0.0103 | 9.25e-110 |
| Mood swings | 0.113 | 0.00512 | 2.09e-108 |
| Forced vital capacity (FVC) | 0.214 | 0.00976 | 1.5e-106 |
| Fed-up feelings | 0.116 | 0.00533 | 2.21e-105 |
| Pain type(s) experienced in last month: None of the above | 0.0886 | 0.00417 | 1.32e-100 |
| Long-standing illness, disability or infirmity | 0.0912 | 0.00436 | 1.42e-97 |
| Alcohol usually taken with meals | 0.173 | 0.00829 | 5.57e-97 |
| Hearing difficulty/problems with background noise | 0.086 | 0.00413 | 1.06e-96 |
| Number of incorrect matches in round | 0.0539 | 0.00263 | 2.16e-93 |

**Table** S7: The top 40 most significant additive heritabilities for the traits analysed in the UK Biobank. Traits are ordered by $P$-values obtained through the block jack-knife. Very similar traits (for example Leg fat percentage (left), Leg fat percentage (right)) have been collapsed to trait with the most significant heritability.

| Phenotype | $h_D^2$ | SE $h_D^2$ | $P$ $h_D^2$ |
|---|---|---|---|
| Non-cancer illness code, self-reported: rheumatoid arthritis | 0.0669 | 0.0195 | 0.000307 |
| Diagnoses - main ICD10: K40 Inguinal hernia | 0.0322 | 0.00952 | 0.000363 |
| Alkaline phosphatase | 0.00936 | 0.00288 | 0.000567 |
| Treatment/medication code: co-codamol | 0.0405 | 0.0132 | 0.00108 |
| Treatment/medication code: thyroxine product | 0.0577 | 0.0202 | 0.00211 |
| Diagnoses - main ICD10: I84 Haemorrhoids | 0.025 | 0.00893 | 0.00255 |
| IGF-1 | 0.00513 | 0.00184 | 0.0027 |
| Added milk to filtered coffee | 0.0866 | 0.0319 | 0.00329 |
| Non-cancer illness code, self-reported: psoriasis | 0.0535 | 0.0206 | 0.00466 |
| Treatment/medication code: multivitamin + mineral preparations | 0.0469 | 0.0184 | 0.0054 |
| Gas or solid-fuel cooking/heating: A gas hob or gas cooker | 0.0067 | 0.00266 | 0.00591 |
| Traffic intensity on the nearest major road | 0.00388 | 0.00165 | 0.00947 |
| Hernia | 0.013 | 0.00555 | 0.00977 |
| Age at last live birth | 0.0116 | 0.00495 | 0.00981 |
| Worried most days during period of worst anxiety | 0.112 | 0.0487 | 0.0104 |
| Treatment/medication code: evening primrose oil | 0.0405 | 0.0176 | 0.0107 |
| Treatment/medication code: amlodipine | 0.0208 | 0.00905 | 0.0109 |
| Lipoprotein A | 0.0101 | 0.00441 | 0.0112 |
| Job SOC coding: Primary and nursery education teaching professionals | 0.0583 | 0.0256 | 0.0112 |
| Diagnoses - main ICD10: I21 Acute myocardial infarction | 0.0358 | 0.0157 | 0.0113 |
| Non-butter spread type details: Other type of spread/margarine | 0.039 | 0.0173 | 0.0122 |
| Major coronary heart disease event | 0.0265 | 0.0118 | 0.0124 |
| Which eye(s) affected by myopia (short sight): Both eyes | 0.132 | 0.0589 | 0.0126 |
| Detention categories: Informal, not formally detained | 0.0118 | 0.00532 | 0.0129 |
| Smoking status: Never | 0.00628 | 0.00282 | 0.013 |
| Fractured bone site(s): Ankle | 0.0414 | 0.0188 | 0.0137 |
| Types of physical activity in last 4 weeks: None of the above | 0.0153 | 0.007 | 0.0147 |
| Father still alive | 0.00761 | 0.0035 | 0.0149 |
| More irritable than usual during worst period of anxiety | 0.0699 | 0.0327 | 0.0162 |
| Tea intake | 0.0037 | 0.00174 | 0.017 |
| Job coding: civil service officer | 0.0698 | 0.0335 | 0.0186 |
| Been in serious accident believed to be life-threatening | 0.0291 | 0.014 | 0.0189 |
| Mixture of day and night shifts worked: Shift pattern for whole of job | 0.0814 | 0.0395 | 0.0195 |
| Aspartate aminotransferase | 0.00341 | 0.00167 | 0.0207 |
| Ever suffered mental distress preventing usual activities | 0.0173 | 0.00853 | 0.021 |
| Immature reticulocyte fraction | 0.00336 | 0.00167 | 0.0222 |
| Job SOC coding: Civil Service executive officers | 0.0672 | 0.0335 | 0.0225 |
| Bread type: Wholemeal or wholegrain | 0.00619 | 0.00311 | 0.0232 |
| Qualifications: NVQ or HND or HNC or equivalent | 0.00732 | 0.00375 | 0.0254 |
| Job coding: primary/nursery school teacher | 0.0513 | 0.0263 | 0.0254 |

**Table** S8: The top 40 most significant dominance heritabilities for the traits analysed in the UK Biobank. Traits are ordered by $P$-values obtained through the block jack-knife.

corporate a non-additive effect at each site which is uncorrelated from the additive contribution at that site. We then ask: what is the variance of this additional contribution? Letting $X^A$ denote the $n$ samples $\times$ $m$ sites matrix of genotypes (after rescaling to enforce a mean of 0 and a variance of 1), we define $X^D$ to be a re-coding of the genotypes whose columns $(X_j^D)$ are orthonormal to the columns of $X^A$ ($X_j^A$) under Hardy-Weinberg equilibrium. This extension to the additive infinitesimal model is:

$$y_i = \sum_{j=1}^{m} X_{i,j}^A \beta_{A_j} + \sum_{j=1}^{m} X_{i,j}^D \beta_{D_j} + \varepsilon_i, \tag{3}$$

where $y_i$ is a continuous phenotype measured in individual $i$, $\beta_{A_j}$ and $\beta_{A_j}$ are the additive and dominance effect sizes at site $j$ respectively, and $\varepsilon_i$ is a noise term. We mean center the genotypes and construct an orthonormal basis using the Gram-Schmidt procedure. The resultant additive and dominance encodings are

$$\frac{1}{\sqrt{2pq}} \begin{bmatrix} -2p \\ 1-2p \\ 2-2p \end{bmatrix} \text{ and } \begin{bmatrix} -p/q \\ 1 \\ -q/p \end{bmatrix}, \tag{4}$$

respectively. We wish to estimate $h_D^2 := \operatorname{Var}\left(\sum_{j=1}^{m} X_{i,j}^D \beta_{D_j}\right)$. By defining additive summary statistics as the marginal effect sizes obtained by regressing $y$ on $X_j^A$ as usual; $\widehat{\beta}_{A_j} = \frac{1}{n}\left(X_j^A\right)^\top y$, and introducing dominance summary statistics as the analogous marginal associations in the dominance encoding; $\widehat{\beta}_{D_j} = \frac{1}{n}\left(X_j^D\right)^\top y$, we may then proceed to derive a dominance LD-score equation relating dominance summary statistics to dominance LD-scores:

$$\mathbb{E}\left[\chi^2_{D_j}\right] = \frac{n h_D^2}{m} l_j^D + 1 \tag{5}$$

where $l_j^D = \sum_{j=1}^{m} (r_{j,k}^D)^2$ is the sum of the squared correlations between SNP $j$ and all other SNPs under the dominance encoding, and $\chi^2_{D_j}$ the chi-squared statistic for SNP $j$ under the dominance encoding, $\chi^2_{D_j} = n\beta^2_{D_j}$.

#### Fine-mapping

To fine-map putatively causal variants, we ran SuSIE (*22*) applied to summary statistics using the `susie_bhat` function in the SusieR package. Required inputs are the summary statistics in the locus under examination, together with the LD matrix in the locus. We are able to use SusieR straight out of the box by passing the in sample dominance LD matrix in the locus, and the dominance summary statistics from our dominance scan. To define loci for each phenotype, we merge the 1.5Mb neighbourhoods around genome-wide significant ($P < 5 \times 10^{-8}$) SNPs using pybedtools (*64, 65*). We exclude the HLA region, which we conservatively set as chr6 25-34Mb, from fine-mapping. We then pass each locus together with the UK Biobank .bgen file to LDstore (*66*) to obtain the additive LD matrix in the region. We then square this matrix entry-wise to estimate the dominance LD matrix at the locus. Finally, we pass the resultant dominance LD matrix and associated dominance summary statistics to SuSIE for each locus in turn for fine-mapping. We allowed up to ten causal variants per analysed region, and used a uniform prior for each variant being causal. To enable rapid parallel computation we make use of the dsub software (*67*). Code to prepare files for and run additive and dominance fine-mapping is available at `github.com/mkanai/xfinemap` (*68*).

#### Simulation studies

##### Fully simulated genotypes and phenotypes

To test the ability of LD score regression to recapitulate heritability parameters we run a series of simulation studies. Using the software msprime (*69*), we simulate collections of genotype data. Setting the number of sites to $10,000,000$ in $10$ independent chromosomes we sample $50,000$ individuals from the simulated European population and subset to variants with MAF $> 5\%$. This resulted in a collection of $\sim 250,000$ variants. We then fix this genotype data and simulate phenotypes from these variant sites using equation (3) with a given pair of additive and dominance heritabilities; $(h_A^2, h_D^2)$. We consider two genetic architectures: fully infinitesimal, and spike and slab with $10\%$ causal variants, and two classes of phenotypes: continuous and case-control. For case control phenotypes we use the liability threshold model to determine cases at $6\%$ prevalence in the population. Finally, we simulate the scenario in which the population prevalence of a case control phenotype is $6\%$, but an ascertained sample prevalence of $18\%$. After evaluating phenotypes, we then run marginal associations on a collection of additively encoded and dominance encoded SNPs. We assume that $20\%$ of sites have genotype information available, reflecting the scenario in which genotyping data is available at a subset of variable sites in the genome. We evaluate LD-scores at the subset of genotyped sites using $10\%$ of the simulated individuals. The LD-score summation is evaluated over all variant sites in the genome, mimicking the real world scenario in which a subset of population matched individuals have whole genome sequencing data available. Finally, we use LD-score regression to estimate the additive and dominance contributions to the heritability of the simulated phenotype. We set $h_A^2 = 0.2$, $h_D^2 \in \{0, 0.05, 0, 2\}$ and run $50$ independent phenotype simulations for each $(h_A^2, h_D^2)$ pair. The distribution of the resultant estimates are shown in Fig. S20-S26, and summarised in Table S3.

##### Real genotypes, simulated phenotypes

We also tested the performance of LD-score dominance by considering a subset of $10,000$, $50,000$, and $100,000$ samples from the Irish and British subset of the UK biobank data-set (*23*). We simulated effect sizes for $1,071,535$ Hapmap3 (HM3) variants (*63*) and used these to generate continuous phenotypes under the model, for all possible pairs of $h_A^2, h_D^2 \in \{0, 0.05, 0, 2\}$. Repeating $50$ independent replicates for each pair $(h_A^2, h_D^2)$. We then perform additive and dominance GWAS for each simulated phenotype using hail, and evaluate estimates of $h_A^2$ and $h_D^2$ using our software (*50*). Additive and dominance LD-scores were evaluated on HM3 variants in the $1000$ genomes European subset as described in (*70*). Results are displayed in Fig. S27-S28 and Summarised for sample size $50,000$ in Table S4.

#### Checking for artefactual dominance

##### Imputation error

We also examined the potential impact of imputation error on the non-additive signals that we observe. We take the info scores provided by the UK Biobank, and compare the distribution at dominance hits to that at sites that did not reach genome-wide significance for any of the phenotypes that we tested. The resultant distributions of info score are displayed in Fig. S5-S7.

##### Deviations from Hardy-Weinberg equilibrium

Another potential source of a dominance signal is through deviations from Hardy-Weinberg equilibrium (HWE). In our dominance re-coding (see section: LD-score dominance extension), we make that assumption that allele-frequencies are in HWE. If this is not the case, the two contributions are no longer uncorrelated, and so the additive signal can be pulled through into the dominance contribution.

We ran a simulation to test the impact of deviations from HWE on dominance $P$-values of association. We consider a single causal site and generate proportions of individuals in the population with each of the three genotype configurations under the following model:

$$P(AA) = (1-f)\,p^2 + pf$$
$$P(Aa) = 2pq(1-f)$$
$$P(aa) = (1-f)\,q^2 + qf$$

where $P$ is the MAF (frequency of A), $q = (1-p)$, and $f$ is the inbreeding coefficient. We then simulate continuous phenotypic data under the following model:

$$y = \beta X + \varepsilon \quad \varepsilon \sim \mathcal{N}\left(0, 1-\beta^2\right)$$

where $X$ is standardised. Thus, the effect of the SNP explains $\beta^2$ of the variance in $y$. We then incorrectly encode under the additive and dominance encodings and estimate effect sizes, as well as evaluating the $P$-value for the SNP not being in HWE using a $\chi^2$ goodness of fit test. Assuming a sample size of 50,000 and a SNP explaining $5\%$ of the phenotypic variance, we vary the MAF from 0.05 to 0.5, and inbreeding coefficient from 0 to 0.05 and display heat maps of the additive $P$-value for association, dominance $P$-value for association and HWE $P$-value in Figure S10.

To similarly test the impact of genotyping error on significance of dominance associations, we consider another split of the genotype configurations:

$$P(AA) = p^2 + 2pq\phi$$
$$P(Aa) = 2pq(1-\phi)$$
$$P(aa) = q^2$$

In this scenario, we assume that a proportion, $\phi$, of heterozygous calls we incorrectly called as homozygous variant. As before, we then incorrectly encode under the additive and dominance encodings and estimate effect sizes and $P$-value for the SNP not being in HWE using a $\chi^2$ goodness of fit test. Assuming a sample size of 50,000 and a SNP explaining $5\%$ of the phenotypic variance, we vary the MAF from 0.05 to 0.5, and $\phi$ from 0 to 0.05. We then display heat maps of the additive $P$-value for association, dominance $P$-value for association and HWE $P$-value in Fig. S9.

To determine if such a phenomenon is present in our results, we examine the distribution for $|\widehat{\beta}_D|$ in MAF and inbreeding coefficient ($f$) bins. If the effect is widespread, we expect to see a trend of increasing $|\widehat{\beta}_D|$ with increasing $|f|$. This trend was not present (Fig. S11 - S13).

##### Heteroskedastic noise

We run a Balding-Nichols model with two-populations, $1,000$ samples and $10,000$ independent markers. Using this simulated genotype data, we then simulate phenotypes under the additive model $y_i = \sum_{i=1}^{m} X_{i,j}\beta_j$, using ldscsim software (*71*). We simulate an initial $100\%$ heritable phenotype $\left(\beta_j \sim \mathcal{N}\left(0, \frac{1}{m}\right)\right)$ before transforming it by adding a noise term sampled from a normal distribution with a phenotype dependent variance. The variance of the noise term is set as:

$$\left(\frac{y_i - \tilde{y}}{-2\tilde{y}} 2(1 - h^2_{het})\right)\right)^{\alpha} \tag{6}$$

where $\tilde{y} = \min(\min(y), -\max(y))$ and $\alpha$ represents the desired heteroskedasticity of the noise term, which we set at $10$. We then run dominance GWAS on the resultant phenotype with heteroskedastic noise. Under this scenario no SNP has a non-additive effect, so any association is artefactual. We run this model under both an infinitesimal model and a spike and slab model: $\beta_j \sim \mathcal{N}\left(0, \frac{1}{\pi m}\right)$ with probability $\pi$, $\beta_j = 0$ with probability $(1-\pi)$ for each SNP $j$. We then inverse rank normalise the phenotype, and rerun GWAS. Results are shown in Fig. S30 and Fig. S31 for the infinitesimal, and spike and slab model setting $\pi = 0.01$, respectively.

### Supplementary Note

#### Additive LD score regression

We first consider the standard LD score regression in which we assume the infinitesimal model: the effect sizes of the genotypes $\beta_j;\ j = 1, 2, \ldots, m$ are independent with mean 0 and variance $\frac{h^2}{m}$. This is formulated as

$$y = X\beta + \varepsilon. \tag{7}$$

That is,

$$y_i = \sum_{j=1}^{m} X_{i,j}\beta_j + \varepsilon_i, \tag{8}$$

where $y$ is a vector of standardised phenotypes, and $X$ is an $n \times m$ matrix of $n$ standardised genotypes both taken from $m$ independent samples (unscaled genotypes $X'_{i,j} \in \{0, 1, 2\}$, subject to rescaling: $X_{i,j} = \left(X'_{i,j} - 2p_j\right) / \sqrt{2p_j(1-p_j)}$, where $p_j$ is the prevalence of the genotype at site $j$, in the population). Finally we assume that there is are independent uncorrelated noise terms which absorb the effect of the environment.

$$\varepsilon \sim \mathcal{N}(0, 1-h^2), \tag{9}$$

Note that $\mathrm{Var}(\varepsilon) = 1 - h^2$ since $\mathrm{Var}(X\beta) = h^2$ by definition of the narrow sense heritability $\left(\mathrm{Var}\left(\sum_{j=1}^{m} X_{i,j}\beta_j\right) = \mathrm{Var}(\beta_j)\,\mathrm{Var}\left(\sum_{j=1}^{m} X_{i,j}\right) = \frac{h^2}{m}\sum_{j=1}^{m}\mathrm{Var}(X_{i,j}) = h^2\right)$ and $\mathrm{Var}(y) = 1$ by construction. Under this model, all of the genetic heritability is additive.

We estimate effect sizes

$$\widehat{\beta}_j = \frac{1}{n}\sum_{i=1}^{n} X_{i,j} y_i = \frac{1}{n} X_j^\top y \tag{10}$$

where $X_j$ here are the rows of $X$. This is the marginal effect of SNP $j$ in the sample, and is simply the estimate of the effect size of SNP $j$ using a linear regression of genotype $X_j$ against the phenotype $y$ $\left(\widehat{\beta}_j = \frac{\sum_{i=1}^{n}\left(X_{i,j}-\overline{X_j}\right)(y_i-\overline{y})}{\sum_{i=1}^{n} X_{i,j}-\overline{X_j}} = \frac{\sum_{i=1}^{n} X_{i,j}y_i}{n}\right)$ Let us define $\chi_1^2$ coefficients for each $j$ as $\chi_j^2 = n\widehat{\beta}_j^2$. Note that $n\widehat{\beta}_j^2$ is $\chi_1^2$ distributed under the null due to the central limit theorem:

Recall that if $X_i$ are IID random variables with mean $\mu$ and variance $\sigma^2$, then

$$\sqrt{n}\left(\frac{1}{n}\sum_{i=1}^{n}(X_i - \mu)\right) \to \mathcal{N}(0, \sigma^2) \tag{11}$$

in distribution. We have $\mathbb{E}[X_{i,j}y_i] = 0$, $\mathrm{Var}(X_{i,j}y_i) = 1$. Thus $\sqrt{n}\left(\sum_{i=1}^{m} X_{i,j}y_i\right) \to \mathcal{N}(0,1)$, so $\widehat{\beta}_j \sim \frac{1}{\sqrt{n}}\mathcal{N}(0,1)$. Now recall that if $Y = \sum_{i=1}^{j} X_i^2$ where $X_i \sim \mathcal{N}(0,1)$, then $Y \sim \chi_j^2$. Thus, $n\beta_j^2 \sim \chi_1^2$ (a $\chi^2$ distribution with 1 degree of freedom).

Given our effect size estimate and our model in Equation (8), we can substitute into Equation (10):

$$\widehat{\beta}_j = \frac{1}{n}\sum_{i=1}^{n} X_{i,j}\left(\sum_{k=1}^{m} X_{i,k}\beta_k + \varepsilon_i\right) \tag{12}$$

$$= \sum_{k=1}^{m}\beta_k\left(\frac{1}{n}\sum_{i=1}^{n} X_{i,j}X_{i,k}\right) + \widetilde{\varepsilon}_j \tag{13}$$

$$= \sum_{k=1}^{m}\widehat{r}_{j,k}\beta_k + \widetilde{\varepsilon}_j \tag{14}$$

where $\widetilde{\varepsilon}_j = \frac{1}{n}\sum_{i=1}^{n} X_{i,j}\varepsilon_i$. We now determine the expectation of our $\chi^2$ coefficients to obtain the key formula.

$$
\begin{aligned}
\mathbb{E}[\chi_j^2] &= \mathbb{E}\left[n\widehat{\beta}_j^2\right] && (15)\\
&= \mathbb{E}\left[n\left(\sum_{k=1}^{m}\widehat{r}_{j,k}\beta_k + \widetilde{\varepsilon}_j\right)^2\right] && (16)\\
&= \mathbb{E}\left[n\sum_{k=1}^{m}\sum_{l=1}^{m}\widehat{r}_{j,k}\widehat{r}_{j,l}\beta_k\beta_l\right] + \underbrace{\mathbb{E}\left[2n\sum_{k=1}^{m}\widehat{r}_{j,k}\widetilde{\varepsilon}_j\right]}_{=0} + \mathbb{E}\left[n\widetilde{\varepsilon}_j^2\right] && (17)\\
&= \underbrace{n\mathbb{E}\left[\sum_{k\neq l}\widehat{r}_{j,k}\widehat{r}_{j,l}\beta_k\beta_l\right]}_{=0} + \mathbb{E}\left[n\sum_{k=1}^{m}\widehat{r}_{j,k}^2\beta_k^2\right] + n\mathbb{E}\left[\widetilde{\varepsilon}_j^2\right] && (18)\\
&= n\left(\sum_{k=1}^{m}\mathbb{E}\left[\widehat{r}_{j,k}^2\right]\mathbb{E}\left[\beta_k^2\right]\right) + n\mathbb{E}\left[\widetilde{\varepsilon}_j^2\right] && (19)
\end{aligned}
$$

by independence and linearity of expectations. Since $\beta_k$ is standardised, $\mathrm{Var}\left(\beta_k\right) = \mathbb{E}[\beta_k^2] = \frac{h^2}{m}$. Also, in an unstructured sample, we can make the approximation $\mathbb{E}\left[\widehat{r}_{j,k}^2\right] \approx r_{j,k}^2 + \frac{1}{n}$. This approximation comes from noting that $\widehat{r}$ is unbiased as we know the population variances for $X_j;\ j \in \{1, 2, \ldots, m\}$ are 1. Therefore, setting $X = X_j$, $Y = X_k$ to make notation simpler and using subscripts $i$ and $l$ to reiterate that summation is over samples,

$$
\begin{aligned}
\mathbb{E}\left[\widehat{r}_{X,Y}^2\right] &= \mathbb{E}\left[\left(\frac{1}{n}\sum_{i=1}^{n} X_iY_i\right)^2\right] && (20)\\
&= \frac{1}{n^2}\mathbb{E}\left[\sum_{l=1}^{n}\sum_{i=1}^{n} X_iY_iX_lY_l\right] && (21)\\
&= \frac{1}{n^2}\sum_{i,l:\, i\neq l}^{n}\mathbb{E}\left[X_iY_i\right]\mathbb{E}\left[X_lY_l\right] + \frac{1}{n^2}\sum_{i=1}^{n}\mathbb{E}\left[X_i^2Y_i^2\right] && (22)\\
&= \frac{1}{n^2}\left(n^2 - n\right) r_{X,Y}^2 + \frac{1}{n^2}\sum_{i=1}^{n}\mathbb{E}\left[X_i^2Y_i^2\right]. && (23)
\end{aligned}
$$

since $\mathbb{E}\left[X_iY_i\right] = r_{X,Y}$ and the genotypes of individuals $i$ and $l$ are independent in an unstructured population. Now, the majority of $X$ and $Y$ will be far apart and so approximately independent. Therefore,

$$\mathbb{E}\left[\widehat{r}^2_{X,Y}\right] \approx \left(1 - \frac{1}{n}\right) r^2_{X,Y} + \frac{1}{n^2} \sum_{i=1}^{n} \mathbb{E}\left[X_i^2\right] \mathbb{E}\left[Y_i^2\right] = \left(1 - \frac{1}{n}\right) r^2_{X,Y} + \frac{1}{n} \tag{24}$$

$$\approx r^2_{X,Y} + \frac{1}{n}. \tag{25}$$

Substituting this approximation into Equation (19), we have

$$\mathbb{E}\left[\chi_j^2\right] \approx \frac{nh^2}{m} \sum_{k=1}^{m} \left(r^2_{j,k} + \frac{1}{n}\right) + \mathbb{E}\left[\widetilde{\varepsilon}_j^2\right]. \tag{26}$$

Since $X_{i,j}$ and $\varepsilon_i$ are independent, the $X_{i,j}$ are standardised, and $\mathbb{E}[\widetilde{\varepsilon}_j] = \mathbb{E}[\varepsilon_i] = 0$,

$$\mathbb{E}\left[\widetilde{\varepsilon}_j^2\right] = \mathrm{Var}\left(\widetilde{\varepsilon}_j\right) = \mathrm{Var}\left(\frac{1}{n} \sum_{i=1}^{n} X_{i,j} \varepsilon_i\right) \tag{27}$$

$$= \frac{1}{n^2} \sum_{i=1}^{n} \mathrm{Var}\left(X_{i,j}\right) \mathrm{Var}\left(\varepsilon_i\right) \tag{28}$$

$$= \frac{1}{n^2} \sum_{i=1}^{n} \mathrm{Var}\left(\varepsilon_i\right) \tag{29}$$

$$= \frac{1}{n}\left(1 - h^2\right). \tag{30}$$

Thus,

$$\mathbb{E}\left[\chi_j^2\right] \approx \frac{nh^2}{m} \sum_{k=1}^{m} r^2_{j,k} + h^2 + 1 - h^2. \tag{31}$$

Finally, letting $l_j := \sum_{k=1}^{m} r^2_{j,k}$; dubbed the LD score of SNP $j$, we obtain the LD score regression formula:

$$\mathbb{E}\left[\chi_j^2\right] = \frac{nh^2}{m} l_j + 1. \tag{32}$$

#### LD score regression with dominance

For a more general model, we now allow a dominance term to affect the phenotype $y$.

$$y = X\beta + \varepsilon \tag{33}$$

$$= X^A \beta_A + X^D \beta_D + \varepsilon \tag{34}$$

$$y_i = \sum_{j=1}^{m} X_{i,j}^A \beta_{Aj} + \sum_{j=1}^{m} X_{i,j}^D \beta_{Dj} + \varepsilon_i \tag{35}$$

where now $X$ is the concatenation of two matrices; $X^A$ and $X^D$ (similarly, $\beta = (\beta_A, \beta_D)$). $X^A$ encodes the genotype (in exactly the same was as $X$ does in Equation (8)), and $X^D$ encodes a dominance contribution to the phenotype $y$. We will pick an zero mean-centered orthonormal encoding for $X^A$ and $X^D$. This encoding isolates the pure dominance and additive contributions of the genotypes. We start with the basis:

$$X^{0\prime} = \begin{bmatrix} 1 \\ 1 \\ 1 \end{bmatrix}, \; X^{A\prime} = \begin{bmatrix} 0 \\ 1 \\ 2 \end{bmatrix}, \; X^{D\prime} = \begin{bmatrix} 0 \\ 1 \\ 0 \end{bmatrix}, \tag{36}$$

where the first, second and third entries are the contributions of homozygous reference, heterozygous, and homozygous variant respectively. We then apply the Gram-Schmidt process (a method to orthonormalise a set of vectors in an inner product space), where our inner product is defined as

$$\langle X, Y \rangle = q^2 X_0 Y_0 + 2pq X_1 Y_1 + p^2 X_2 Y_2. \tag{37}$$

The subscript here denotes the genotype of the entry: (hom ref, het, hom var) for (0,1,2). We use this inner product as we wish to weight the contributions of the genotypes according to their prevalence in the population, assuming Hardy-Weinberg equilibrium. The inclusion of $[1, 1, 1]^\top$ will ensure that our resultant orthonormal basis has mean 0. Proceeding with Gram-Schmidt:

$$X^0 = \frac{1}{\|X^{0\prime}\|} \begin{bmatrix} 1 \\ 1 \\ 1 \end{bmatrix} = \begin{bmatrix} 1 \\ 1 \\ 1 \end{bmatrix}, \tag{38}$$

$$X^A = \frac{X^{A'} - \langle X^{A'}, X^0 \rangle X^0}{\left\| X^{A'} - \langle X^{A'}, X^0 \rangle X^0 \right\|} = \frac{1}{\sqrt{2pq}} \left( \begin{bmatrix} 0 \\ 1 \\ 2 \end{bmatrix} - \left(2pq + 2p^2\right) \begin{bmatrix} 1 \\ 1 \\ 1 \end{bmatrix} \right) \tag{39}$$

$$= \frac{1}{\sqrt{2pq}} \begin{bmatrix} -2p \\ 1-2p \\ 2-2p \end{bmatrix} \tag{40}$$

$$= \frac{1}{\sqrt{2pq}} \begin{bmatrix} -2p \\ q-p \\ 2q \end{bmatrix}, \tag{41}$$

$$X^D = \frac{X^{D'} - \langle X^{D'}, X^0 \rangle X^0 - \langle X^{D'}, X^A \rangle X^A}{\left\| X^{D'} - \langle X^{D'}, X^0 \rangle X^0 - \langle X^{D'}, X^A \rangle X^A \right\|} = \frac{\begin{bmatrix} 0 \\ 1 \\ 0 \end{bmatrix} - 2pq \begin{bmatrix} 1 \\ 1 \\ 1 \end{bmatrix} - (q-p) \begin{bmatrix} -2p \\ q-p \\ 2q \end{bmatrix}}{\left\| X^{D'} - \langle X^{D'}, X^0 \rangle X^0 - \langle X^{D'}, X^A \rangle X^A \right\|} \tag{42}$$

$$= \frac{\begin{bmatrix} -2p^2 \\ 2pq \\ -2q^2 \end{bmatrix}}{\left\| X^{D'} - \langle X^{D'}, X^0 \rangle X^0 - \langle X^{D'}, X^A \rangle X^A \right\|} \tag{43}$$

$$= \frac{1}{\sqrt{q^2p^4 + 2p^3q^3 + p^2q^4}} \begin{bmatrix} -p^2 \\ pq \\ -q^2 \end{bmatrix} \tag{44}$$

$$= \frac{1}{pq} \begin{bmatrix} -p^2 \\ pq \\ -q^2 \end{bmatrix}. \tag{45}$$

Each site has additive and dominance effect sizes ($\beta_A$ and $\beta_D$) which act along $X^A$ and $X^D$ respectively. $\beta_A$ and $\beta_D$ are drawn from distributions with mean 0 and variances $\frac{h_A^2}{m}$, $\frac{h_D^2}{m}$ respectively. We assume $\beta_j = (\beta_{Aj}, \beta_{Dj})$ are independent across sites, but may have dependencies at a given locus. Let's determine the variance of $X_i\beta$ ($= X_i^A\beta_A + X_i^D\beta_D$), where here, $X_i$ denotes the $i^{\text{th}}$ row of the matrix $X$.

$$\text{Var}\left(X_i\beta\right) = \text{Var}\left(\sum_{j=1}^{m} X_{i,j}\beta_j\right) \tag{46}$$

$$= \mathbb{E}\left[\sum_{j=1}^{m}\sum_{l=1}^{m} X_{i,j}X_{i,l}\beta_j\beta_l\right] \tag{47}$$

$$= \mathbb{E}\left[\sum_{j=1}^{m}\sum_{l=1}^{m} X_{i,j}^{A}X_{i,l}^{A}\beta_{Aj}\beta_{Al} + X_{i,j}^{A}X_{i,l}^{D}\beta_{Aj}\beta_{Dl} + X_{i,j}^{D}X_{i,l}^{A}\beta_{Dj}\beta_{Al} + X_{i,j}^{D}X_{i,l}^{D}\beta_{Dj}\beta_{Dl}\right]. \tag{48}$$

We may remove all cross terms over sites as we assume that $\beta_{Aj}$, $\beta_{Dj}$ are independent across sites, and also independent of the genetic data encoded in $X^A$ and $X^D$. Thus,

$$\mathbb{E}\left[X_i\beta\right] = \mathbb{E}\left[\sum_{j=1}^{m} {X_{i,j}^{A}}^2\beta_{Aj}^2 + X_{i,j}^{A}X_{i,j}^{D}\beta_{Aj}\beta_{Dj} + X_{i,j}^{D}X_{i,j}^{A}\beta_{Dj}\beta_{Aj} + {X_{i,j}^{D}}^2\beta_{Dj}^2\right]. \tag{49}$$

Now we may use that $\mathbb{E}\left[X_{i,j}^{A}X_{i,j}^{D}\right] = \mathbb{E}\left[X_{i,j}^{A}\right]\mathbb{E}\left[X_{i,j}^{D}\right] = 0$, as $X^A$ and $X^D$ are orthogonal by construction. Thus, we are left with

$$\text{Var}\left(X_i\beta\right) = \mathbb{E}\left[\sum_{j=1}^{m} {X_{i,j}^{A}}^2\beta_{Aj}^2\right] + \mathbb{E}\left[\sum_{j=1}^{m} {X_{i,j}^{D}}^2\beta_{Dj}^2\right] \tag{50}$$

$$= \sum_{j=1}^{m}\mathbb{E}\left[{X_{i,j}^{A}}^2\right]\mathbb{E}\left[\beta_{Aj}^2\right] + \sum_{j=1}^{m}\mathbb{E}\left[{X_{i,j}^{D}}^2\right]\mathbb{E}\left[\beta_{Dj}^2\right] \tag{51}$$

$$= \frac{h_A^2}{m}\sum_{j=1}^{m} 1 + \frac{h_D^2}{m}\sum_{j=1}^{m} 1 = h_A^2 + h_D^2. \tag{52}$$

So $\text{Var}\left(X\beta\right) = h_A^2 + h_D^2 = h^2$. As in the standard infinitesimal model, the noise term $\varepsilon$ has mean 0 and variance $1 - h^2$. $h_A^2 \leq h_A^2 + h_D^2 = h^2$. In particular, equalities hold when the dominance terms don't contribute to the variance.

We wish to determine the extent to which $h^2 > h_A^2$. To achieve this we can simply compare the sums of the first order terms of two LD score regression models. The first under the standard infinitesimal model as outlined above, and the second according the model introduced in Equation (33). It remains to derive the analogue of Equation (32), which can be obtained in a similar manner to the original LD score regression model.

Again, we estimate effect sizes but this time regress our dominance encoded SNPs against the phenotypes as well:

$$\widehat{\beta}_{A_j} = \frac{1}{n}\sum_{i=1}^{n} X_{i,j}^{A} y_i = \frac{1}{n} X_j^{A\top} y; \qquad \widehat{\beta}_{D_j} = \frac{1}{n}\sum_{i=1}^{n} X_{i,j}^{D} y_i = \frac{1}{n} X_j^{D\top} y. \tag{53}$$

This time, we substitute $y_i$ using Equation (33):

$$\widehat{\beta}_{A_j} = \frac{1}{n}\sum_{i=1}^{n} X_{i,j}^{A} \left( \sum_{k=1}^{m} X_{i,k}^{A}\beta_{A_k} + \sum_{k=1}^{m} X_{i,k}^{D}\beta_{D_k} + \varepsilon_i \right) \tag{54}$$

$$= \sum_{k=1}^{m} \beta_{A_k} \frac{1}{n}\sum_{i=1}^{n} X_{i,j}^{A} X_{i,k}^{A} + \sum_{k=1}^{m} \beta_{D_k} \frac{1}{n}\sum_{i=1}^{n} X_{i,j}^{A} X_{i,k}^{D} + \frac{1}{n}\sum_{i=1}^{n} X_{i,j}^{A}\varepsilon_i \tag{55}$$

$$= \sum_{k=1}^{m} \widehat{r}_{j,k}^{AA}\beta_{A_k} + \sum_{k=1}^{m} \widehat{r}_{j,k}^{AD}\beta_{D_k} + \frac{1}{n}\sum_{i=1}^{n} X_{i,j}^{A}\varepsilon_i. \tag{56}$$

Similarly,

$$\widehat{\beta}_{D_j} = \sum_{k=1}^{m} \widehat{r}_{j,k}^{DA}\beta_{A_k} + \sum_{k=1}^{m} \widehat{r}_{j,k}^{DD}\beta_{D_k} + \frac{1}{n}\sum_{i=1}^{n} X_{i,j}^{D}\varepsilon_k. \tag{57}$$

Setting $\widetilde{\varepsilon}_j = \frac{1}{n}\sum_{i=1}^{n} X_{i,j}^{A}\varepsilon_i$, we now determine the expectation of the $\chi^2$ statistic,

$$\chi_{A_j}^2 := n\widehat{\beta}_{A_j}. \tag{58}$$

(Obtaining the expectation of $\chi_{D_j}^2 := n\widehat{\beta}_{D_j}$ is analogous).

$$\mathbb{E}\left[\chi_{A_j}^2\right] = \mathbb{E}\left[n\beta_{A_j}^2\right] = n\mathbb{E}\left[\left( \sum_{k=1}^{m} \widehat{r}_{j,k}^{AA}\beta_{A_k} + \sum_{k=1}^{m} \widehat{r}_{j,k}^{AD}\beta_{D_k} + \widetilde{\varepsilon}_j \right)^2\right] \tag{59}$$

$$= n\mathbb{E}\left[\sum_{k=1}^{m}\sum_{l=1}^{m} \left( \widehat{r}_{j,k}^{AA}\beta_{A_k}\widehat{r}_{l,k}^{AA}\beta_{A_l} + \widehat{r}_{j,k}^{AA}\beta_{A_k}\widehat{r}_{l,k}^{AD}\beta_{D_l} + \right.\right. \tag{60}$$

$$\left.\left. \widehat{r}_{j,k}^{AD}\beta_{D_k}\widehat{r}_{l,k}^{AA}\beta_{A_l} + \widehat{r}_{j,k}^{AD}\beta_{D_k}\widehat{r}_{l,k}^{AD}\beta_{D_l} \right) + \underbrace{\widetilde{\varepsilon}_j \left( \sum_{k=1}^{m} \widehat{r}_{j,k}^{AA}\beta_{A_k} + \widehat{r}_{j,k}^{AD}\beta_{D_k} \right)}_{=0} + \widetilde{\varepsilon}_j^2 \right]$$

Again, we may remove all cross terms over sites due to the independence of the $\beta$s.

$$= n\mathbb{E}\left[\sum_{k=1}^{m} \widehat{r}_{j,k}^{AA^2} \beta_{A_k}^2\right] + 2n\mathbb{E}\left[\sum_{k=1}^{m} \widehat{r}_{j,k}^{AA} \beta_{A_k} \widehat{r}_{j,k}^{AD} \beta_{D_k}\right] + \tag{61}$$

$$n\mathbb{E}\left[\sum_{k=1}^{m} \widehat{r}_{j,k}^{AD^2} \beta_{D_k}^2\right] + n\mathbb{E}\left[\widehat{\varepsilon}_j^2\right]$$

We may replace the $\widehat{r}_{j,k}^{AA^2}$ and $\widehat{r}_{j,k}^{AD^2}$ as before, but we also have a collection of cross terms (i.e. $\widehat{r}_{j,k}^{AA} \beta_{A_k} \widehat{r}_{j,k}^{AD} \beta_{D_k}$) to consider. Let's determine the expectation of this summation of cross terms.

$$\mathbb{E}\left[\sum_{k=1}^{m} \beta_{A_k} \beta_{D_k} \widehat{r}_{j,k}^{AA} \widehat{r}_{j,k}^{AD}\right] = \sum_{k=1}^{m} \mathbb{E}\left[\beta_{A_k} \beta_{D_k}\right] \mathbb{E}\left[\widehat{r}_{j,k}^{AA} \widehat{r}_{j,k}^{AD}\right]. \tag{62}$$

There may be dependency between the additive a dominance effect sizes a locus. So let's turn our attention to $\mathbb{E}\left[\widehat{r}_{j,k}^{AA} \widehat{r}_{j,k}^{AD}\right]$:

$$\mathbb{E}\left[\widehat{r}_{j,k}^{AA} \widehat{r}_{j,k}^{AD}\right] = \mathbb{E}\left[\frac{1}{n^2} \sum_{i=1}^{n} \sum_{l=1}^{n} X_{i,j}^A X_{i,k}^A X_{l,j}^A X_{l,k}^D\right] \tag{63}$$

$$= \frac{1}{n^2} \sum_{i,l:\, i \neq l} \mathbb{E}\left[X_{i,j}^A X_{i,k}^A X_{l,j}^A X_{l,k}^D\right] + \frac{1}{n^2} \sum_{i=1}^{n} \mathbb{E}\left[X_{i,j}^{A^2} X_{i,k}^A X_{i,k}^D\right] \tag{64}$$

$$= \frac{1}{n^2} \left(n^2 - n\right) r_{j,k}^{AA} r_{j,k}^{AD} + \frac{1}{n^2} \sum_{i=1}^{n} \mathbb{E}\left[X_{i,j}^{A^2} X_{i,k}^A X_{i,k}^D\right]. \tag{65}$$

Since, $j$ and $k$ are far apart for most SNPs, we make the approximation

$$\mathbb{E}\left[\widehat{r}_{j,k}^{AA} \widehat{r}_{j,k}^{AD}\right] \approx \left(1 - \frac{1}{n}\right) r_{j,k}^{AA} r_{j,k}^{AD} + \sum_{i=1}^{n} \mathbb{E}\left[X_{i,j}^{A^2}\right] \mathbb{E}\left[X_{i,k}^A X_{i,k}^D\right]. \tag{66}$$

Note that $X_{i,j}^D$ and $X_{i,j}^A$ are independent for all $j$ by construction. So,

$$\mathbb{E}\left[\widehat{r}_{j,k}^{AA} \widehat{r}_{j,k}^{AD}\right] \approx r_{j,k}^{AA} r_{j,k}^{AD}. \tag{67}$$

Plugging in this approximation, and the approximation in Equation (25) into Equation (61), we

| $j/k$ | 0 | 1 |
|---|---|---|
| 0 | $1-p_j-p_k$ | $p_j-p_{j,k}$ |
| 1 | $p_k-p_{j,k}$ | $p_{j,k}$ |

**Table** S9:

have

$$\mathbb{E}\left[\chi^2_{A_j}\right] \approx n\sum_{k=1}^{m}\underbrace{\left(r_{j,k}^{AA^2}+\frac{1}{n}\right)}_{\text{by Equation (25)}}\mathbb{E}\left[\beta^2_{A_k}\right]+n\sum_{k=1}^{m}\underbrace{\left(r_{j,k}^{AD^2}+\frac{1}{n}\right)}_{\text{by Equation (25)}}\mathbb{E}\left[\beta^2_{D_k}\right]+ \tag{68}$$

$$\sum_{k=1}^{m}\mathbb{E}\left[\beta_{A_k}\beta_{D_k}\right]\underbrace{r_{j,k}^{AA}r_{j,k}^{AD}}_{\text{by Equation (67)}}+n\mathbb{E}\left[\widetilde{\varepsilon}_j^2\right]$$

$$=\frac{nh_A^2}{m}\sum_{k=1}^{m}r_{j,k}^{AA^2}+\frac{nh_D^2}{m}\sum_{k=1}^{m}r_{j,k}^{AD^2}+h_A^2+h_D^2+\sum_{k=1}^{m}\mathbb{E}\left[\beta_{A_k}\beta_{D_k}\right]r_{j,k}^{AA}r_{j,k}^{AD}+\frac{n(1-h^2)}{n} \tag{69}$$

$$=\frac{nh_A^2}{m}\sum_{k=1}^{m}r_{j,k}^{AA^2}+\frac{nh_D^2}{m}\sum_{k=1}^{m}r_{j,k}^{AD^2}+\sum_{k=1}^{m}\mathbb{E}\left[\beta_{A_k}\beta_{D_k}\right]r_{j,k}^{AA}r_{j,k}^{AD}+1. \tag{70}$$

Similarly,

$$\mathbb{E}\left[\chi^2_{D_j}\right]=\frac{nh_A^2}{m}\sum_{k=1}^{m}r_{j,k}^{DA^2}+\frac{nh_D^2}{m}\sum_{k=1}^{m}r_{j,k}^{DD^2}+\sum_{k=1}^{m}\mathbb{E}\left[\beta_{A_k}\beta_{D_k}\right]r_{j,k}^{DD}r_{j,k}^{AD}+1. \tag{71}$$

We can investigate the contribution of each of the correlation terms analytically. Let $p_j$ and $p_k$ denote the probability of observing the alternate genotype at sites $j$ and $k$ respectively, and $p_{j,k}$ denote the probability of observing the alternate genotype at both sites $j$ and $k$.

This then leads to the probability mass function of linked genotypes in offspring (in the absence of inbreeding) by multiplying the relevant entries of Table S9. We may then determine the correlation coefficients ($r^{AA}$, $r^{AD}$, $r^{DA}$, $r^{DD}$):

$$r^{QR}=\mathbb{E}\left[X_Q^\top X_R\right]=\sum_{g_j,g_k\in\{0,1,2\}}P(g_j,g_k)X_j^Q(g_j)X_k^R(g_k);\quad Q,R\in\{A,D\}. \tag{72}$$

where

$$X_j^A = \frac{1}{\sqrt{2p_jq_j}}\begin{bmatrix} -2p_j \\ q_j - p_j \\ 2q_j \end{bmatrix}, \; X_j^D = \frac{1}{p_jq_j}\begin{bmatrix} -p_j^2 \\ p_jq_j \\ -q_j^2 \end{bmatrix}. \tag{73}$$

Here, using an abuse of notation, we mean the encoding of the genotypes (homozygous reference, heterozygous, homozygous variant) at site $j$. After cancelling, we obtain:

$$r^{AD} = 0, \quad r^{DA} = 0, \tag{74}$$

$$r^{AA} = \frac{p_{j,k} - p_jp_k}{\sqrt{p_jp_k\left(1-p_j\right)\left(1-p_k\right)}}, \quad r^{DD} = r^{AA^2} = \frac{\left(p_{j,k} - p_jp_k\right)^2}{p_jp_k\left(1-p_j\right)\left(1-p_k\right)}. \tag{75}$$

Equations (70) and (71) then simplify to

$$\mathbb{E}\left[\chi^2_{A_j}\right] = \frac{nh_A^2}{m} l_j^A + 1; \tag{76}$$

$$\mathbb{E}\left[\chi^2_{D_j}\right] = \frac{nh_D^2}{m} l_j^D + 1, \tag{77}$$

where $l_j^A = \sum_{j=1}^m r_{j,k}^{AA^2}$ and $l_j^D = \sum_{j=1}^m r_{j,k}^{DD^2}$ To check that $r^{DD} = r^{AA^2}$ is true empirically, we consider chromosome 22 in the 1000 genomes data (*55*) and plot the proportions of samples with each genotype combination, and contrast this to the analytic expectation under random mixing. We also compare the empirical squared additive and dominance genetic correlation as we move away from a lead SNP. We find that empirical estimates track closely with the analytic solution under random mixing of individuals, shown in Fig. S1.

Note that, as with the additive LD-score regression, we can redefine LD-scores to allow any allele frequency dependence. The only point in the derivation which requires an allele frequency effect size assumption is in the evaluation of $\mathbb{E}\left[\beta_{D,j}^2\right]$. For example, akin to Schoech *et al.* (*72*), we can define some parameter $\alpha_D$ to modulate the allele frequency dependence. To do so, we need only define a new flavour of dominance LD-score following a rescaling within the summation. We define:

$$l_{j,\alpha}^D = \sum_{k=1}^m r_{j,k}^{DD^2} \left(pq\right)^2 \alpha_D. \tag{78}$$

Equivalently, we may consider a re-coding of genotypes as in the MAMA manuscript (*73*) and achieve the same result. This is an underappreciated extension to LD-score regression which is already present in the software via the `--pq-exp` flag.

#### Similarity of logistic and linear regression with small effect sizes

Under the logistic model, we have:

$$p = \frac{1}{1 + \exp\left(-\left(\beta_0 + \beta_1 X\right)\right)}.$$

When effects are small, $\left|\exp\left(-\left(\beta_0 + \beta_1 X\right)\right)\right| < 1$, so

$$\frac{1}{1 + \exp\left(-\left(\beta_0 + \beta_1 X\right)\right)} = \sum_{n=0}^{\infty} \left(-\exp\left(-\left(\beta_0 + \beta_1 X\right)\right)\right)^n$$

Again, when effects are small,

$$\begin{aligned} \sum_{n=0}^{\infty} \left(-\exp\left(-\left(\beta_0 + \beta_1 X\right)\right)\right)^n &\approx 1 - \exp\left(-\left(\beta_0 + \beta_1 X\right)\right) \\ &\approx \beta_0 + \beta_1 X. \end{aligned}$$

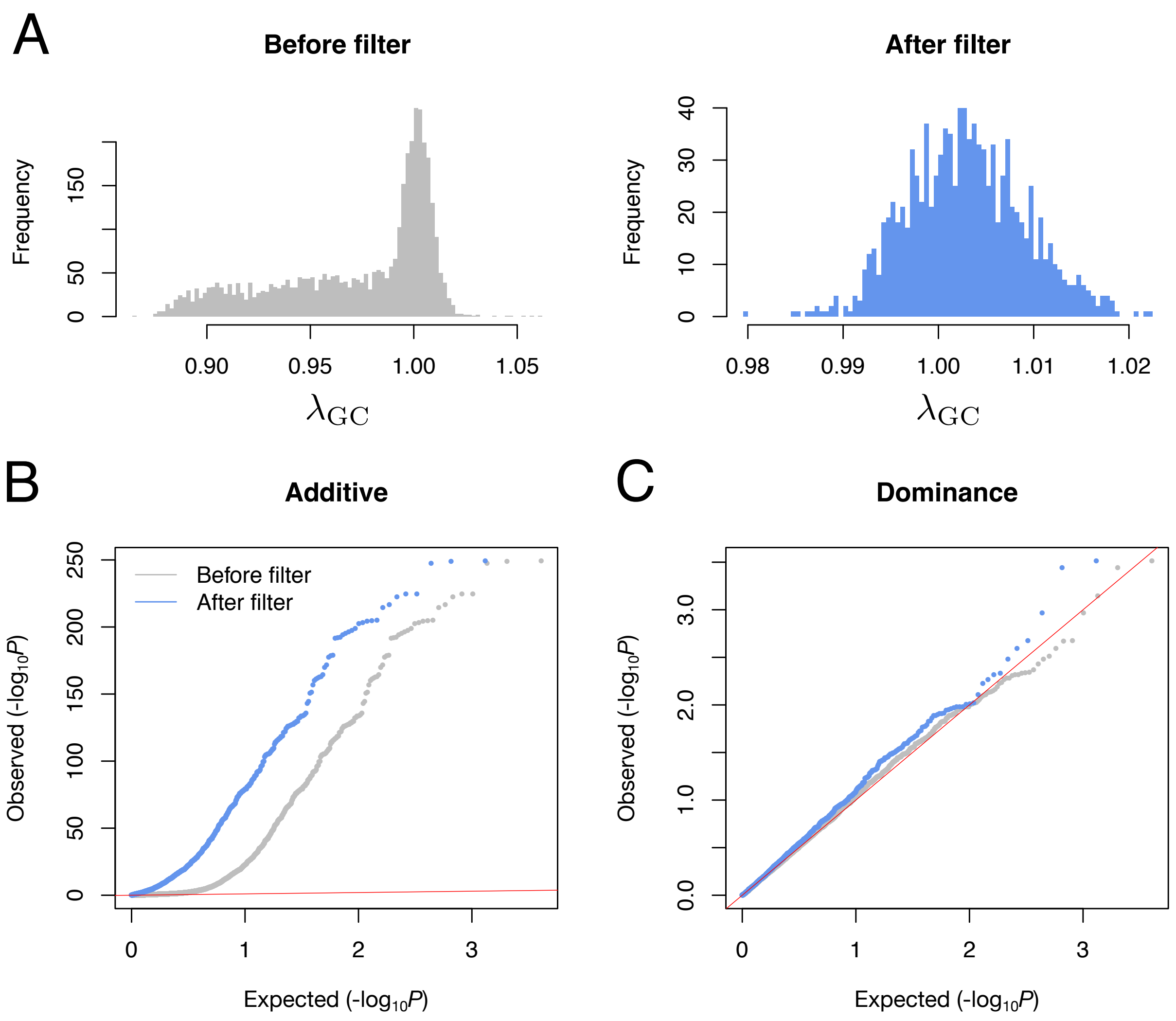


**Figure** S32: Filtering phenotypes. In A, following GWAS we examined $\lambda_{GC}$ of the HapMap SNPs used in heritability estimation. We observed significant deflation in marginal dominance effect sizes. Following removal of continuous phenotypes with at least 50,000 non-missing values and categorical phenotypes with at least 3,000 cases and controls this deflated tail was removed. In B, the QQ plots comparing the null uniform distribution of $P$-values to those observed, we see strong evidence for phenotypic variance explained by additive genetics, which increases with availability of non-missing data. By contrast, in C, the QQ plot comparing the uniform null distribution of $P$-values against those observed for non-additive genetic contribution does not display any significant inflation in the tail, before and after restriction to phenotypes with a large proportion of non-missing values. Throughout, points and bars in grey and blue are represent phenotypes before and after the restriction to phenotypes with large numbers of non-missing data (continous phenotypes with $> 50,000$ non-missing values and categorical phenotypes with $> 3,000$ cases and controls) respectively.

50. Dominance LD-score github repository, `https://github.com/astheeggeggs/ldscgxe`.

51. Neale lab heritability browser, https://nealelab.github.io/UKBB_ldsc/h2_browser.html.

52. W. Huang, T. F. C. Mackay, *PLoS Genet.* **12**, e1006421 (2016).

53. J. M. Alvarez-Castro, O. Carlborg, *Genetics* **176**, 1151 (2007).

54. Z. G. Vitezica, A. Legarra, M. A. Toro, L. Varona, *Genetics* **206**, 1297 (2017).

55. A. Auton, *et al.*, *Nature* **526**, 68 (2015).

56. Modified PHESANT github repository, https://github.com/astheeggeggs/PHESANT.

57. FinnGen endpoint definitions, https://www.finngen.fi/en/researchers/clinical-endpoints.

58. L. A. C. Millard, N. M. Davies, T. R. Gaunt, G. Davey Smith, K. Tilling, *International journal of epidemiology* **47**, 29 (2018).

59. W. McLaren, *et al.*, *Genome biology* **17**, 122 (2016).

60. hail 0.1 documentation, https://hail.is/docs/0.1/.

61. hail github repository, https://github.com/hail-is/hail.

62. hail dominance modified github repository, https://github.com/astheeggeggs/hail.

63. *Nature* **426**, 789 (2003).

64. R. K. Dale, B. S. Pedersen, A. R. Quinlan, *Bioinformatics (Oxford, England)* **27**, 3423 (2011).

65. A. R. Quinlan, I. M. Hall, *Bioinformatics (Oxford, England)* **26**, 841 (2010).

66. C. Benner, *et al.*, *American journal of human genetics* **101**, 539 (2017).

67. dsub github repository, https://github.com/googlegenomics/dsub.

68. xfinemap github repository, https://github.com/mkanai/xfinemap.

69. J. Kelleher, A. M. Etheridge, G. McVean, *PLoS computational biology* **12**, e1004842 (2016).

70. P. Turley, *et al.*, *Nature genetics* **50**, 229 (2018).

71. ldscsim github repository, https://github.com/nikbaya/ldscsim.

72. A. P. Schoech, *et al.*, *Nature communications* **10**, 790 (2019).

73. P. Turley, *et al.*, *bioRxiv* (2021).
